# Engulfment by brain macrophages in a short-lived vertebrate

**DOI:** 10.64898/2026.04.28.721434

**Authors:** Rahul Nagvekar, Angela N. Pogson, Prateek R. Kalakuntla, Helena J. Barr, Azalia M. Martínez Jaimes, S.V. Perry, Emma K. Costa, Jingxun Chen, Felix Boos, Paloma Navarro Negredo, Luise A. Seeker, James B. Jaggard, Rogelio Barajas, Philippe Mourrain, Param Priya Singh, Stephen R. Quake, Tony Wyss-Coray, Kristy Red-Horse, Beth Stevens, Bo Wang, Claire N. Bedbrook, Ravi D. Nath, Anne Brunet

## Abstract

Engulfment by macrophages is critical for waste clearance in the vertebrate brain. Understanding clearance mechanisms may open new therapeutic possibilities to counter brain aging and neurodegenerative diseases. However, few *in vivo* models exist to study engulfment in the brain and characterize this process during aging and across species. Here we present a genetic model for secretion of a fluorescent protein by neurons in the brain of the African turquoise killifish, the shortest-lived vertebrate that can be bred in captivity. We use this model to identify a population of brain macrophages in the killifish responsible for engulfment of material from the brain extracellular space. Intriguingly, many of these cells bear similarities to mammalian border-associated and monocyte-derived macrophages, rare subsets of macrophages in mouse and human brains noted for their engulfment capabilities. We also find that in our model, killifish brain macrophages decline in engulfment capacity with age. This work highlights how vertebrate brain macrophages, particularly those at brain border regions, can play a critical role in clearance and provides an opportunity to test interventions that can boost engulfment by these macrophages to promote brain resilience in old age and disease.

## Introduction

The vertebrate brain uses varied strategies to clear extracellular material such as extracellular matrix and aggregation-prone proteins^1–3^. Engulfment by specialized cells^4–7^ – including macrophages such as microglia^8–16^ – permits degradation and clearance of extracellular proteins and other macromolecules in the brain. Disruptions in clearance can lead to dysregulated accumulation of extracellular material in the brain interstitial space and cerebrospinal fluid (CSF), with detrimental consequences during aging and in neurodegenerative diseases^17–30^. Thus, clearance mechanisms could represent key strategies for maintaining brain homeostasis, with promising therapeutic potential to counter brain aging and neurodegenerative diseases^31–37^. Studying clearance strategies in varied species could reveal mechanisms not present in mammals and help determine how they could be used to counter brain aging.

An interesting model system in which to study clearance strategies is the African turquoise killifish *Nothobranchius furzeri* – the shortest-lived vertebrate that can be bred in captivity^38^. The killifish has a median lifespan of only six months, 5 times shorter than the mouse^38^, enabling scalable studies of processes that are disrupted with age^39–59^. The killifish also recapitulates many conserved hallmarks of brain aging, including neurodegeneration^60–62^, cognitive decline^63–65^, and behavioral deficits^66^. However, waste clearance in the brain has not been examined in this species. A short-lived vertebrate model in which to study clearance would enable longitudinal studies as well as the identification of interventions that boost clearance in the aged brain.

Here, we present a genetic model for studying clearance of a secreted fluorescent protein in the brain of the killifish. Using this genetic model, we show that a population of macrophages in the killifish brain is responsible for engulfment of diverse extracellular molecules. Intriguingly, we find that many killifish brain macrophages exhibit similarities to mammalian border-associated macrophages (BAMs) and monocyte-derived macrophages (MDMs), which are rare cell types in the adult mammalian brain. We also find that in our model, the engulfment capacity of killifish brain macrophages declines with age, which could contribute to clearance dysregulation and abnormal accumulation of extracellular material in brain aging.

### A model for engulfment by vertebrate brain myeloid cells

To model clearance of extracellular proteins in the brain, we engineered a killifish strain using a CRISPR knockin method developed for killifish^67^. In this strain, a secreted form of the red fluorescent protein oScarlet is expressed from one allele of the *ELAVL3* locus, a vertebrate-conserved gene predominantly expressed in neurons^68^ (**Fig. 1a**). We chose oScarlet for visualizing clearance processes due the relatively high stability of mScarlet-derived red fluorescent proteins^69,70^. Importantly, to allow the secretion of oScarlet into the interstitial space, we added a signal peptide (SP, derived from the killifish HSPA5 protein^71^) to the N-terminal region of oScarlet. We also added a P2A self-cleaving peptide to prevent fusion of oScarlet to the ELAVL3 protein^72^ (**Fig. 1a**). We maintained the killifish line at the heterozygous state and refer to it as SP-oScarlet.

**Figure 1.**
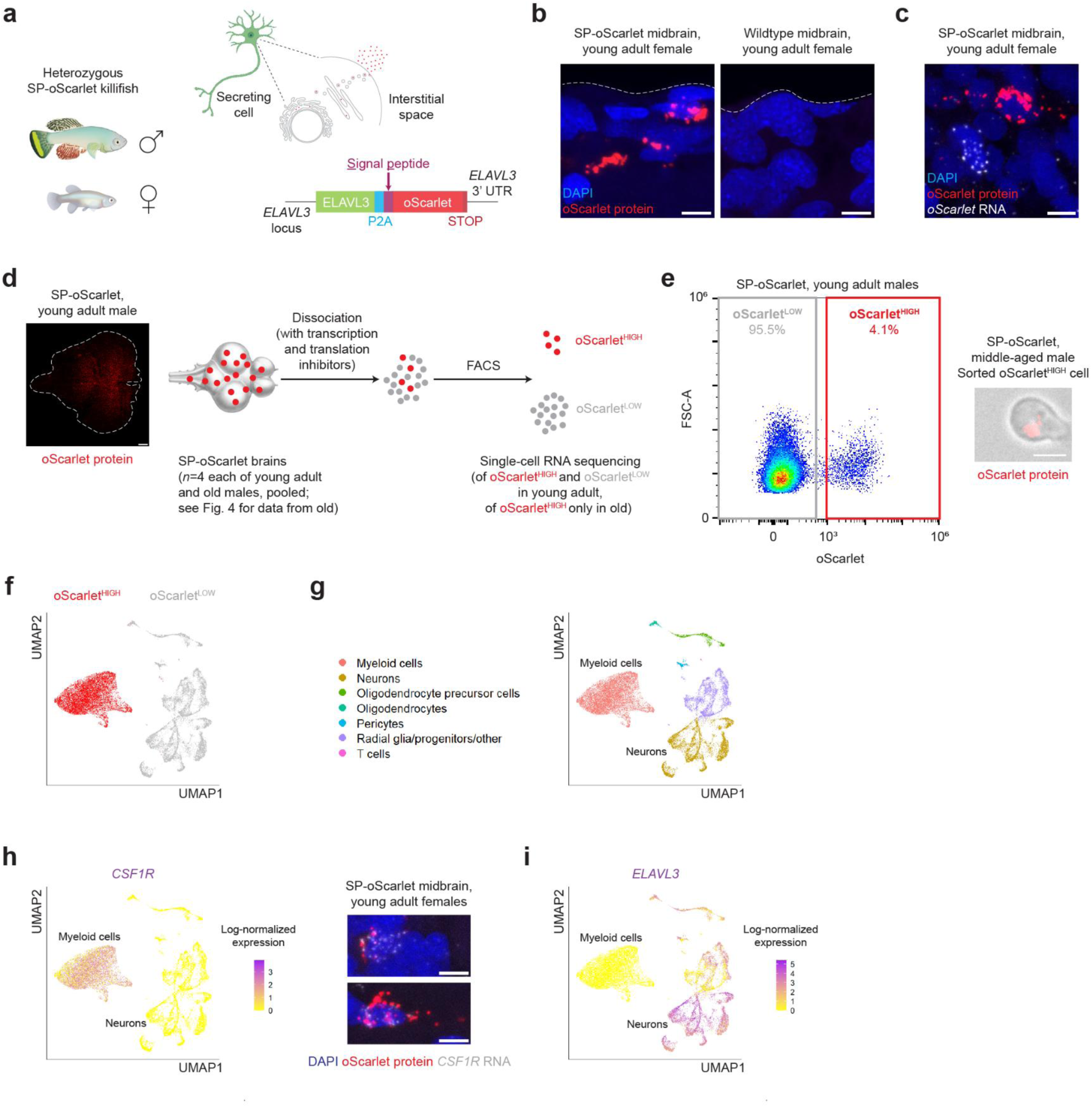
A model for engulfment by vertebrate brain myeloid cells. **a,** Schematic for the SP-oScarlet killifish line, which features expression and secretion of a red fluorescent protein, oScarlet, from non-immune cells of the killifish brain using CRISPR knockin at the *ELAVL3* locus. P2A, self-cleaving peptide; UTR, untranslated region. The SP-oScarlet line is maintained in the heterozygous state. Art from BioRender (cell) and by Rogelio Barajas (killifish). **b,** Representative images of brain sections from young adult (59 days) female heterozygous SP-oScarlet and young adult (59 days) female wildtype killifish, highlighting oScarlet protein signal (native fluorescence plus anti-mCherry immunofluorescence staining) in the SP-oScarlet brain (representative of n = 13 SP-oScarlet fish over three experiments and n = 3 wildtype fish in one experiment). Scale bars = 5 μm. **c,** Representative image of a young adult (59 days) female heterozygous SP-oScarlet killifish brain section, highlighting oScarlet protein signal (native fluorescence plus anti-mCherry immunofluorescence staining) in a cell that does not express *oScarlet* RNA (representative of n = 10 fish, 59-63 days females, over two experiments). Scale bar = 5 μm. **d,** (Left) 3D image of a cleared (modified Adipo-Clear method) young adult (50 days) male heterozygous SP-oScarlet killifish midbrain/hindbrain, highlighting punctate oScarlet protein signal (native fluorescence plus anti-RFP immunofluorescence staining) (n = 1 fish). (Right) Schematic for single-cell RNA sequencing of cells with and without oScarlet protein signal in young adult (49 days) and old (134 days) male heterozygous SP-oScarlet killifish brains (n = 4 fish, pooled, per age group, in one experiment). For data from old fish and for an independent experiment in old fish, see Fig. 4. **e,** (Left) Fluorescence-activated cell sorting (FACS) plot for young adult pooled brains from experiment in **d**, highlighting oScarlet^LOW^ and oScarlet^HIGH^ cells. Each dot represents one cell. (Right) Image of an oScarlet^HIGH^ cell isolated from a middle-aged (76 days) male heterozygous SP-oScarlet killifish brain by FACS, highlighting oScarlet protein signal (native fluorescence only) (n = 1 fish). Scale bar = 2 μm. **f,** Uniform Manifold Approximation and Projection (UMAP) of 11,503 oScarlet^LOW^ and 9,680 oScarlet^HIGH^ high-quality young adult cells from experiment in **d**. Each dot represents one cell. **g**, UMAP as in **f** colored according to manually annotated cell types, labeled in legend. Each dot represents one cell. **h,** (Left) UMAP as in **f** colored according to log-normalized *CSF1R* (*LOC107387189*) expression. Each dot represents one cell. (Right) Representative images of young adult (59-63 days) female heterozygous SP-oScarlet killifish brain sections, highlighting oScarlet protein signal (native fluorescence plus anti-mCherry immunofluorescence staining) in cells expressing *CSF1R* RNA (representative of n = 10 fish, 59-63 days females, over two experiments). Scale bars = 5 μm. **i,** UMAP as in **f** colored according to log-normalized *ELAVL3* (*LOC107378364*) expression. Each dot represents one cell.

We confirmed insertion of the SP-oScarlet knockin construct at the *ELAVL3* locus by PCR and long-read sequencing of the entire locus, which confirmed appropriate junctions with the *ELAVL3* gene and the presence of the SP-oScarlet insertion (**Fig. 1 – Fig. Supplement 1a**). We noted the presence of two tandem copies of the SP-oScarlet insertion, although the presence of a stop codon at the C-terminus of oScarlet likely leads to the production of a single oScarlet protein (**Fig. 1 – Fig. Supplement 1a**).

We verified that *ELAVL3* and *oScarlet* transcripts were expressed in the same cells of SP-oScarlet fish, as expected, using *in situ* hybridization chain reaction (HCR)^73^ on adult brain sections (**Fig. 1 – Fig. Supplement 1b-c**). By immunofluorescence staining for oScarlet, we also observed bright, punctate oScarlet signal in adult SP-oScarlet brains (**Fig. 1b**, **Fig. 1 – Fig. Supplement 1d**). Using HCR on brain sections, we verified the presence of cells with bright oScarlet protein signal (oScarlet^HIGH^) that did not express *oScarlet* transcripts in the brains of SP-oScarlet fish (**Fig. 1c**, **Fig. 1 – Fig. Supplement 1d**). This suggests that the oScarlet protein is efficiently secreted from the neurons that express the *oScarlet* transcript, resulting in oScarlet protein that is likely engulfed by other cell types in the brain. Thus, the bright oScarlet puncta in the brains of SP-oScarlet fish can be found within cells that likely engulf, and do not produce, this protein.

To identify the cells that engulf oScarlet, we developed a flow cytometry strategy to distinguish cells with bright oScarlet protein signal (oScarlet^HIGH^) from other cells in the adult brain (oScarlet^LOW^) (**Fig. 1d-e**, **Fig. 1 – Fig. Supplement 1e**). We isolated oScarlet^HIGH^ cells from adult SP-oScarlet brains (**Fig. 1e**), and we verified that these cells were not present in the brains of wildtype killifish, even at older ages where more autofluorescent cells might be expected (**Fig. 1 – Fig. Supplement 1f**).

To unbiasedly characterize oScarlet^HIGH^ cells, we performed single-cell RNA sequencing on enriched oScarlet^HIGH^ cells FACS-isolated from the brains of young adult (49 days) and old (134 days) male SP-oScarlet killifish (n = 4 fish per age group, pooled), as well as oScarlet^LOW^ cells isolated from the young adult fish. We used transcription and translation inhibitors to minimize gene expression changes due to dissociation, as done in studies of mammalian brain macrophages^74–76^. We performed Uniform Manifold Approximation and Projection (UMAP)^77^ analysis of the single-cell RNA sequencing data from young adult animals (for analysis of the old dataset, see **Fig. 4**). This analysis revealed that most oScarlet^HIGH^ cells were transcriptionally distinct from their oScarlet^LOW^ counterparts (**Fig. 1f**).

By manually annotating cell types based on matching top UMAP cluster marker genes (**Fig. 1 – Fig. Supplement 2a-b**) to cell type markers known from literature in mammals, zebrafish, and killifish^68,78–80^, we identified the vast majority (99.6%) of oScarlet^HIGH^ cells as myeloid immune cells (**Fig. 1g**).

oScarlet^LOW^ cells included neurons, oligodendroglia, radial glia and progenitors, pericytes, and T cells (**Fig. 1g**). We noted that endothelial cells were not well represented in our dataset, possibly due to our use of dissociation and FACS in our workflow. Only a very small percentage (0.5%) of oScarlet^LOW^ cells were myeloid cells, indicating that most myeloid cells likely engulfed oScarlet (**Fig. 1g**).

We could confirm by HCR on brain sections *in situ* that oScarlet^HIGH^ cells expressed a homolog of the conserved myeloid marker *CSF1R*^81,82^ (**Fig. 1h**). We also verified that oScarlet^HIGH^ cells largely did not express *ELAVL3*, the driver for *oScarlet* expression (**Fig. 1i**).

Collectively, these data show that most oScarlet^HIGH^ cells are myeloid cells that have likely engulfed secreted oScarlet protein in the brains of SP-oScarlet fish. Thus, we have generated a genetic model for chronic engulfment by brain myeloid cells in a short-lived vertebrate.

### The killifish brain includes a population of macrophages with transcriptional similarities to mammalian monocyte-derived macrophages

What type of myeloid cells engulf oScarlet in the killifish brain? Analysis of our single-cell RNA sequencing data indicated that oScarlet^HIGH^ cells are likely macrophages. Indeed, in addition to *CSF1R*, oScarlet^HIGH^ cells were enriched for *APOEB* (a homolog of human *APOE*) as well as antigen presentation-associated (e.g., *HLA-DPB1*, *CD74*), complement-associated (e.g., *C1QC*, *C1QA*), and scavenger receptor-encoding (e.g., *MRC1*, *MARCO*) genes (**Fig. 2a**). Moreover, by analyzing gene signatures from our single cell RNA sequencing dataset, we found that oScarlet^HIGH^ cells were enriched for signatures characteristic of macrophage functions, such as endocytosis-related and lysosomal genes (**Fig. 2b**). We also observed punctate oScarlet protein signal in proximity to late endosomes and lysosomes (marked by the protein marker LAMP1) by immunofluorescence staining (**Fig. 2 – Fig. Supplement 1a**), consistent with endocytosis and attempted degradation of extracellular material by macrophages. mScarlet, from which oScarlet is derived^70^, is known to be relatively stable at the acidic lysosomal pH^69^, which could explain the persistence of oScarlet signal in cells that engulf the protein.

**Figure 2.**
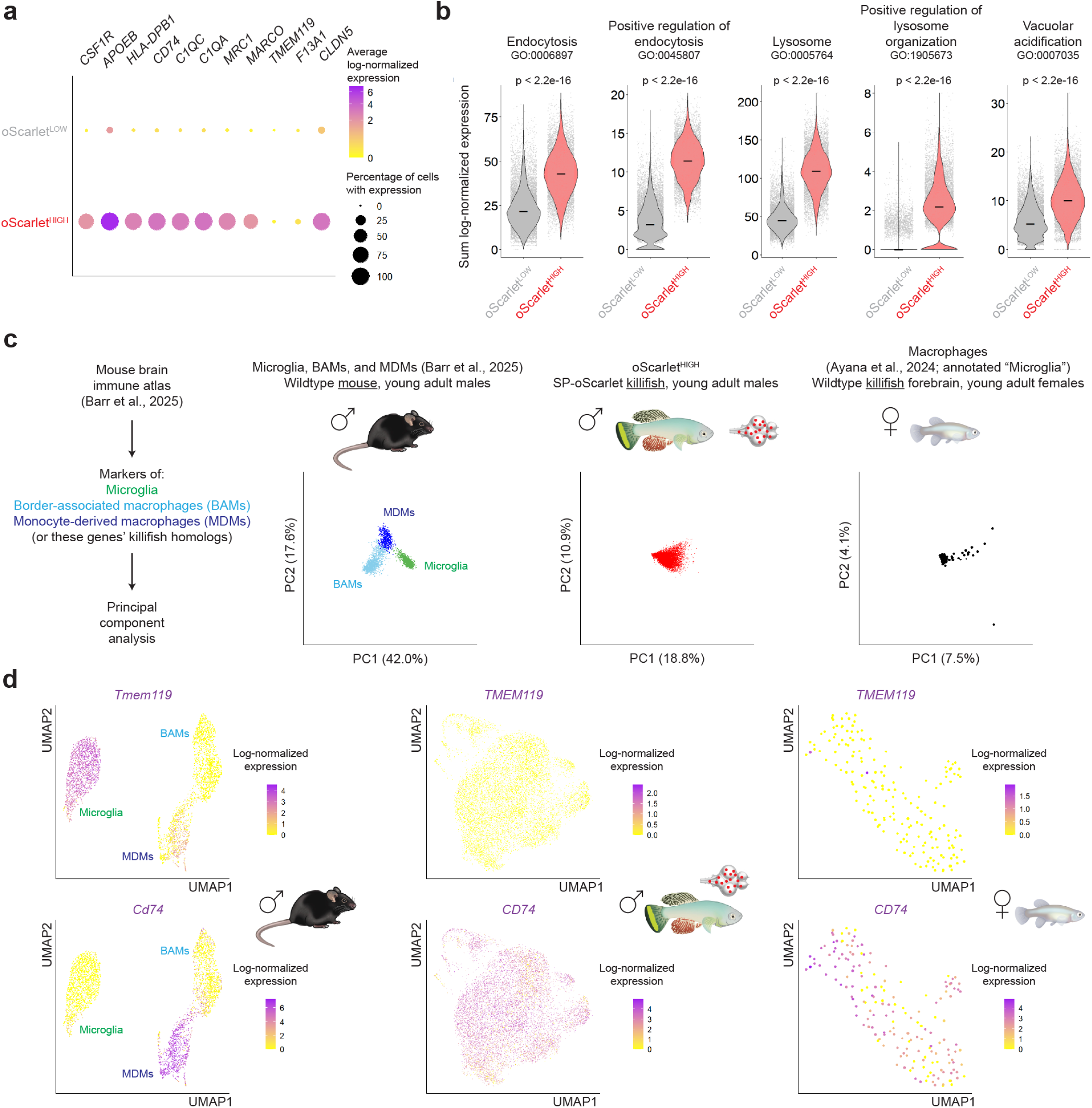
The killifish brain includes a population of macrophages with transcriptional similarities to mammalian monocyte-derived macrophages. **a,** Dotplot highlighting expression of selected genes (*CSF1R* (*LOC107387189*), *APOEB* (*LOC10737395*), *HLA-DPB1* (*LOC107372638*), *CD74* (*LOC107375728*), *C1QC* (*LOC107385068*), *C1QA* (*LOC107385067*), *MRC1* (*LOC107383768*), *MARCO* (*LOC107390282*), *TMEM119* (*LOC107393987*), *F13A1* (*LOC107377057*), and *CLDN5* (*LOC107394244*)) in young adult oScarlet^LOW^ and oScarlet^HIGH^ cells from experiment in **1d**. The color of each circle corresponds to average log-normalized expression among cells with detectable expression of the respective gene. The size of each circle corresponds to percentage of cells in each population with detectable expression of the gene. **b,** Violin plots for sum log-normalized expression of genes in Gene Ontology: Biological Process terms “endocytosis” (GO:0006897), “positive regulation of endocytosis” (GO:0045807), “lysosome” (GO:0005764), “positive regulation of lysosome organization” (GO:1905673), and “vacuolar acidification” (GO: 0007035) in young adult oScarlet^LOW^ and oScarlet^HIGH^ cells from experiment in **1d**. Each dot represents one cell. Lines: medians; p-values, unpaired Wilcoxon rank-sum test at per-cell level (fish were pooled for single cell RNA-sequencing experiments to obtain sufficient numbers of oScarlet^HIGH^ cells). **c,** Principal component analysis (PCA) plots (principal components 1 vs. 2) of wildtype young adult mouse microglia, border-associated macrophages (BAMs), and monocyte-derived macrophages (MDMs) (Barr et al., 2025^75^; 3,230 high-quality cells from clusters manually annotated by Barr et al.), young adult oScarlet^HIGH^ cells from experiment in **1d**, and wildtype young adult killifish forebrain macrophages (Ayana et al., 2024^68^; 176 high-quality cells from *APOEB*+ *IBA1*+ *LCP1*+ cluster) using marker genes of microglia, BAMs, and MDMs from the Barr et al. dataset, or all killifish homologs of these genes. Each dot represents one cell. Percentages on axes represent the percentage of variance explained by each respective principal component. Art by Rogelio Barajas. **d,** UMAPs of wildtype young adult mouse brain macrophages (Barr et al., 2025^75^; 3,488 high-quality cells from clusters manually annotated by Barr et al.), young adult oScarlet^HIGH^ cells from experiment in **1d**, and wildtype young adult killifish forebrain macrophages (Ayana et al., 2024^68^) colored according to log-normalized *Tmem119* (or killifish homolog *TMEM119* (*LOC107393987*)) or *CD74* (or killifish homolog *CD74* (*LOC107375728*)). Each dot represents one cell.

In the mammalian brain, most macrophages fall into one of three main classes^75,83^: parenchymal microglia (derived from the embryonic yolk sac), and two classes of macrophages found at interfaces between the brain and periphery (blood vessels, meninges, and choroid plexus) – border-associated macrophages (BAMs) (also derived from the embryonic yolk sac), and monocyte-derived macrophages (MDMs) (of peripheral, hematopoietic origin)^84–93^. BAMs and MDMs can strongly engulf substrates from the CSF^10^, with BAMs noted for their exceptional endocytic capacity^75^.

Using principal component analysis (PCA) with marker genes, mouse brain macrophages from a dataset from Barr et al.^75^ could be clearly transcriptionally separated into microglia, BAMs, and MDMs (**Fig. 2c**, **Fig. 2 – Fig. Supplement 1b**). In contrast, using principal component analysis with killifish homologs of the same marker genes, oScarlet^HIGH^ cells from SP-oScarlet killifish could not be transcriptionally separated into these classes (**Fig. 2c**, **Fig. 2 – Fig. Supplement 1b**). Consistent with this finding, wildtype killifish forebrain macrophages (i.e., cells from an *APOEB+ IBA1*+ *LCP1*+ cluster, originally annotated as “Microglia”) from an independent dataset from Ayana et al.^68^ could not be transcriptionally separated into microglia, BAMs, and MDMs in the same analysis (**Fig. 2c**, **Fig. 2 – Fig. Supplement 1b**). Thus, we did not find evidence for the existence of transcriptionally distinct microglia, BAMs, and MDMs in killifish brains, possibly suggesting a more plastic population of brain macrophages in the killifish.

Killifish homologs of many of the strongest markers of mouse microglia, BAMs, and MDMs were not detected in killifish brain macrophages, while some killifish brain macrophage markers (e.g., *CLDN5*) were not detected in mouse brain macrophages (**Fig. 2a**, **Fig. 2d**, **Fig. 2 – Fig. Supplement 2a**, **Fig. 2 – Fig. Supplement 3a**, **Fig. 2 – Fig. Supplement 4a**).

Nonetheless, the expression profile of oScarlet^HIGH^ cells overlapped most with that of mouse MDMs (**Fig. 2d**, **Fig. 2 – Fig. Supplement 2a**, **Fig. 2 – Fig. Supplement 3a**). Similar expression of key MDM markers was observed in many wildtype macrophages in the Ayana et al. dataset (**Fig. 2d**, **Fig. 2 – Fig. Supplement 2a**, **Fig. 2 – Fig. Supplement 4a**). Hence, the adult killifish brain includes a population of macrophages with some transcriptional similarities to mammalian monocyte-derived macrophages (consistent with studies in zebrafish^80,94,95^).

### Macrophages in the killifish brain are endocytic and present at brain borders

To functionally test whether oScarlet^HIGH^ cells have *in vivo* engulfment capabilities like mammalian BAMs and MDMs, we injected the macromolecule dextran (a 70 kDa polysaccharide, conjugated to a fluorophore) into the brains of young adult to middle-aged (63-91 days) female SP-oScarlet animals (**Fig. 3a**). We aimed our injection for the brain ventricle to maximize dextran distribution via the CSF^96^. In mice, similar approaches have been shown to result in engulfment of fluorescently-tagged substrates by BAMs and MDMs, and less so by microglia^10,97,98^.

**Figure 3.**
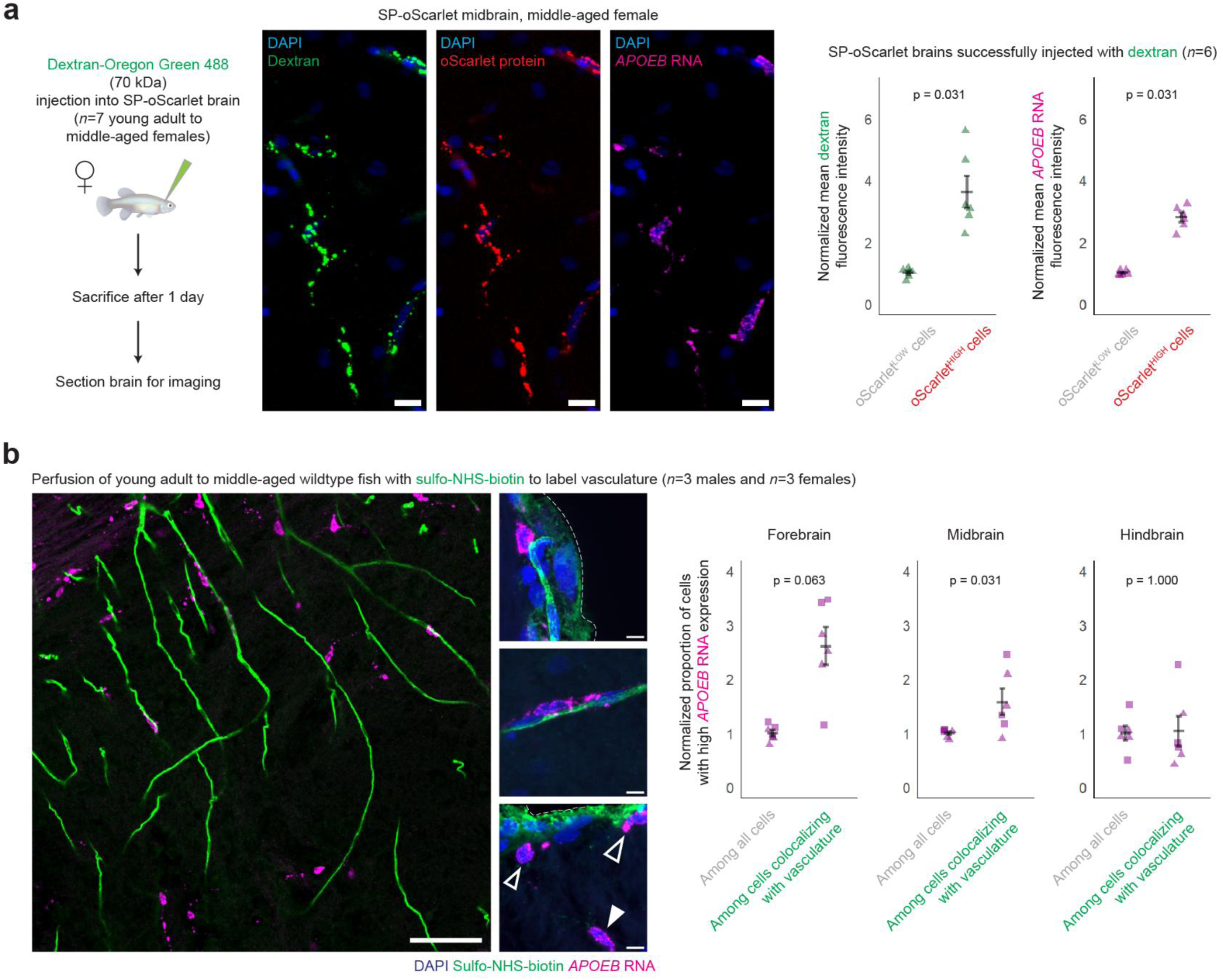
Macrophages in the killifish brain are endocytic and present at brain borders. **a,** (Left) Schematic for injection of dextran-Oregon Green 488 (70 kDa) into the brains of young adult to middle-aged (63-91 days) female heterozygous SP-oScarlet killifish (n = 7 fish over two experiments). (Center) Representative images of middle-aged (91 days) female heterozygous SP-oScarlet killifish brain sections highlighting co-occurrence of injected dextran signal, oScarlet protein signal (native fluorescence plus anti-mCherry immunofluorescence staining), and *APOEB* RNA in the same cells. Scale bars = 10 μm. (Right) Quantification of mean dextran fluorescence intensity and mean *APOEB* RNA fluorescence intensity in oScarlet^LOW^ and oScarlet^HIGH^ cells (n = 6 successfully injected fish, 63-91 days females, with 5-6 regions of interest over 1-2 sagittal brain sections per fish, over two experiments). Four images (one image per fish for each of four fish) of regions of interest with a visually apparent very high density of cells with high *APOEB* RNA expression, which could plausibly reflect an injection-related injury, were excluded from quantifications. The excluded images are listed in **Supplementary Table 4**. Each triangle represents one fish. Mean +/-standard error of mean; p-values, paired Wilcoxon rank-sum test at per-animal level. **b,** (Left) Representative images of young adult to middle-aged (74-101 days) male and female wildtype killifish brain sections, highlighting cells with high *APOEB* RNA expression that do and do not colocalize with vasculature as labeled by perfused sulfo-NHS-biotin (representative of n = 6 fish, 3 males, 74-101 days, and 3 females, 74-101 days, over two experiments). Scale bars = 50 μm (leftmost image), 5 μm (all others). Dashed lines: brain outside border. (In bottom right image) Filled arrows: example of cell with high *APOEB* RNA expression that does not colocalize with vasculature or other brain borders. Unfilled arrows: examples of cells with high *APOEB* RNA expression that are found near brain borders. (Right) Quantification of proportion of cells with high *APOEB* RNA expression among all cells and among cells colocalizing with vasculature, as defined by centroid-centroid distance to nearest sulfo-NHS-biotin-positive cell, in forebrain, midbrain, and hindbrain regions imaged (n = 6 fish, 74-101 days, with 1-2 regions of interest per brain region over 1-2 sagittal brain sections per fish, over two experiments). Regions of interest imaged generally avoided brain outside borders due to very high brightness of sulfo-NHS-biotin signal at some parts of brain outside borders, which could bias quantification. Each square (male) or triangle (female) represents one fish.

We performed HCR and immunofluorescence staining on brain sections of fish one day post-injection. Cells with bright oScarlet protein signal were enriched for dextran engulfment, as well as for expression of *APOEB*, as predicted by our single-cell RNA sequencing data (p = 0.031 for each, paired Wilcoxon test) (**Fig. 3a**, **Fig. 3 – Fig. Supplement 1a**). We verified that the colocalization seen between dextran and oScarlet signal (**Fig. 3a**) was not a result of bleed-through or autofluorescence (**Fig. 3 – Fig. Supplement 1b-c**) and likely resulted from these substrates being localized to the same endosomes and lysosomes. Collectively, these data show that killifish brain macrophages can engulf diverse extracellular substrates (protein, polysaccharide) (see **Fig. 5** for additional, *ex vivo* engulfment assays).

As noted above, in mammals, engulfment of substrates injected into the brain ventricles is a distinguishing functional characteristic of BAMs and MDMs, consistent with their localization at brain borders exposed to CSF flow^10,11^. To determine the spatial localization of killifish brain macrophages, we perfused young adult to middle-aged (74-101 days) male and female wildtype killifish with sulfo-NHS-biotin to label vasculature^99^ and performed HCR on brain sections for *APOEB*. We observed cells with high *APOEB* expression adjacent to (and on the parenchymal-facing side of, not within) vasculature in wildtype killifish brains, both near the brain’s external borders and within the brain interior (**Fig. 3b**). While we also observed some cells with high *APOEB* expression within the parenchyma, away from any detectable vasculature or other brain border, the distribution of cells with high *APOEB* expression was enriched toward sulfo-NHS-biotin-labeled vasculature in some brain regions (**Fig. 3b**, see **Source Data Table 5** for additional metrics of colocalization with vasculature).

Thus, we have identified a population of macrophages in the killifish brain with several characteristics of mammalian BAMs and MDMs – expressing several markers associated with brain macrophages including complement components and scavenger receptors, exhibiting endocytic functions, and being at least in part localized near brain borders.

### Killifish brain macrophages change transcriptionally with age in our model

We leveraged the killifish’s short lifespan to characterize brain macrophages with aging. We first used single-cell RNA sequencing to characterize oScarlet^HIGH^ cells from old SP-oScarlet killifish (same experiment as described in **Fig. 1d**). By performing UMAP analysis of oScarlet^HIGH^ cells from young adult and old SP-oScarlet brains, we found evidence of transcriptional differences between young adult and old oScarlet^HIGH^ cells (**Fig. 4a**). We found that oScarlet^HIGH^ cells from old brains were also predominantly macrophages, expressing markers (*CSF1R*, *APOEB*, *HLA-DPB1*, *C1QC*, *MRC1*) at similar levels to young adult oScarlet^HIGH^ cells (**Fig. 4b**).

**Figure 4.**
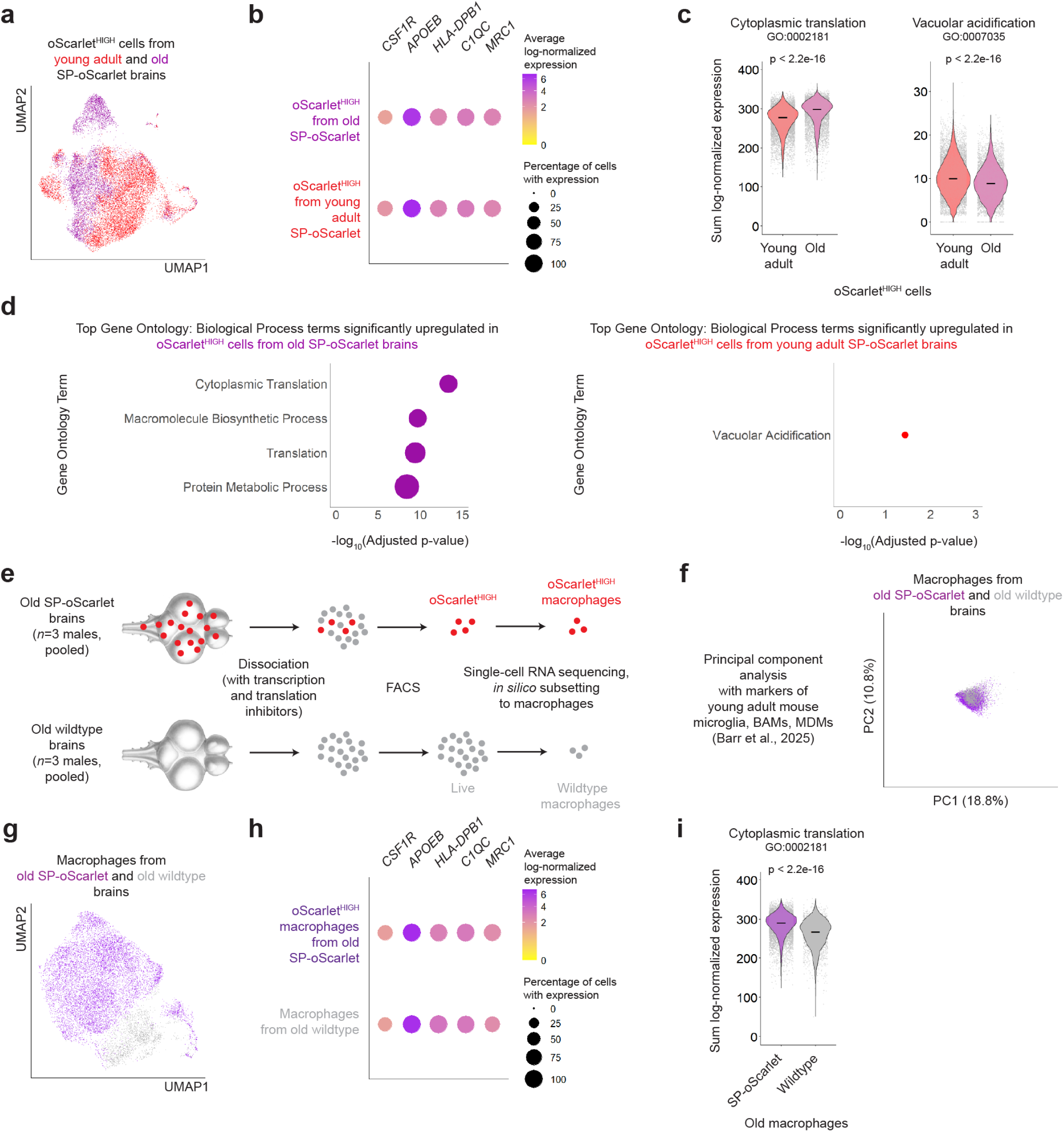
Killifish brain macrophages change transcriptionally with age in our model. **a,** UMAP of 6,489 oScarlet^HIGH^ high-quality cells from old (134 days) male heterozygous SP-oScarlet brains (n = 4 fish, pooled, in one experiment), as well as 9,680 young adult oScarlet^HIGH^ high-quality cells from experiment in **1d** (same experiment). **b,** Dotplot highlighting expression of selected homologs of mammalian brain macrophage marker genes (*CSF1R* (*LOC107387189*), *APOEB* (*LOC10737395*), *HLA-DPB1* (*LOC107372638*), *C1QC* (*LOC107385068*), and *MRC1* (*LOC107383768*)) in old and young adult oScarlet^HIGH^ cells from experiment in **1d**. The color of each circle corresponds to average log-normalized expression among cells with detectable expression of the respective gene. The size of each circle corresponds to percentage of cells in each population with detectable expression of the gene. **c,** Violin plots for sum log-normalized expression of genes in Gene Ontology: Biological Process terms “cytoplasmic translation” (GO:0002181) and “vacuolar acidification” (GO: 0007035) in young adult and old oScarlet^HIGH^ cells from experiment in **1d**. Each dot represents one cell. Lines: medians; p-values, unpaired Wilcoxon rank-sum test at per-cell level. **d**, Top Gene Ontology: Biological Process terms upregulated in old (left) and young adult (right) oScarlet^HIGH^ cells from experiment in **1d**, as ranked by adjusted p-value (Fisher’s exact test, Benjamini-Hochberg false discovery rate (FDR)-corrected). Size of each circle corresponds to number of distinct human homologs among differentially expressed genes from each term. Full lists of differentially expressed genes and upregulated terms in **Supplementary Tables 9-11**. **e,** Schematic for single-cell RNA sequencing of oScarlet^HIGH^ cells from old (130 days) male heterozygous SP-oScarlet killifish brains and all Live cells from old (122 days) male wildtype killifish brains (n = 3 fish per genotype, pooled in one experiment; for an independent experiment involving old SP-oScarlet fish, see **1d**). **f**, PCA plot (principal components 1 vs. 2) of oScarlet^HIGH^ and wildtype old killifish brain macrophages from experiment in **e** using all killifish homologs of marker genes of microglia, BAMs, and MDMs (Barr et al., 2025^75^). Each dot represents one cell. Percentages on axes represent the percentage of variance explained by each respective principal component. **g,** UMAP of 12,575 old oScarlet^HIGH^ and old wildtype macrophages, as defined by *APOEB*+ *IBA1*+ *LCP1*+ clusters among all cells from experiment in **e** (resolution = 0.2). Each dot represents one cell. **h,** Dotplot highlighting expression of selected homologs of mammalian brain macrophage marker genes (*CSF1R* (*LOC107387189*), *APOEB* (*LOC10737395*), *HLA-DPB1* (*LOC107372638*), *C1QC* (*LOC107385068*), and *MRC1* (*LOC107383768*)) in old oScarlet^HIGH^ and old wildtype macrophages from experiment in **e**. The color of each circle corresponds to average log-normalized expression among cells with detectable expression of the respective gene. The size of each circle corresponds to percentage of cells in each population with detectable expression of the gene. **i,** Violin plot for sum log-normalized expression of genes in Gene Ontology: Biological Process term “cytoplasmic translation” (GO:0002181) in old oScarlet^HIGH^ and old wildtype macrophages from experiment in **e**. Each dot represents one cell. Lines: medians; p-values, unpaired Wilcoxon rank-sum test at per-cell level.

What transcriptional pathways in these killifish brain macrophages are impacted during aging? Gene ontology (GO) analysis showed an upregulation of translation-related pathways with age in oScarlet^HIGH^ cells (**Fig. 4c-d**). GO analysis also identified a modest downregulation in vacuolar acidification with age (**Fig. 4c-d**). Analysis of specific pathways relevant to engulfment and endolysosomal functions showed that these were also modestly decreased during aging (**Fig. 4 – Fig. Supplement 1a**). Inflammation-related genes were not strongly upregulated with age in oScarlet^HIGH^ cells (**Fig. 4 – Fig. Supplement 1b**), unlike in brain macrophage populations described in mammals^75,79,100,101^.

Could some of the transcriptional responses observed with age be due to the lifelong, chronic engulfment of oScarlet, particularly given the relative resistance of oScarlet to rapid degradation? To address this possibility, we performed an independent single-cell RNA sequencing experiment of oScarlet^HIGH^ cells FACS-sorted from the brains of old (130 days) SP-oScarlet male animals (n = 3 fish, pooled), as well all Live cells FACS-sorted from the brains of old (122 days) wildtype male brains (n = 3 fish, pooled) (**Fig. 4e**). We subset each population to macrophages (i.e., cells from *APOEB+ IBA1*+ *LCP1*+ clusters).

We found that neither old oScarlet^HIGH^ nor old wildtype macrophages could be transcriptionally separated into microglia, BAMs, and MDMs (**Fig. 4f**, **Fig. 4 – Fig. Supplement 1c**). When comparing old oScarlet^HIGH^ macrophages to old wildtype macrophages, we found that these populations expressed marker genes (*CSF1R*, *APOEB*, *HLA-DPB1*, *C1QC*, *MRC1*) at similar levels (**Fig. 4g-h**).

GO analysis showed that translation-related pathways were enriched in old oScarlet^HIGH^ macrophages compared to wildtype macrophages (**Fig. 4i**, **Fig. 4 – Fig. Supplement 1d**).

Some inflammation- and pathogen-response-related pathways were mildly depleted in old oScarlet^HIGH^ macrophages compared to old wildtype macrophages (**Fig. 4 – Fig. Supplement 1d**). The engulfment-related pathways examined showed no consistent directional pattern when comparing old oScarlet^HIGH^ to old wildtype macrophages (**Fig. 4 – Fig. Supplement 1e**). These results suggest that the translation-related changes observed with age in oScarlet^HIGH^ cells may reflect a response to lifelong oScarlet exposure, rather than to aging *per se*. In contrast, changes in engulfment-related pathways are unlikely to be solely driven by lifelong, chronic oScarlet exposure.

Overall, our single-cell RNA sequencing results indicate that killifish brain macrophages in SP-oScarlet animals experience notable transcriptional changes with age, with translation-related genes increasing in expression with age (possibly as part of a response to chronic oScarlet engulfment) and some engulfment-relevant (vacuolar acidification) genes decreasing in expression with age.

### Engulfment capacity of killifish brain macrophages declines with age in our model

We next functionally characterized killifish brain macrophages with age, using oScarlet^HIGH^ status as a proxy for macrophage identity – as supported by our single-cell RNA sequencing data – given the lack of antibodies and reporter lines in killifish to otherwise identify these cells. By performing flow cytometry on the brains of young adult (41-63 days) and old (148-167 days) male and female SP-oScarlet killifish (n = 6 fish per age group and per sex), we noted a decline with age in the oScarlet fluorescence intensity of oScarlet^HIGH^ cells in both males and females (p = 0.002 for each sex, unpaired Wilcoxon test) (**Fig. 5a**, **Fig. 5 – Fig. Supplement 1a**). These observations are consistent with the possibility that aging is accompanied by a potential decline in engulfment by brain macrophages, though the decline in oScarlet brightness in old oScarlet^HIGH^ cells could also be due to changes in oScarlet expression, secretion, or degradation, especially given that *ELAVL3* gene expression has been shown to decrease with age in the killifish brain (**Fig. 5 – Fig. Supplement 1b**)^55^.

**Figure 5.**
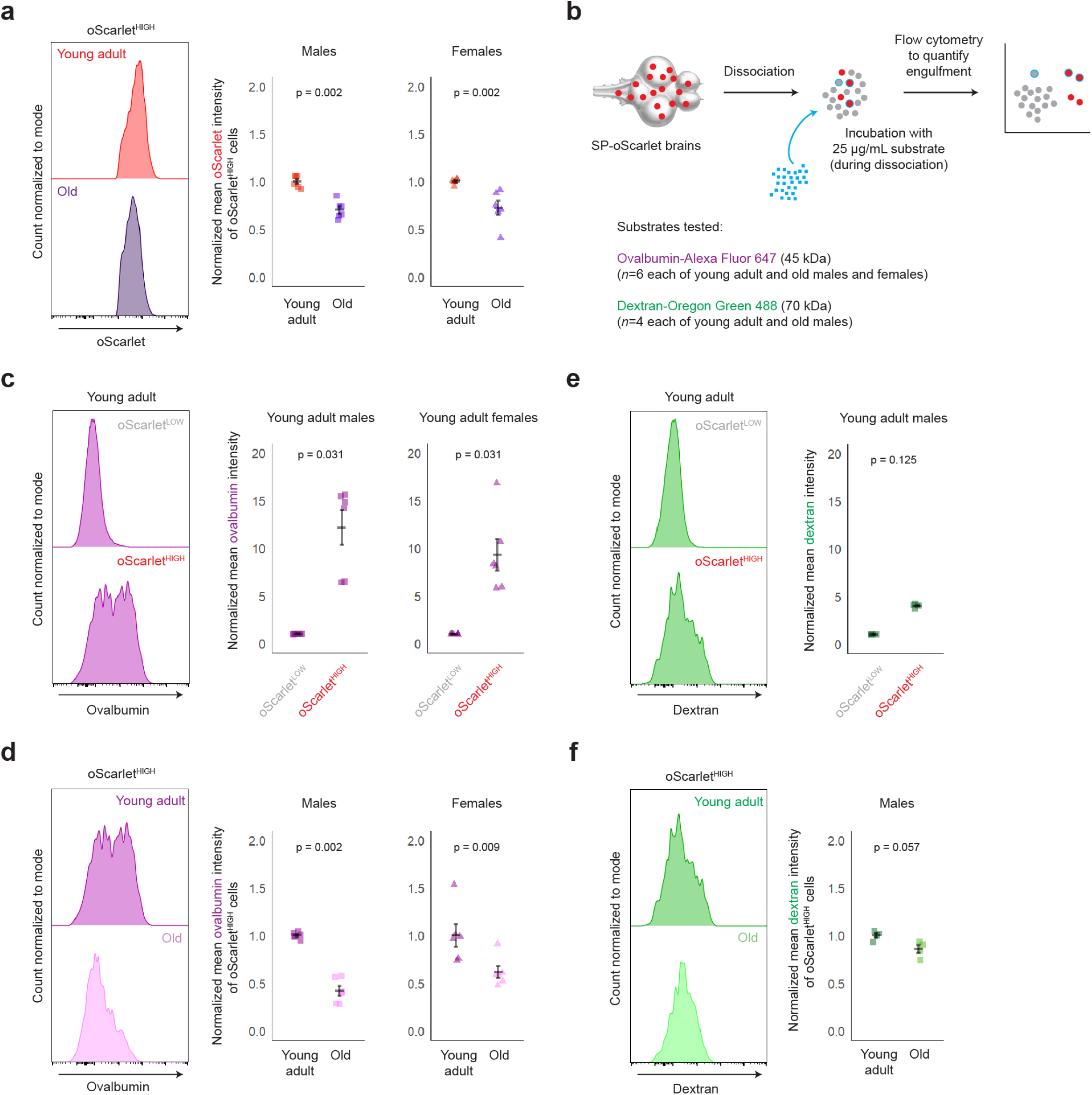
Engulfment capacity of killifish brain macrophages declines with age in our model. **a,** (Left) Representative flow cytometry histograms from young adult (41 days, above) and old (160 days, below) male heterozygous SP-oScarlet killifish brains, highlighting mean oScarlet fluorescence intensity in oScarlet^HIGH^ cells. (Right) Quantification of mean oScarlet fluorescence intensity among young adult and old oScarlet^HIGH^ cells (n = 6 fish per age per sex, over two experiments per sex; young adult males: 41-63 days; old males: 148-160 days; young adult females: 57-63 days; old females: 161-167 days). Each square (male) or triangle (female) represents one fish. Mean +/- standard error of mean; p-values, unpaired Wilcoxon rank-sum test at per-animal level. **b,** Schematic for *ex vivo* engulfment assays performed on young adult and old heterozygous SP-oScarlet killifish brains (for ovalbumin-Alexa Fluor^®^ 647: n = 6 fish per age and sex, over two experiments per sex; young adult males: 41-63 days; old males: 148-160 days; young adult females: 57-63 days; old females: 161-167 days; for dextran-Oregon Green 488 (70 kDa): n = 4 male fish per age in one experiment; young adult: 56 days; old: 156 days). **c,** (Left) Representative flow cytometry histograms from young adult (41 days) male heterozygous SP-oScarlet killifish brains, highlighting mean ovalbumin fluorescence intensity in oScarlet^LOW^ and oScarlet^HIGH^ cells. (Right) Quantification of mean ovalbumin fluorescence intensity among young adult oScarlet^LOW^ and oScarlet^HIGH^ cells (same experiments as in **a**). Each square (male) or triangle (female) represents one fish. Mean +/- standard error of mean; p-values, paired Wilcoxon rank-sum test at per-animal level. **d,** (Left) Representative flow cytometry histograms from young adult (41 days) (above) and old (160 days) (below) male heterozygous SP-oScarlet killifish brains, highlighting mean ovalbumin fluorescence intensity in oScarlet^HIGH^ cells. (Right) Quantification of mean ovalbumin fluorescence intensity among young adult and old oScarlet^HIGH^ cells (same experiments as in **a**). Each square (male) or triangle (female) represents one fish. Mean +/- standard error of mean; p-values, unpaired Wilcoxon rank-sum test at per-animal level. **e,** (Left) Representative flow cytometry histograms from young adult (56 days) male heterozygous SP-oScarlet killifish brains, highlighting mean dextran fluorescence intensity in oScarlet^LOW^ and oScarlet^HIGH^ cells. (Right) Quantification of mean dextran fluorescence intensity among young adult oScarlet^LOW^ and oScarlet^HIGH^ cells. Each square represents one fish. Mean +/- standard error of mean; p-values, paired Wilcoxon rank-sum test at per-animal level. **f,** (Left) Representative flow cytometry histograms from young adult (56 days) (above) and old (156 days) (below) male heterozygous SP-oScarlet killifish brains, highlighting mean dextran fluorescence intensity in oScarlet^HIGH^ cells. Each dot represents one cell. (Right) Quantification of mean dextran fluorescence intensity among young adult and old oScarlet^HIGH^ cells (same experiment as in **e**). Each square represents one fish. Mean +/-standard error of mean; p-values, unpaired Wilcoxon rank-sum test at per-animal level.

To assess more directly if the engulfment capacity of brain macrophages changes with age, we developed an acute *ex vivo* assay in the context of SP-oScarlet animals. In this assay, macromolecule substrates such as ovalbumin (a 45 kDa protein) and dextran (a 70 kDa polysaccharide) (conjugated to far-red and green fluorophores, respectively) were mixed *ex vivo* with brain cells during the 30 minutes of tissue dissociation^75^, followed by flow cytometry to quantify fluorescent substrate engulfment by cells (**Fig. 5b**). Using this assay, we verified that oScarlet^HIGH^ cells from young adult SP-oScarlet brains were enriched for their ability to rapidly engulf ovalbumin compared to oScarlet^LOW^ cells (p = 0.031 for each sex, paired Wilcoxon test) (**Fig. 5c**, see **Fig. 5 – Fig. Supplement 1c** for additional metrics of engulfment).

We used this *ex vivo* engulfment assay to examine ovalbumin engulfment differences with age among oScarlet^HIGH^ cells in both males and females (n = 6 fish per age and per sex). Crucially, we showed that oScarlet^HIGH^ cells in old brains showed a decreased capacity to engulf ovalbumin compared their counterpart cells in young adult brains, in both males (p = 0.002, unpaired Wilcoxon test) and females (p = 0.009, unpaired Wilcoxon test) (**Fig. 5d**). Fewer old oScarlet^HIGH^ cells engulfed ovalbumin, and those that did showed reduced ovalbumin brightness compared to young adult oScarlet^HIGH^ cells that engulfed ovalbumin (**Fig. 5 – Fig. Supplement 1d**, see **Fig. 5 – Fig. Supplement 2a-c** for alternative gating schemes).

In a separate experiment, we also performed this *ex vivo* engulfment assay using dextran as a substrate (n = 4 male fish per age) and verified that oScarlet^HIGH^ cells from young adult (56 days) male SP-oScarlet brains also showed a trending, non-significant increased capacity to rapidly engulf dextran compared to oScarlet^LOW^ cells (p = 0.125, unpaired Wilcoxon test) (**Fig. 5e**, see **Fig. 5 – Fig. Supplement 1e** for additional metrics of engulfment). oScarlet^HIGH^ cells in old (156 days) male brains also showed a trending, non-significant decreased capacity to engulf dextran compared their counterpart cells in young adult brains (p = 0.057, unpaired Wilcoxon test) (**Fig. 5f**). Fewer old oScarlet^HIGH^ cells engulfed dextran, and those that did showed reduced dextran brightness compared to young adult oScarlet^HIGH^ cells that engulfed dextran (**Fig. 5 – Fig. Supplement 1f**, see **Fig. 5 – Fig. Supplement 2d-e** for alternative gating schemes).

Collectively, these data suggest that brain macrophages in killifish exhibit a functional decline in engulfment capacity with age.

## Discussion

In this study, we generated a genetically modified killifish line to investigate engulfment of a secreted protein in the vertebrate brain, providing a model to study clearance. Using the secreted red fluorescent protein oScarlet as a model extracellular protein (here secreted principally from neurons), we found that a macrophage population in the adult killifish brain exhibits a high engulfment capacity. Our *in vivo* genetic model could be used to study secretion and engulfment in the vertebrate brain, help elucidate functional roles of brain macrophages, and identify interventions that boost engulfment in the aged or diseased brain. We note that some limitations of this genetic model include that oScarlet is expressed constitutively and is relatively resistant to degradation, precluding acute examination of oScarlet engulfment and potentially even affecting the state of cells that engulf oScarlet. We therefore developed additional methods in this study to assess acute engulfment with other substrates both *in vivo* and *ex vivo* in the killifish brain. Together, these tools in killifish complement recent approaches to study engulfment in mouse brains^10,75^, and they provide a resource to understand the functions of brain macrophages in a short-lived vertebrate.

We found that many killifish brain macrophages display similarities to border-associated macrophages (BAMs) and monocyte-derived macrophages (MDMs), known engulfment-capable macrophage populations in mammalian central nervous systems^10,75,76,89,102–115^. These similarities include: expression of marker genes such as scavenger receptors, localization to brain border regions, and functional enrichment for engulfment capacity for several substrates (the protein oScarlet *in vivo*, the polysaccharide dextran *in vivo* and the protein ovalbumin and polysaccharide dextran *ex vivo*). Our findings suggest that the adult killifish brain – in contrast to the mouse brain^102^ – represents a system where the need for waste clearance may be addressed by relatively abundant cells with characteristics of BAMs and MDMs, parallel to findings from adult zebrafish brains^80,94,95^. This potential evolutionary difference is intriguing and could be due to differences in myelination^116–118^, neurogenesis^119–121^, or demands for pathogen response^122,123^ or immune regulation^124^ between fish and mammals or between aquatic and terrestrial environments. These observations also suggest there may be more plasticity among fish brain macrophages than previously anticipated. Understanding how macrophages other than microglia perform brain clearance in species where they are abundant could provide new avenues to promote clearance in mammals.

We showed that in our model, killifish brain macrophages decline in engulfment capacity with age. This is in line with recent mammalian studies highlighting that BAMs and MDMs, despite being rare populations in the mammalian brain, play key functional roles but experience alterations with age and under pathological conditions^10,11,75,97,98,100,125–167^. Thus, BAMs and MDMs could represent promising avenues for therapeutic intervention in age-associated and other brain diseases. In mammalian BAMs, decline in engulfment with age has been linked to decreased expression of clathrin-mediated endocytosis receptors^75^. While similar transcriptional changes with age were not observed in our single-cell RNA sequencing datasets, key endocytic machinery in the killifish brain may be regulated at the level of translation (which has been shown to decrease globally with age in the killifish brain^54^), or at a post-translational level (e.g., decreased efficiency of proton pumping to acidify lysosomes^168^), resulting in an evolutionarily conserved decline in scavenging capacity with age.

Overall, our findings highlight the underappreciated role that engulfment by macrophages, particularly those at brain borders, could play in critical waste clearance processes in the brain. This study also underscores the potential of the killifish as a powerful vertebrate model to investigate questions relevant to aging, disease, and evolution of the complex and specialized cell types of the vertebrate brain.

## Acknowledgements

We thank Jason W. Miklas, Nimrod Rappoport, and Daniel J. Richard for critical reading of the manuscript; Natalie Schmahl and Sarah R. Boyle for help with killifish colony management; Jacob Chung, Rishad Khondker, Maddie Housh, Jason Yang, and Ian H. Guldner for help with killifish husbandry; Justin You, Aleksandra S. Tsenter, Merve Heinzer-Avar, Daniel Heinzer-Avar, and Julliana Ramirez-Matias for advice on brain sectioning and imaging analysis; Giulia Notarangelo, Andy P. Tsai, and Jaeyoon Lee for advice on flow cytometry; Olivia Y. Zhou for advice on design of single-cell RNA sequencing experiments, Nimrod Rappoport and Daniel J. Richard for advice on single-cell RNA sequencing data analysis; Alec J. Walker and Constanze Depp for advice on interpretation of single-cell RNA sequencing data; Ryann M. Fame and Blake J. Laham for advice on brain injections; Katrin I. Andreasson, Kyle G. Daniels, Andrew Z. Fire, Daniel F. Jarosz, and Judith Frydman, Charu Ramakrishnan, Lu Zhou, Gil Vantomme, Katharina Papsdorf, Eric D. Sun and all of the Brunet lab for helpful discussions and advice; and Nimrod Rappoport and Daniel J. Richard for code-checking. This project was supported, in part, by Award Number 1S10OD034400-01A1 from the National Center for Research Resources (NCRR). Its contents are solely the responsibility of the authors and do not necessarily represent the official views of the NCRR or the National Institutes of Health. Funding for this study was provided by the Simons Foundation (to A.B., B.S., T.W.C., and P.M.), the Knight Initiative for Brain Resilience (to A.B., T.W.C., and R.D.N.), the Glenn Foundation for Medical Research (to A.B.), the NOMIS Foundation (to A.B. and T.W.C.), the National Institute on Aging K99 Pathway to Independence Award (to R.D.N. and C.N.B.), the Simons Foundation Fellows-to-Faculty Award (to J.B.J.), and the Stanford Knight-Hennessy Scholars (to R.N.).

## Author contributions

This study was conceived by R.N., R.D.N., C.N.B., and A.B. R.N. designed the SP-oScarlet killifish line, and R.D.N. generated this line with help from R.N.. C.N.B. designed probes for some hybridization chain reaction experiments and checked code. R.N. designed and performed all experiments and analyses unless noted. A.N.P. performed structured illumination microscopy for LAMP1 imaging and helped with brain dissociations and FACS for *ex vivo* engulfment assays and single-cell RNA sequencing. P.R.K. (under the guidance of B.W.) performed cell type evolution analysis that aided interpretation of single-cell RNA sequencing data. H.J.B. (under the guidance of B.S.) helped design *ex vivo* engulfment assays and interpret single-cell RNA sequencing data, including by sharing unpublished mouse single-cell RNA sequencing datasets. A.M.M.J. (under the guidance of K.R.H.) performed brain clearing and 3D imaging. S.V.P. helped with brain injections. E.K.C. (under the guidance of T.W.C.) and J.C. performed sulfo-NHS-biotin perfusion for vasculature colocalization analysis. J.C. also helped with sample processing for single-cell RNA sequencing. F.B. helped with brain dissociations and flow cytometry for *ex vivo* engulfment assays and checked code. L.A.S. (under the guidance of S.R.Q.) and P.N.N. helped with 10x Genomics single-cell RNA sequencing. J.B.J. (under the guidance of P.M.) helped develop the brain injection protocol. E.K.C. (under the guidance of T.W.C.) and R.B. helped develop the FACS sorting strategy. R.B. also helped with brain sectioning and immunofluorescence staining optimization. P.P.S. produced the gene names conversion table used to identify homologs across species. The manuscript was written by R.N. and A.B., and all authors provided comments.

## Data and code availability

Sequencing reads will be made publicly available at NCBI GEO upon publication. Seurat objects generated in this study are available at Dropbox (<u>link</u>). Other raw data (images, etc.) will be made available upon request. Code is available at Github (https://github.com/RNagvekar/killifish_brain_macrophages).

---

**Figure 1 – Figure Supplement 1.**
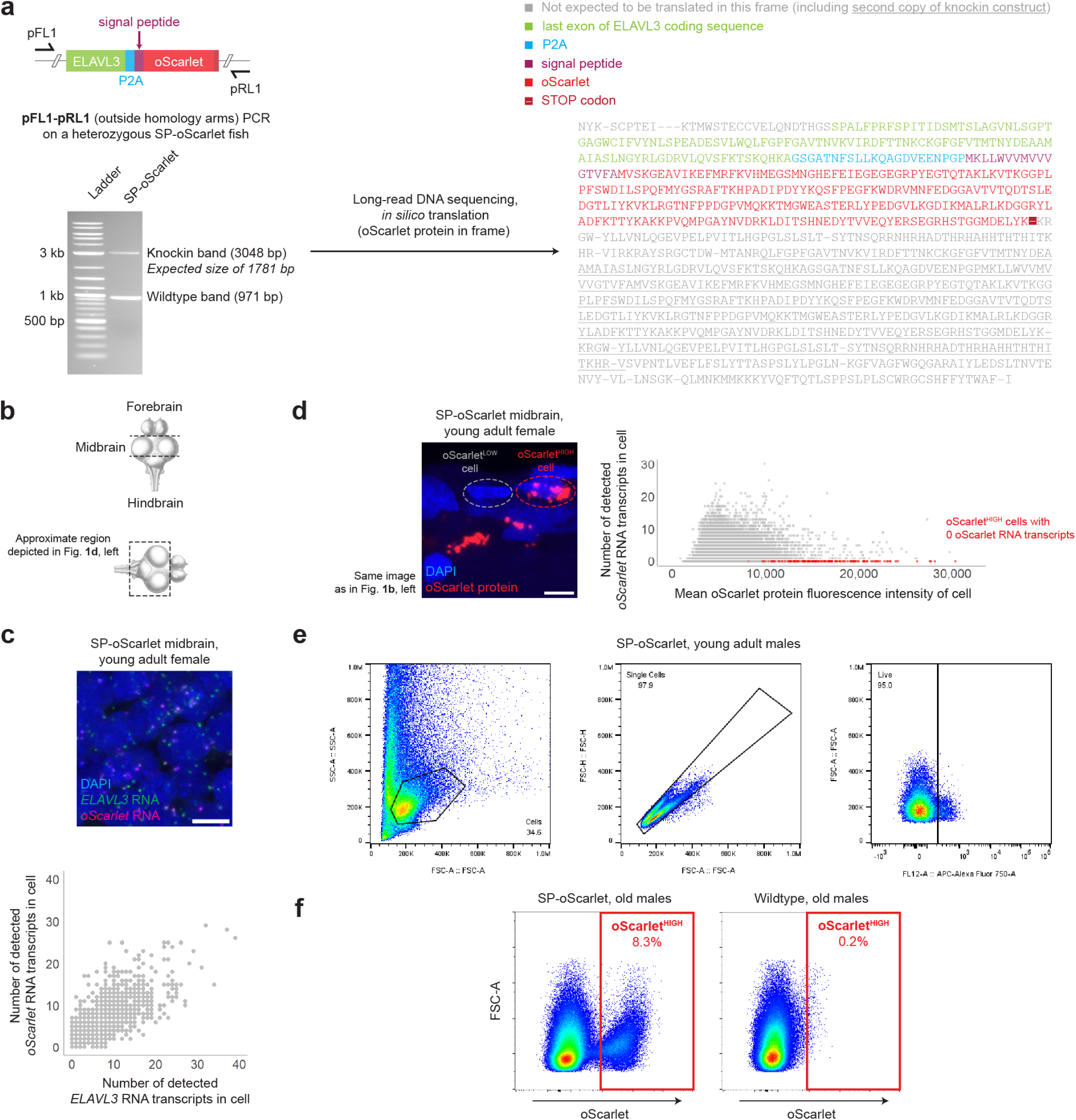
**a,** (Left) Schematic for PCR genotyping of a middle-aged (78 days) male heterozygous SP-oScarlet killifish, and DNA gel result (representative of n = 2 fish, 78 days males, in one experiment). (Right) *In silico* translated sequence of sequenced SP-oScarlet insertion (n = 1 fish, 78 days male). **b**, Representation of killifish brain highlighting approximate brain regions as referred to in this study. Art by Massimo Demma (adapted with permission from D’Angelo et al., 2013^169^). **c,** (Above) Representative image of a young adult (59 days) female heterozygous SP-oScarlet killifish brain section, highlighting *ELAVL3* transcripts and *oScarlet* transcripts in the periventricular gray zone, a neuron-dense region of the killifish midbrain. Scale bar = 5 μm. (Below) Per-cell quantification of number of detected *ELAVL3* transcripts and number of detected *oScarlet* transcripts (1,498 cells from n = 4 fish, 59 days females, with 1 region of interest in 1 sagittal brain section per fish, in one experiment). **d**, Same image as in **1b** (left), highlighting examples of oScarlet^LOW^ and oScarlet^HIGH^ cells as detected *in situ*. (Below) Per-cell quantification of mean oScarlet fluorescence intensity and number of detected *oScarlet* transcripts from experiment in **1c** (25,622 cells from n = 10 fish, 59-63 days females, with 3-6 regions of interest over 1-2 sagittal brain sections per fish, over two experiments). Each dot represents one cell. oScarlet^HIGH^ cells that have 0 detected *oScarlet* RNA transcripts are highlighted in red. **e,** FACS plots highlighting gating scheme used to isolate Live cells (as depicted in **1e**) from dissociated young adult male SP-oScarlet brains from experiment in **1d**. **f,** FACS plots highlighting oScarlet^HIGH^ cells notably detected in old (130 days old) male heterozygous SP-oScarlet but not in old (122 days old) male wildtype brains (n = 3 fish per genotype, pooled in one experiment). Each dot represents one cell.

**Figure 1 – Figure Supplement 2.**
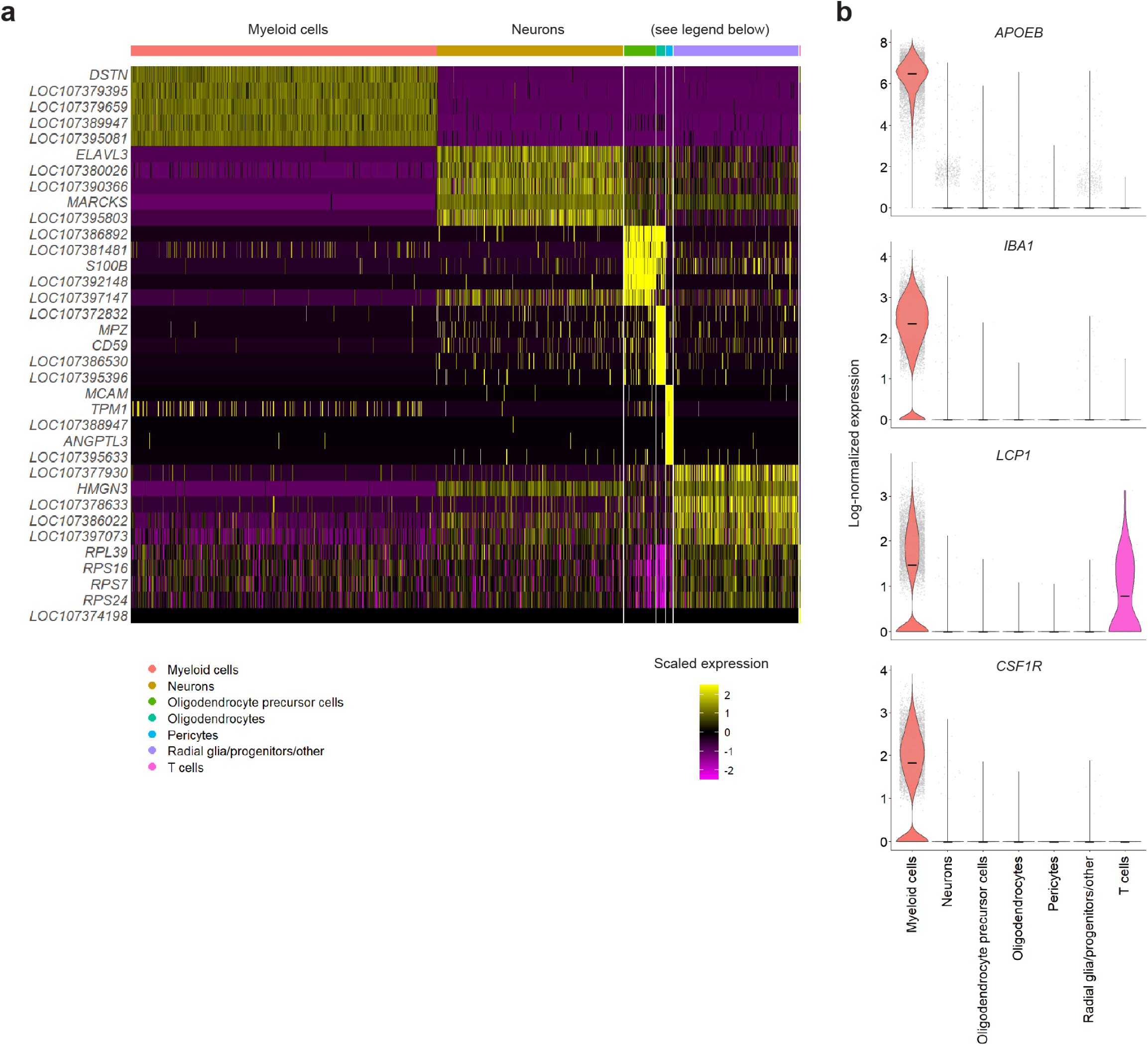
**a,** Heatmap of expression of top 5 marker genes per cell type, as ranked by area under curve (Presto), in every young adult cell from experiment in **1d**. Full list of cell type marker genes in **Supplementary Table 8**. **b,** Violin plots from experiment in **1d** highlighting expression of *APOEB* (*LOC10737395*), *IBA1* (*LOC107378674*), *LCP1* (*LOC107389591*), and *CSF1R* (*LOC107387189*) in each cell type. Each dot represents one cell. Lines: medians.

**Figure 2 – Figure Supplement 1.**
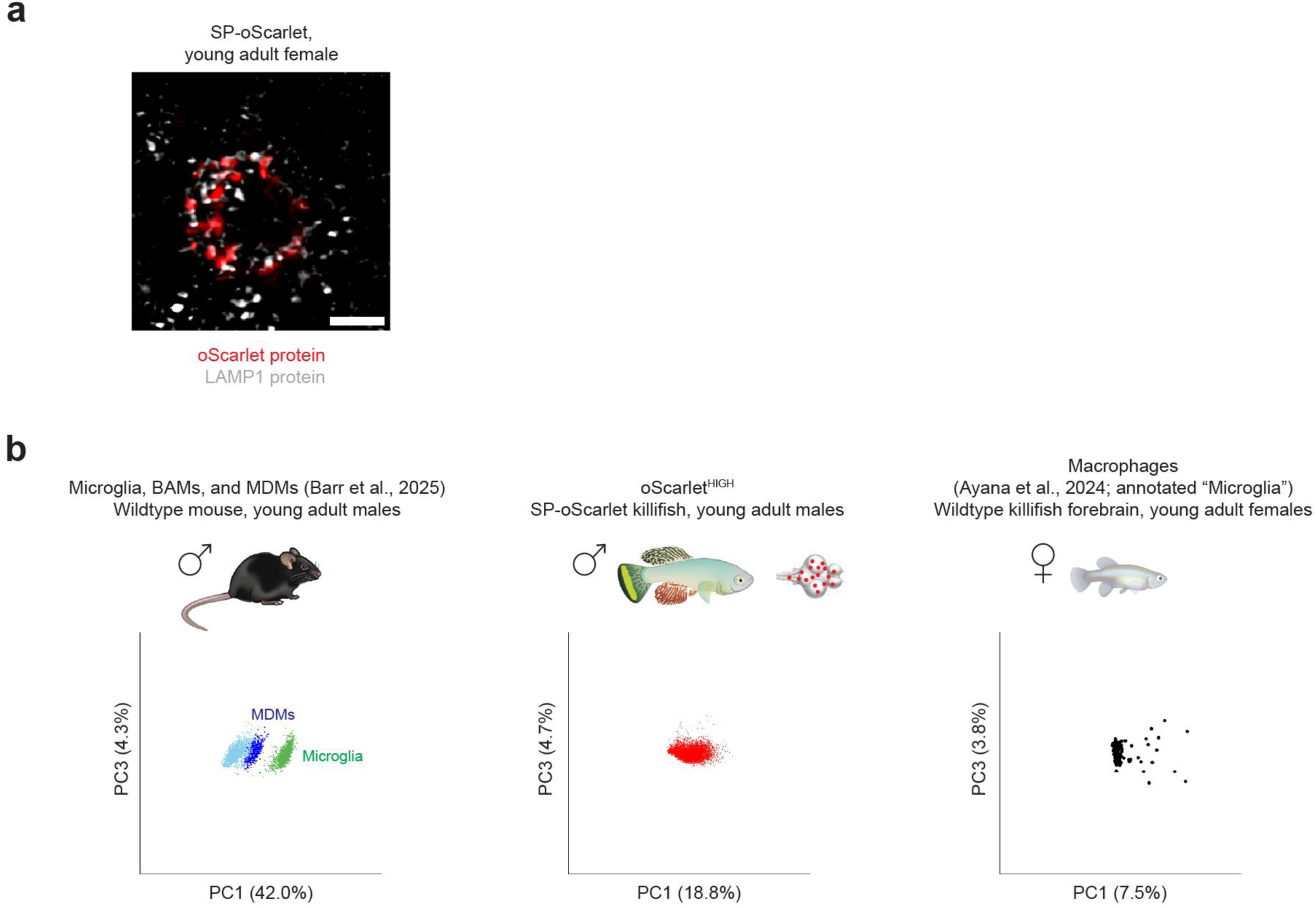
**a,** Representative image of a young adult (59 days) female heterozygous SP-oScarlet killifish brain section, highlighting oScarlet protein signal colocalized with the late endosome and lysosome marker LAMP1 in an SP-oScarlet brain (representative of n = 8 fish, 59-63 days females, over two experiments). Scale bar = 2 μm. **b,** Principal component analysis (PCA) plots (principal components 1 vs. 3) of wildtype young adult mouse brain macrophages (Barr et al., 2025^75^), young adult oScarlet^HIGH^ cells from experiment in **1d**, and wildtype young adult killifish forebrain macrophages (Ayana et al., 2024^68^) using marker genes of microglia, BAMs, and MDMs from the Barr et al. dataset, or all killifish homologs of these genes. Each dot represents one cell. Percentages on axes represent the percentage of variance explained by each respective principal component.

**Figure 2 – Figure Supplement 2.**
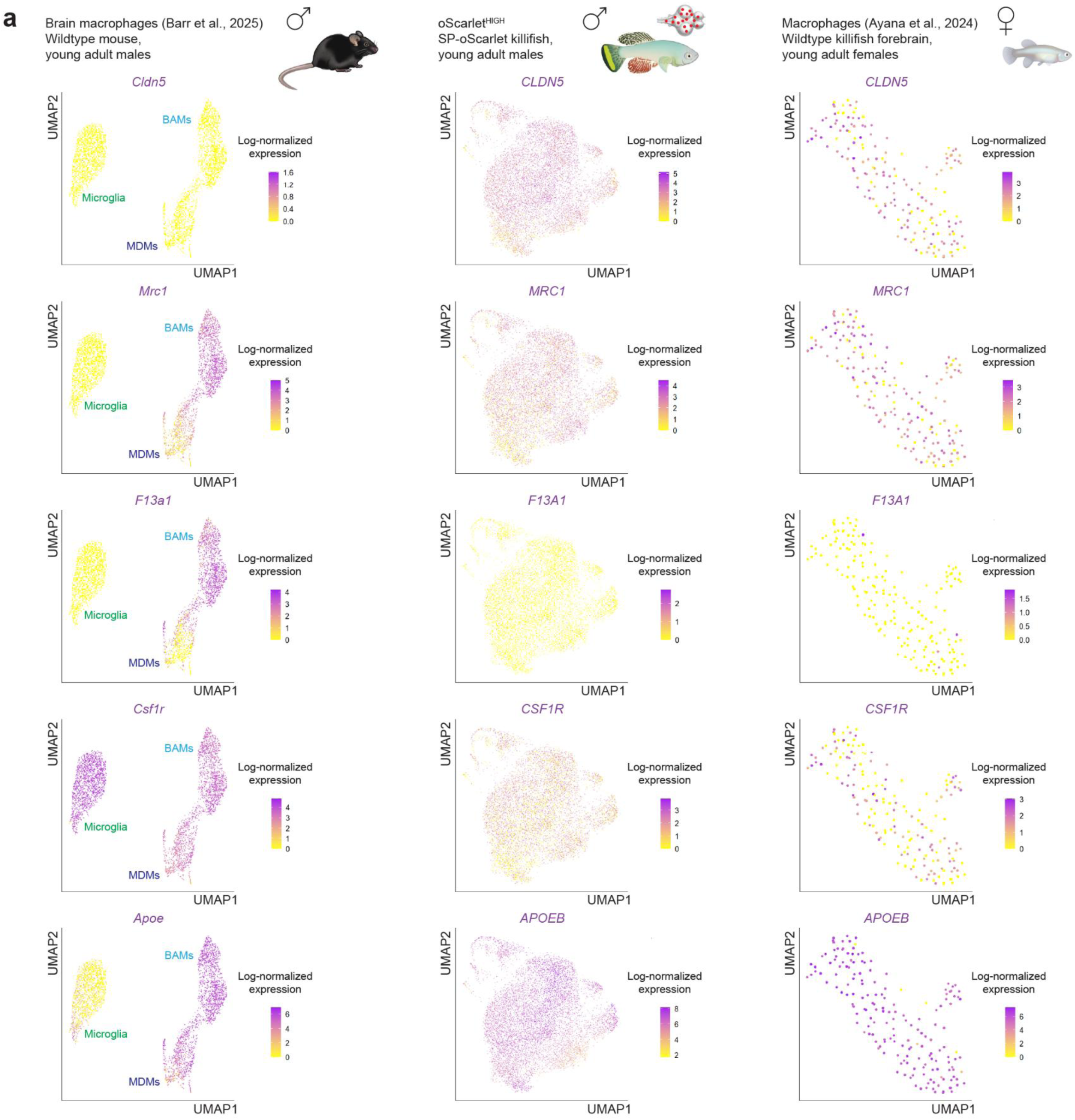
**a,** UMAPs of wildtype young adult mouse brain macrophages (Barr et al., 2025^75^), young adult oScarlet^HIGH^ cells from experiment in **1d**, and wildtype young adult killifish forebrain macrophages (Ayana et al., 2024^68^) colored according to log-normalized *Cldn5* (or killifish homolog *CLDN5* (*LOC107394244*)), *Mrc1* (or killifish homolog *MRC1* (*LOC107383768*)), *F13a1* (or killifish homolog *F13A1* (*LOC107377057*)), *Csf1r* (or killifish homolog *CSF1R* (*LOC107381415*)), or *Apoe* (or killifish homolog *APOEB* (*LOC107379395*)) expression. Each dot represents one cell.

**Figure 2 – Figure Supplement 3.**
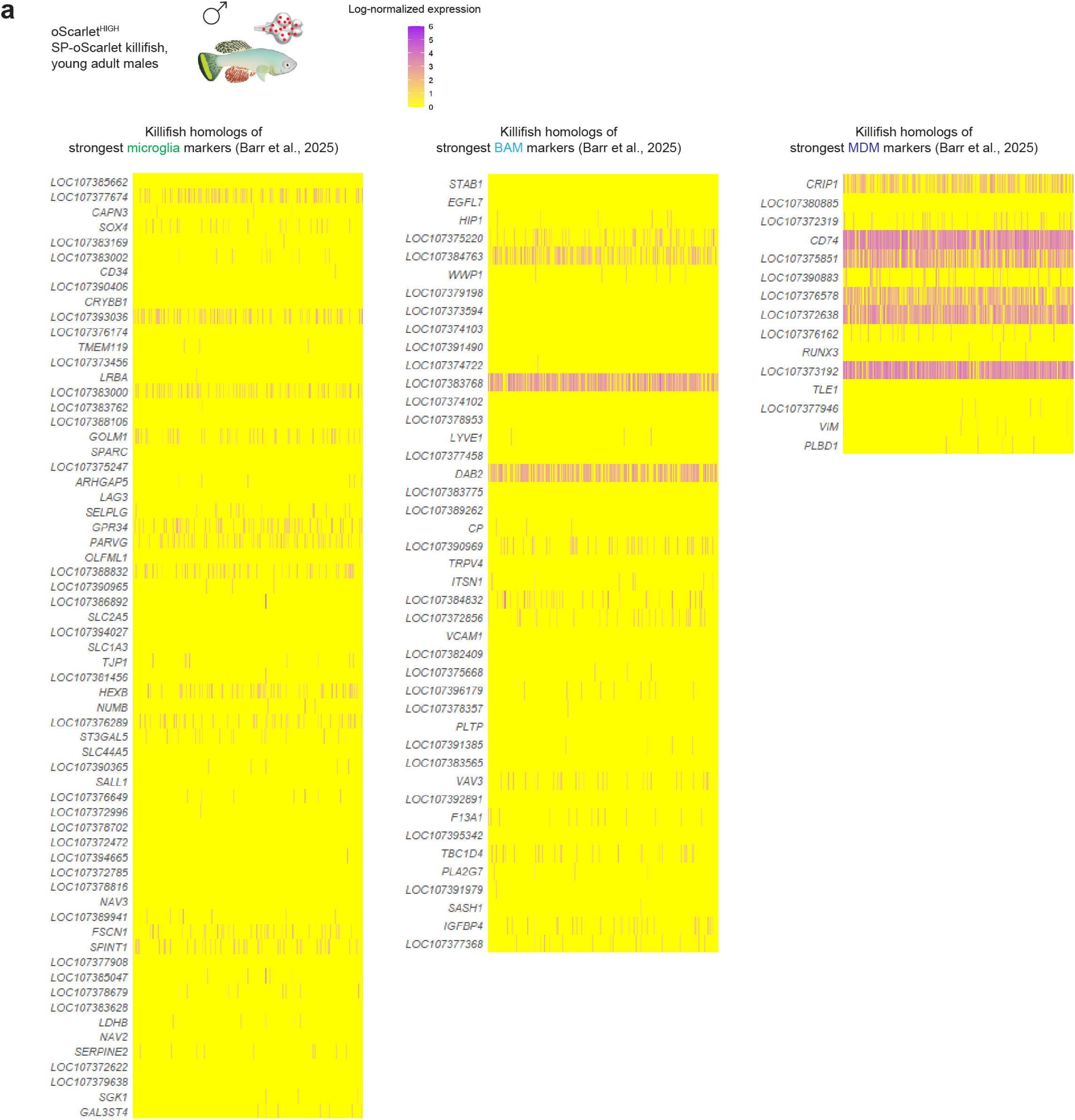
**a,** Heatmaps of expression of killifish homologs of strongest microglia, BAM, and MDM marker genes (Barr et al., 2025^75^) in young adult oScarlet^HIGH^ cells from experiment in **1d**.

**Figure 2 – Figure Supplement 4.**
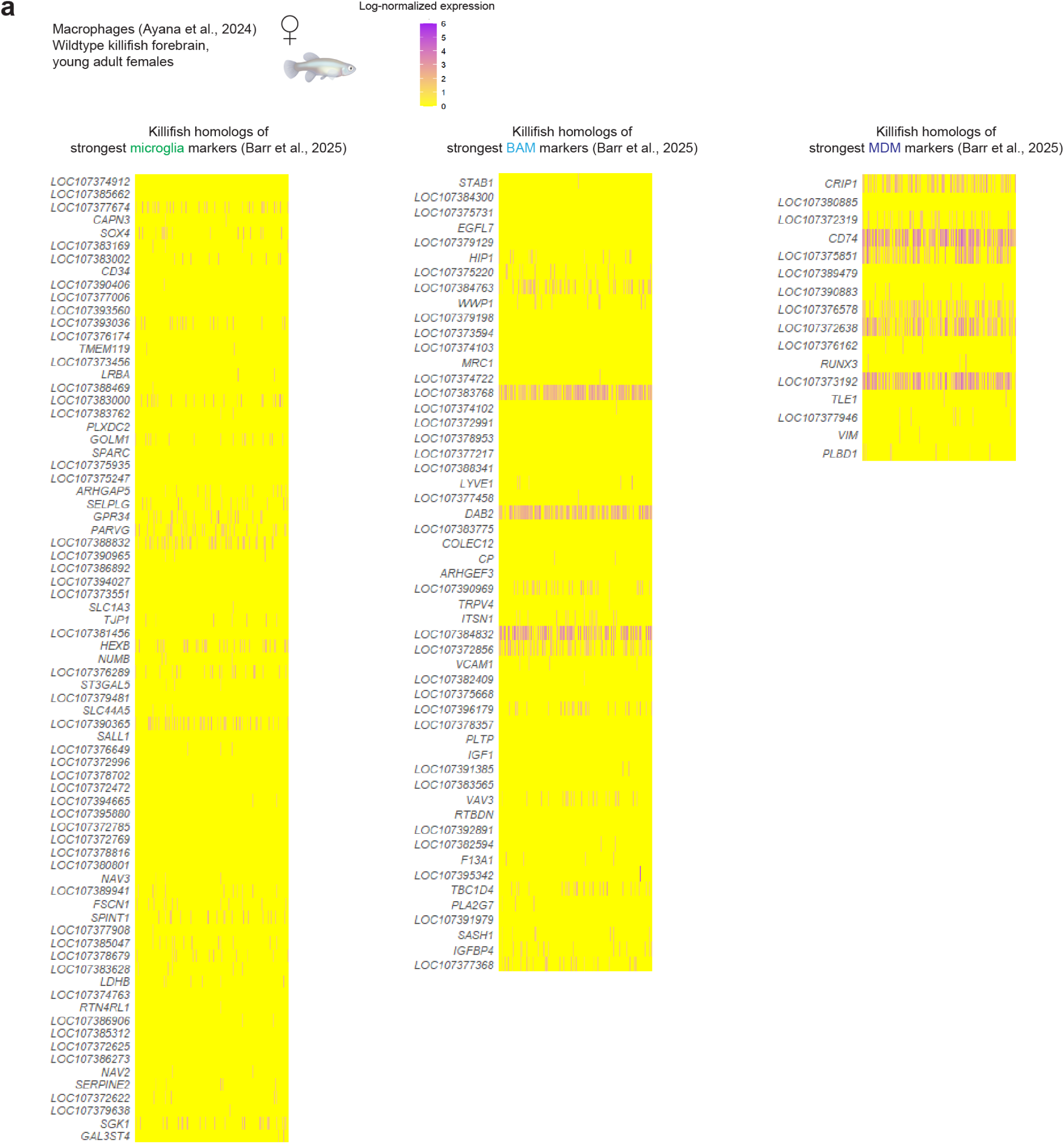
**a,** Heatmaps of expression of killifish homologs of strongest microglia, BAM, and MDM marker genes (Barr et al., 2025^75^) in wildtype young adult killifish forebrain macrophages (Ayana et al., 2024^68^).

**Figure 3 – Figure Supplement 1.**
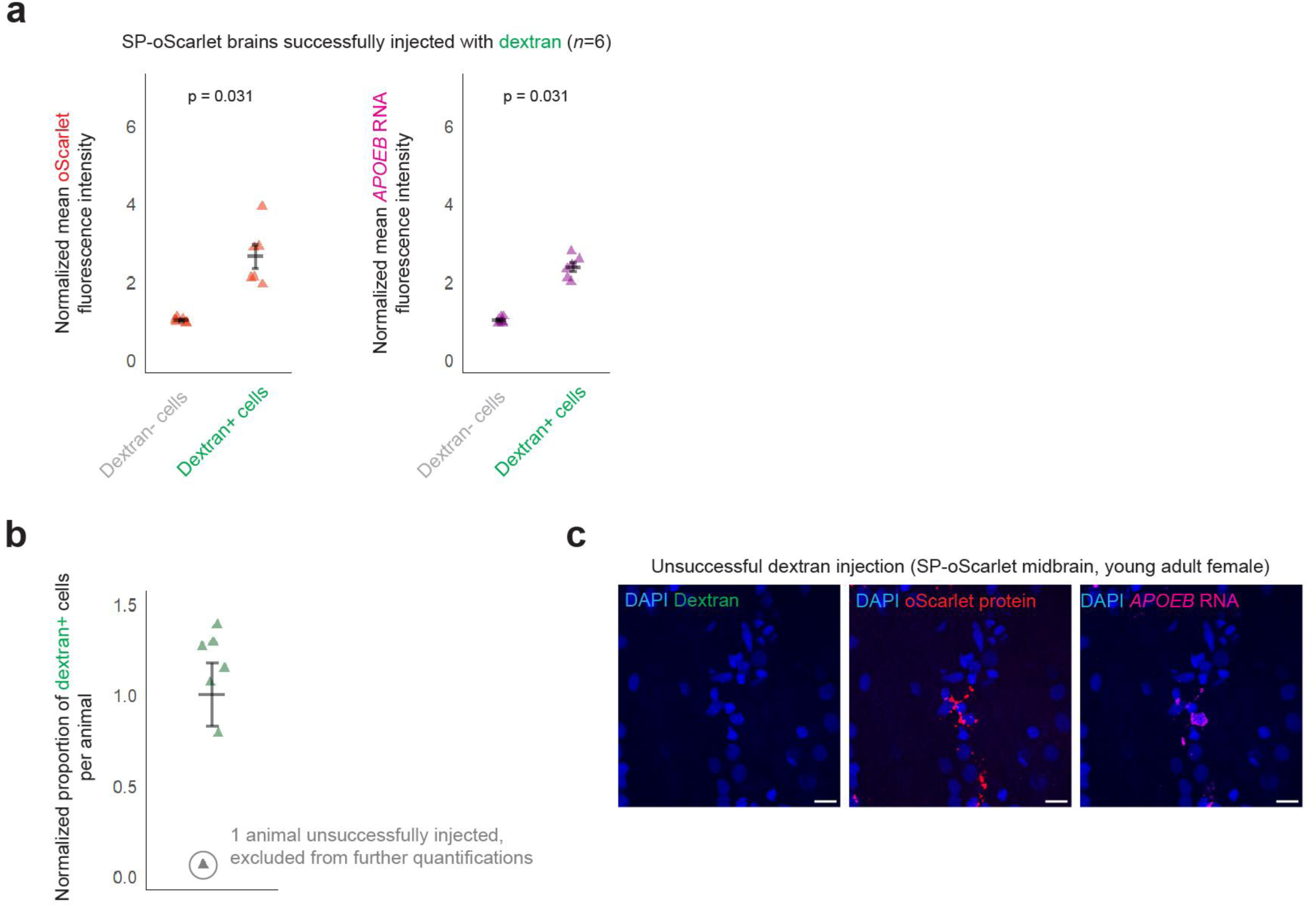
**a,** Quantification of mean oScarlet fluorescence intensity and mean *APOEB* RNA fluorescence intensity in dextran-negative and dextran-positive cells (n = 6 successfully injected fish, 63-91 days females, with 5-6 regions of interest over 1-2 sagittal brain sections per fish, over two experiments). Four images (one image per fish for each of four fish) of regions of interest with a visually apparent very high density of cells with high *APOEB* RNA expression, which could plausibly reflect an injection-related injury, were excluded from quantifications. The excluded images are listed in **Supplementary Table 4**. Each triangle represents one fish. Mean +/-standard error of mean; p-values, paired Wilcoxon rank-sum test at per-animal level. **b,** Quantification from experiment in **3a** of proportion of dextran-positive cells per fish (n = 7 fish, 63-91 days females, with 5-6 regions of interest over 1-2 sagittal brain sections per fish, over two experiments), highlighting in grey one fish in which dextran was not observed to circulate throughout the brain (also confirmed by visual inspection of sections). This unsuccessfully injected fish (63 days, female) was excluded from subsequent quantifications. Each triangle represents one fish. **c,** Representative image of brain sections from the young adult (63 days) female heterozygous SP-oScarlet killifish unsuccessfully injected with dextran (n = 1 fish). Scale bars = 10 μm.

**Figure 4 – Figure Supplement 1.**
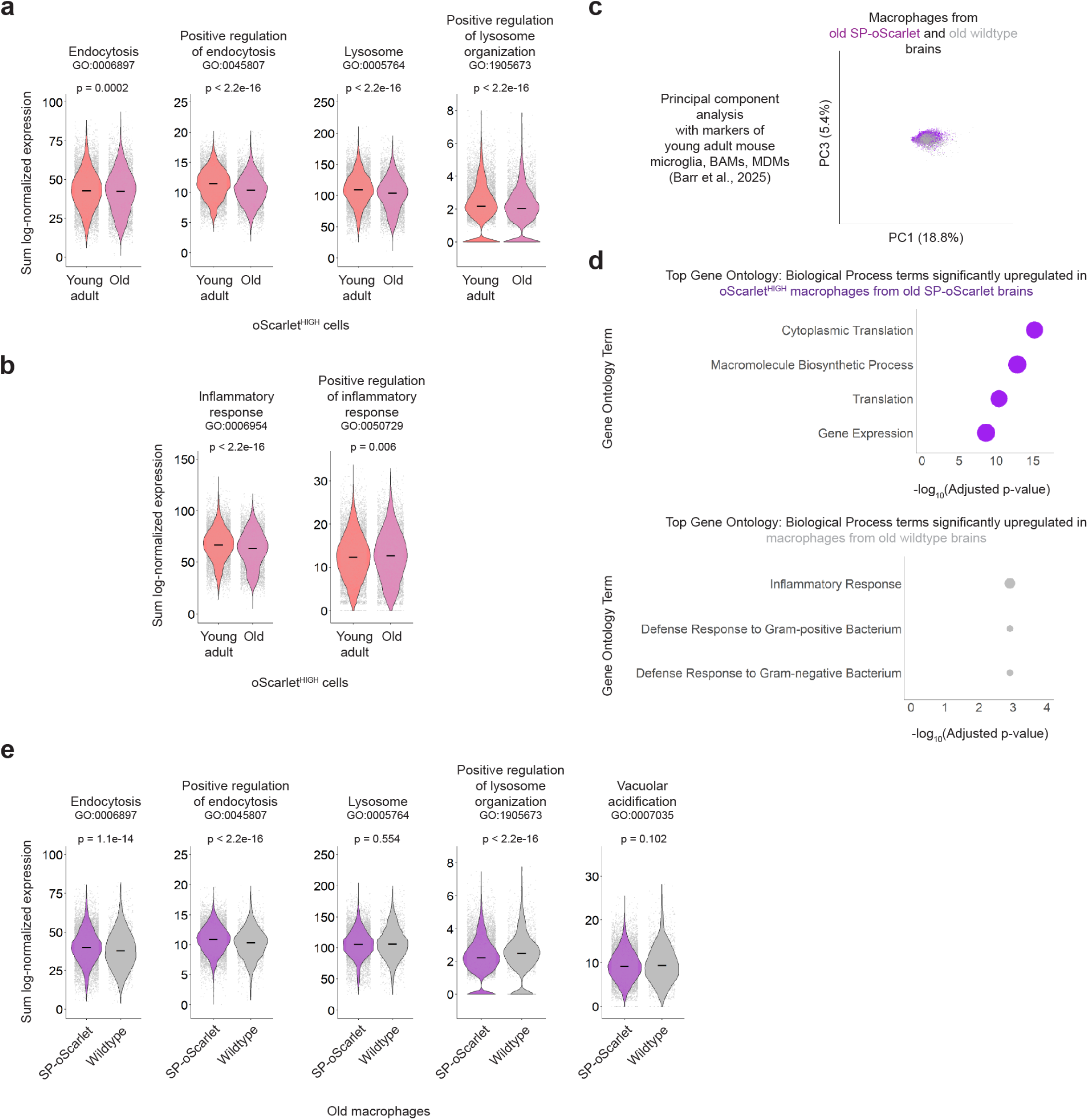
**a,** Violin plots for sum log-normalized expression of genes in Gene Ontology: Biological Process terms “endocytosis” (GO:0006897), “positive regulation of endocytosis” (GO:0045807), “lysosome” (GO:0005764), and “positive regulation of lysosome organization” (GO:1905673) in young adult and old oScarlet^HIGH^ cells from experiment in **1d**. Each dot represents one cell. Lines: medians; p-values, unpaired Wilcoxon rank-sum test at per-cell level (fish were pooled for single cell RNA-sequencing experiments to obtain sufficient numbers of oScarlet^HIGH^ cells). **b**, Violin plots for sum log-normalized expression of genes in Gene Ontology: Biological Process terms “inflammatory response” (GO:0006954) and “positive regulation of inflammatory response” (GO:0050729) in young adult and old oScarlet^HIGH^ cells from experiment in **1d**. Each dot represents one cell. Lines: medians; p-values, unpaired Wilcoxon rank-sum test at per-cell level. **c,** PCA plot (principal components 1 vs. 3) of oScarlet^HIGH^ and wildtype old killifish brain macrophages from experiment in **4e** using all killifish homologs of marker genes of microglia, BAMs, and MDMs (Barr et al., 2025^75^). Each dot represents one cell. Percentages on axes represent the percentage of variance explained by each respective principal component. **d,** Top Gene Ontology: Biological Process terms upregulated in old oScarlet^HIGH^ macrophages (above) and in old wildtype macrophages (below) from experiment in **4e**, as ranked by adjusted p-value (Fisher’s exact test, Benjamini-Hochberg FDR-corrected). Size of each circle corresponds to number of distinct human homologs among differentially expressed genes from each term. Full lists of differentially expressed genes and upregulated terms in **Supplementary Tables 12-14**. **e,** Violin plots for sum log-normalized expression of genes in Gene Ontology: Biological Process terms “endocytosis” (GO:0006897), “positive regulation of endocytosis” (GO:0045807), “lysosome” (GO:0005764), “positive regulation of lysosome organization” (GO:1905673), and “vacuolar acidification” (GO: 0007035) in old oScarlet^HIGH^ and old wildtype macrophages from experiment in **4e**. Each dot represents one cell. Lines: medians; p-values, unpaired Wilcoxon rank-sum test at per-cell level.

**Figure 5 – Figure Supplement 1.**
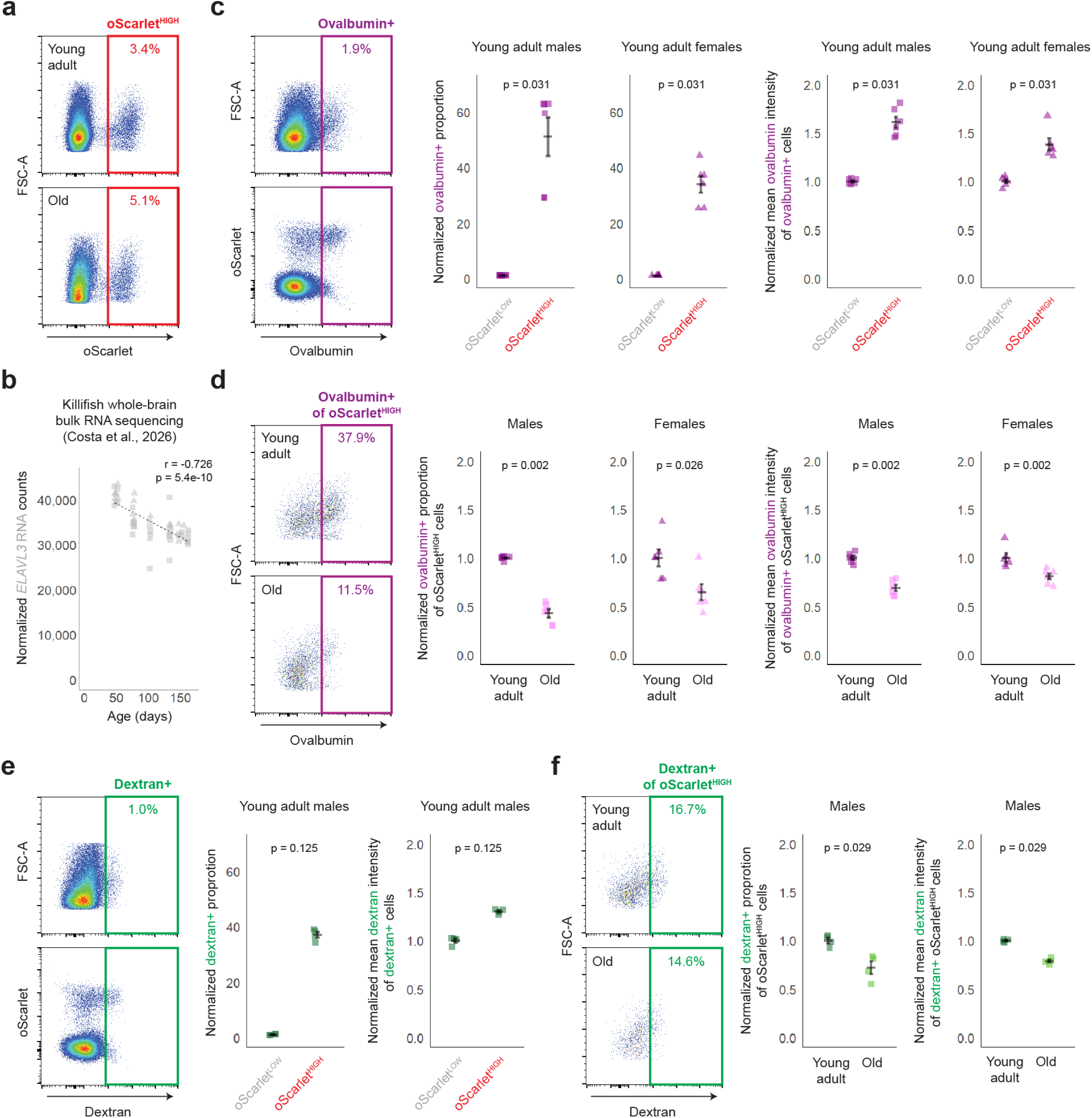
**a,** Representative flow cytometry plots from young adult (41 days, above) and old (160 days, below) male heterozygous SP-oScarlet killifish brains, highlighting oScarlet^HIGH^ cells. Each dot represents one cell. **b,** Plot of normalized *ELAVL3* (*LOC107378364*) RNA counts across age from bulk RNA sequencing of whole killifish brains (Costa et al., 2026^55^). Each square (male) or triangle (female) represents one fish. Dashed line: linear regression. r: Pearson correlation coefficient; p-value, two-sided t-test. **c,** (Left) Representative flow cytometry plots from young adult (41 days) male heterozygous SP-oScarlet killifish brains, highlighting ovalbumin+ cells as identified from all Live cells, with mean ovalbumin fluorescence intensity plotted against either FSC-A (above) or mean oScarlet fluorescence intensity (below). Each dot represents one cell. (Right) Quantifications of proportion of ovalbumin+ cells and mean ovalbumin fluorescence intensity of ovalbumin+ cells among young adult oScarlet^LOW^ and oScarlet^HIGH^ cells (same experiments as in **5a**). Each square (male) or triangle (female) represents one fish. Mean +/-standard error of mean; p-values, paired Wilcoxon rank-sum test at per-animal level. **d,** (Left) Representative flow cytometry plots from young adult (41 days) (above) and old (160 days) (below) male heterozygous SP-oScarlet killifish brains, highlighting ovalbumin+ oScarlet^HIGH^ cells. Each dot represents one cell. (Right) Quantifications of proportion of ovalbumin+ cells and mean ovalbumin fluorescence intensity of ovalbumin+ cells among young adult and old oScarlet^HIGH^ cells (same experiments as in **5a**). Each square (male) or triangle (female) represents one fish. Mean +/- standard error of mean; p-values, unpaired Wilcoxon rank-sum test at per-animal level. **e,** (Left) Representative flow cytometry plots from young adult (56 days) male heterozygous SP-oScarlet killifish brains, highlighting dextran+ cells as identified from all Live cells, with mean dextran fluorescence intensity plotted against either FSC-A (above) or mean oScarlet fluorescence intensity (below). Each dot represents one cell. (Right) Quantifications of proportion of dextran+ cells and mean dextran fluorescence intensity of dextran+ cells among young adult oScarlet^LOW^ and oScarlet^HIGH^ cells (same experiment as in **5e**). Each square represents one fish. Mean +/- standard error of mean; p-values, paired Wilcoxon rank-sum test at per-animal level. **f,** (Left) Representative flow cytometry plots from young adult (56 days) (above) and old (156 days) (below) male heterozygous SP-oScarlet killifish brains, highlighting dextran+ oScarlet^HIGH^ cells. Each dot represents one cell. (Right) Quantifications of proportion of dextran+ cells and mean dextran fluorescence intensity of dextran+ cells among young adult and old oScarlet^HIGH^ cells (same experiment as in **5e**). Each square represents one fish. Mean +/- standard error of mean; p-values, unpaired Wilcoxon ranksum test at per-animal level.

**Figure 5 – Figure Supplement 2.**
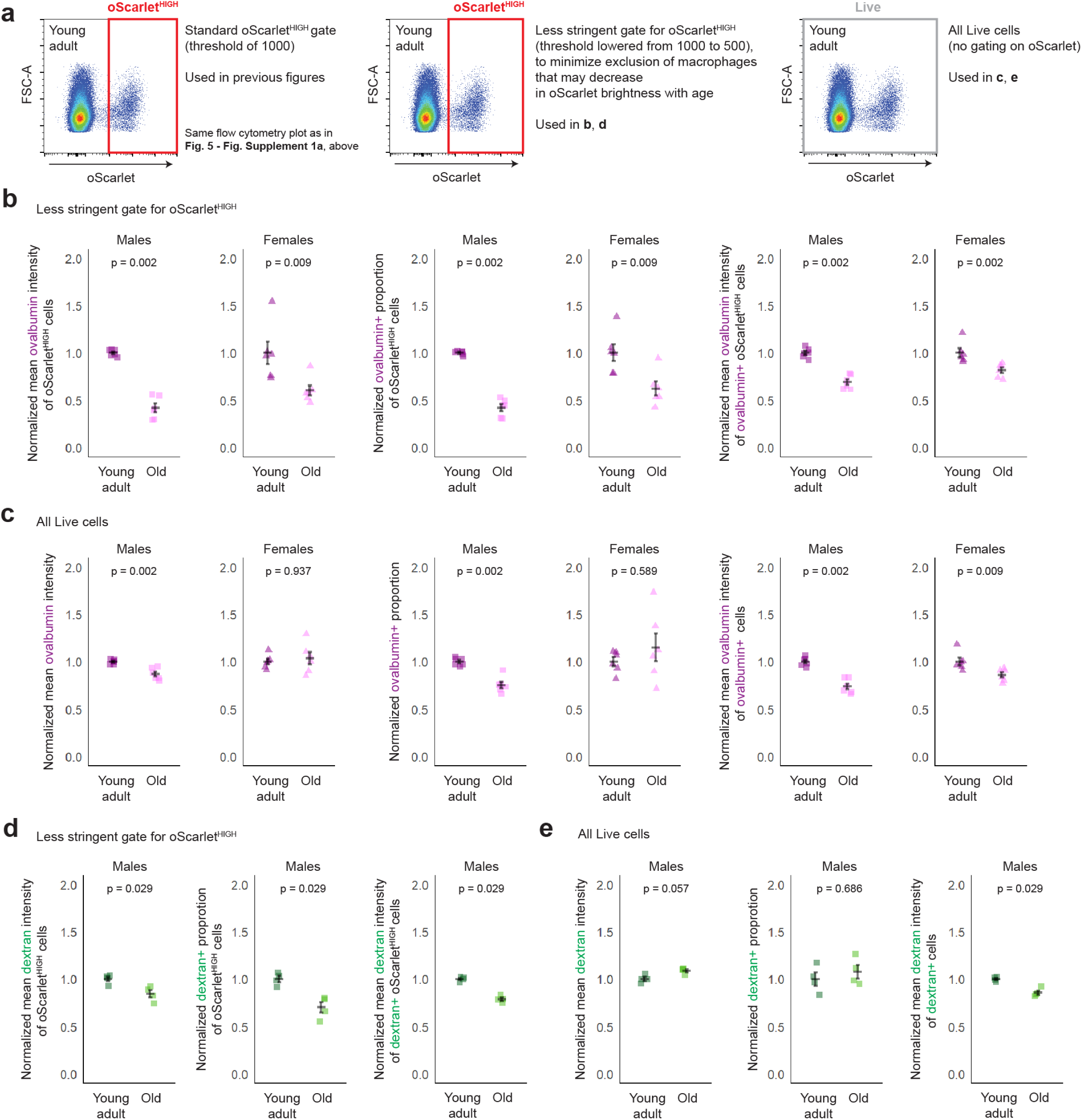
**a,** Same flow cytometry plot as in **5S1a** (above), illustrating alternative gating schemes used in subsequent panels. **b,** Quantifications of mean ovalbumin fluorescence intensity, proportion of ovalbumin+ cells, and mean ovalbumin fluorescence intensity of ovalbumin+ cells among young adult and old oScarlet^HIGH^ cells (same experiments as in **5a**) with a less stringent oScarlet^HIGH^ gate (lowered from 1000 to 500). Each square (male) or triangle (female) represents one fish. Mean +/-standard error of mean; p-values, unpaired Wilcoxon rank-sum test at per-animal level. **c,** Quantifications of mean ovalbumin fluorescence intensity, proportion of ovalbumin+ cells, and mean ovalbumin fluorescence intensity of ovalbumin+ cells among all young adult and old cells (same experiments as in **5a**, no gating for oScarlet). Each square (male) or triangle (female) represents one fish. Mean +/- standard error of mean; p-values, unpaired Wilcoxon rank-sum test at per-animal level. **d,** Quantifications of mean dextran fluorescence intensity, proportion of dextran+ cells, and mean dextran fluorescence intensity of dextran+ cells among young adult and old oScarlet^HIGH^ cells (same experiment as in **5e**) with a less stringent oScarlet^HIGH^ gate (lowered from 1000 to 500). Each square represents one fish. Mean +/- standard error of mean; p-values, unpaired Wilcoxon rank-sum test at per-animal level. **e,** Quantifications of mean dextran fluorescence intensity, proportion of dextran+ cells, and mean dextran fluorescence intensity of dextran+ cells among all young adult and old Live cells (same experiment as in **5e**, no gating for oScarlet). Each square represents one fish. Mean +/-standard error of mean; p-values, unpaired Wilcoxon rank-sum test at per-animal level.

## Supplementary Tables

**Supplementary Table 1.** Animals used in this study.

**Supplementary Table 2.** Nucleic acid sequences used in this study.

**Supplementary Table 3.** Antibodies used in this study.

**Supplementary Table 4.** Images taken in this study.

**Supplementary Table 5.** Gene names conversion table used to identify homologs across species in this study.

**Supplementary Table 6.** Quality control metrics for Seurat objects generated from raw reads of killifish single-cell RNA sequencing data in this study.

**Supplementary Table 7.** Cluster marker genes based on all young adult cells from August 2025 single-cell RNA sequencing experiment.

**Supplementary Table 8.** Cell type marker genes based on all young adult cells from August 2025 single-cell RNA sequencing experiment.

**Supplementary Table 9.** Differentially expressed genes between young adult and old oScarlet^HIGH^ cells from August 2025 single-cell RNA sequencing experiment. avg_log2FC > 0 indicates enrichment in old oScarlet^HIGH^ cells. avg_log2FC < 0 indicates enrichment in young adult oScarlet^HIGH^ cells. pct. 1 corresponds to the percentage of old oScarlet^HIGH^ cells with detectable expression of the gene. pct. 2 corresponds to the percentage of young adult oScarlet^HIGH^ cells with detectable expression of the gene.

**Supplementary Table 10.** Gene Ontology: Biological Process (2025) terms upregulated in young adult oScarlet^HIGH^ cells from August 2025 single-cell RNA sequencing experiment.

**Supplementary Table 11.** Gene Ontology: Biological Process (2025) terms upregulated in old oScarlet^HIGH^ cells from August 2025 single-cell RNA sequencing experiment.

**Supplementary Table 12.** Differentially expressed genes between old oScarlet^HIGH^ and old wildtype macrophages from October 2024 single-cell RNA sequencing experiment. avg_log2FC > 0 indicates enrichment in old oScarlet^HIGH^ macrophages. avg_log2FC < 0 indicates enrichment in old wildtype macrophages. pct. 1 corresponds to the percentage of old oScarlet^HIGH^ macrophages with detectable expression of the gene. pct. 2 corresponds to the percentage of old wildtype macrophages with detectable expression of the gene.

**Supplementary Table 13.** Gene Ontology: Biological Process (2025) terms upregulated in old oScarlet^HIGH^ macrophages from October 2024 single-cell RNA sequencing experiment.

**Supplementary Table 14.** Gene Ontology: Biological Process (2025) terms upregulated in old wildtype macrophages from October 2024 single-cell RNA sequencing experiment.

## Source Data Tables

**Source Data Table 1**. Corresponding to Fig. 3a.

**Source Data Table 2**. Corresponding to **Fig. 3 – Supplementary** Fig. 1a.

**Source Data Table 3**. Corresponding to **Fig. 3 – Supplementary** Fig. 1b.

**Source Data Table 4**. Supporting information for analysis in Fig. 3b.

**Source Data Table 5**. Supporting information for analysis in Fig. 3b.

**Source Data Table 6**. Corresponding to Fig. 3b.

**Source Data Table 7**. Corresponding to Fig. 5a, Fig. 5c**-d**, and **Fig. 5 – Fig. Supplement 1c-d**.

**Source Data Table 8**. Corresponding to **Fig. 5 – Fig. Supplement 2b**.

**Source Data Table 9**. Corresponding to **Fig. 5 – Fig. Supplement 2c**.

For this analysis, the threshold for oScarlet^HIGH^ was set to 0.

**Source Data Table 10**. Corresponding to Fig. 5e**-f** and **Fig. 5 – Fig. Supplement 1e-f**.

**Source Data Table 11**. Corresponding to **Fig. 5 – Fig. Supplement 2d**.

**Source Data Table 12**. Corresponding to **Fig. 5 – Fig. Supplement 2e**.

For this analysis, the threshold for oScarlet^HIGH^ was set to 0.

## Additional Supplements

**Supplement 1.** SnapGene map file of expected SP-oScarlet insertion.

**Supplement 2.** Compensation matrix from first *ex vivo* ovalbumin engulfment experiment.

**Supplement 3.** Compensation matrix from second *ex vivo* ovalbumin engulfment experiment.

**Supplement 4.** Compensation matrix from *ex vivo* dextran engulfment experiment.

## Methods

### Killifish husbandry

The short-lived *Nothobranchius furzeri* GRZ strain^38^ was used as the background killifish strain in this study. Fish were maintained using an Aquaneering recirculating aquatic system at a temperature of ∼27°C, pH of 7-7.5, conductivity of 1200 μS with a 12 hour/12 hour light/dark cycle with the light on between ∼7 a.m. and ∼7 p.m. For the first 1 month of life, fish were fed live brine shrimp (Brine Shrimp Direct, BSEP6LB). Brine shrimp feeding solution was generated by mixing 50 mL of brine eggs hatched over 48 hours at ∼27°C in a cone with 10 L reverse osmosis (RO) water and 1.5 cups salt (approximately 350 mL; Instant Ocean; Amazon, B000256EUS)). Fish were fed brine shrimp twice daily at ∼9 a.m. and ∼3 p.m. on weekdays (0.5-2 mL per feeding, with volume per feeding approximately scaling with each week of development). Fish were fed once daily on weekends and holidays. After 1 month, fish were fed food pellets (Reed Mariculture, Otohime C1) twice daily at ∼9 a.m. and ∼3 p.m. (and once daily on weekends and holidays) with ∼18 mg per feeding. Tests for pathogens were conducted every three months (Charles River Labs). Fish were housed in Stanford ChEM-H/Neuro Vivarium and Research Animal Facility (RAF), according to IACUC protocol 13645.

For breeding and collecting killifish embryos, plastic weigh boats (e.g., Heathrow Scientific, HS1420BB) filled up to halfway with sand were dropped in breeding tanks containing 1 male and 1-10 females (all 1-4 months old, 1.5-2.5 month old breeding females used when available; 2.8 L breeding tank for up to 3 females, 6.8 L breeding tank for up to 6 females; 9.8 L breeding tank for up to 10 females) to initiate mating, and approximately 3-6 hours later the embryos were collected by pouring the contents of the sand tray through a sieve.

Collected embryos were placed in 5-15 mL embryo solution liquid (embryo solution was made by adding 2 Ringer tablets (EMD Millipore, 1.15525.0001) and 100 μL methylene blue (Kordon, 37344) to 1 L Milli-Q water, for a final concentration of 0.01% methylene blue) in 35 or 60 mm diameter Petri dishes (Greiner, 627102 and 628161 respectively; dish size used dependent on number of embryos, with a maximum of ∼40 embryos in a 35 mm and ∼150 embryos in a 60 mm Petri dish) for 2 weeks at 28°C in a dark incubator (e.g., Thermo 50125590H). Dead embryos were removed each day, with the embryo solution exchanged on the first day and on an as-needed basis thereafter. Embryos were generally treated with Povidone Iodine Prep Solution U.S.P. (Dynarex; Amazon, B00859URLK) (added to the Petri dish with embryo solution at ∼1:1000 dilution, followed by 2-3 washes with embryo solution) immediately after collection.

At 14-16 days post fertilization (dpf), embryos were placed on autoclaved coconut fiber (Fluker’s; Amazon, B01HO27S4M) in 35 or 60 mm diameter Petri dishes (with a maximum of ∼40 embryos in a 35 mm and ∼150 embryos in a 60 mm Petri dish) and maintained at 27-30°C for an additional 2 weeks before hatching by placing in ∼5 mL cold (4°C) humic acid solution (Sigma-Aldrich, 53680, 1 g/L in RO water) in a 35 mm Petri dish left overnight at room temperature, followed by transfer to the aquatic system. In certain cases (noted in **Supplementary Table 1**), some embryos entered diapause, even at 28°C (this did not appear to be linked to maternal age or any other obviously noticeable parameter), and had to be induced out of diapause using a thermocycler program of 6 hours at 28°C and 6 hours at 37°C repeated 6 times before being allowed to continue to develop at 28°C and transferred to coconut fiber followed by hatching in cold humic acid.

Newly hatched killifish (fry) were housed on the system at a density of up to 20 fry per 0.8 L tank, with successive reductions in density as the fish grew older until reaching a density of 1 male fish per 1.8-2.8 L tank, or 1-10 female fish per 1.8-9.8 L tank (2.8 L tank for up to 4 females, 6.8 L breeding tank for up to 6 females; 9.8 L breeding tank for up to 10 females), beyond 1 month of age (sexual maturity).

Unless otherwise noted, original data in this study from all figures were collected from animals raised in Stanford ChEM-H/Neuro Vivarium after a general system bleaching (January 2025) and treated as described above. Data from Fig. 1b, Fig. 1c, Fig. 1h (images), **Fig. 1 – Fig. Supplement 1c**, **Fig. 1 – Fig. Supplement 1d** (4 of 10 fish), and Fig. 3b were obtained in another facility at Stanford (RAF), with specific husbandry variations noted in **Supplementary Table 1** (see “Notes”). Data from **Fig. 1 – Fig. Supplement 1a**, **Fig. 1 – Fig. Supplement 1f**, Fig. 4f**-i**, and **Fig. 4 – Fig. Supplement 1c-e** were collected in ChEM-H/Neuro Vivarium before system bleaching (with bryozoans and mycobacteria prevalent), with specific husbandry variations noted in **Supplementary Table 1** (see “Notes”). A list of animals used in each experiment/figure panel, with information on age (with hatch and sacrifice dates), notes on husbandry conditions, sex, and genotype, can be found in **Supplementary Table 1**. Importantly, the key phenotypes observed in this study were robust to variations in husbandry conditions.

### Generation of SP-oScarlet killifish

To generate a killifish knockin strain expressing a secreted form of the red fluorescent protein oScarlet in the nervous system, we used our CRISPR-based knockin method^67^. Overall, a construct encoding a signal peptide (SP) from killifish HSPA5 (signal peptide predicted using SignalP 5.0^71^) fused to the N-terminus of the red fluorescent protein oScarlet^70^ was inserted by homology-directed repair just 5’ of the endogenous stop codon of the *ELAVL3* (*LOC107378364*) locus in the killifish GRZ wildtype strain. Across multiple vertebrate species, *ELAVL3* encodes an RNA-binding protein expressed strongly in neurons, as well as some glia such as oligodendrocytes^68,78^, and has successfully be used in killifish as a driver for pan-neuronal expression^67^.

The signal peptide allows the secretion of oScarlet from ELAVL3-expressing cells. To prevent unwanted fusion of oScarlet to ELAVL3, the construct also encodes a P2A self-cleaving peptide^72^ at the C-terminal end of ELAVL3 (full sequence of construct, including 200 bp homology arms at either end, in **Supplementary Table 2**; annotated SnapGene map file in **Supplement 1**).

To generate the killifish strain expressing this secreted form of oScarlet, which we refer to as SP-oScarlet, we followed the step-by-step CRISPR knockin protocol described by Bedbrook et al., 2023^67^. For each round of injection attempted, a sand tray was dropped in a harem breeding tank (6.8-9.8L tank) containing 1 GRZ wildtype male and 5-10 GRZ wildtype females (all 1-4 months old, with 1.5-2.5 months old breeders used when available) to initiate mating, and approximately 3-4 hours later the embryos were collected by pouring the contents of the sand tray through a sieve.

Injection needles were generated from borosilicate glass capillaries (Sutter, BF100-58-10) pulled on P-97 pipette pullers (Sutter Instruments). A ramp test was performed and a Heat value of [RAMP value + 30] was used. Other settings used were Pull = 20-30, Velocity = 19-30, Time = 200. These parameters were optimized on each specific pipette puller used, with the resulting needles tested in pilot injections.

In the 30 minutes before collection, 1 μL of a crRNA targeting the 3’ end of the coding sequence of *ELAVL3* (at 100 μM in Nuclease-Free Duplex Buffer (IDT, 11-01-03-01); sequence in **Supplementary Table 2**) and 1 μL of tracrRNA (at 100 μM in Nuclease-Free Duplex Buffer; IDT, 1072532/3/4) were added to 31.3 μL Nuclease-Free Duplex Buffer and annealed at 95°C for 5 minutes to generate the guide RNA (gRNA) solution. After this annealing, 10 μL of the resultant gRNA solution was added to 0.5 μL rCas9 protein (10 μg/μL; IDT, 1081059), and 5.5 μL PBS (Corning, 21-040-CV) and this mixture was heated at 37°C for 10 minutes to generate the ribonucleoprotein complex. After this heating, 8 μL of the ribonucleoprotein complex was added to 1 μL dsDNA homology-directed repair (HDR) template (150 ng/μL; IDT custom-synthesized, sequence in **Supplementary Table 2**, annotated SnapGene map file in **Supplement 1**), 1 μL HDR enhancer solution (IDT 10007921; 10 μM), and 0.33 μL phenol red (e.g., Sigma-Aldrich P0290-100ML) to generate the injection solution.

For injection, embryos at the 1 cell stage were aligned in rows on an injection pad (1% agarose in embryo solution, solidified over a custom-3D-printed mold with rows) with the single cell facing upwards placed under a dissecting microscope (Leica, MZ6). Embryos were individually injected with injection solution dispensed through the aforementioned needles (loaded using Microloader (Calibre Scientific, EPE-5242956003) tips; per-injection volume estimated to be in the nanoliter range). Needles were unsealed by brushing against sides of embryos. Injections were performed with a MPPI-2 Pressure Injector (ASI) with Air Out = Continuous, Pressure ∼ 10 psi. This procedure allowed for a constant, slow stream of injection solution flowing out of the needle (visible due to the use of phenol red); the needle was inserted into the single cell of an embryo and injection solution allowed to flow out until the single cell was visibly red (∼2-3 seconds).

After injection, embryos were placed in embryo solution in 35 or 60 mm diameter Petri dishes for 2 weeks at 28°C and dead embryos were removed daily. At 9-11 dpf, we identified F0 fish expressing the red fluorescent protein oScarlet in the nervous system (by visual inspection of embryos under a fluorescent dissecting microscope). At 14-16 dpf, embryos were placed on coconut fiber in 35 or 60 mm diameter Petri dishes and maintained at 27-30°C for an additional 2 weeks before hatching by placing in cold humic acid solution left at room temperature overnight (see above), followed by transfer to the aquatic system.

To generate a stable SP-oScarlet line, adult F0 fish (as determined by the visual screening of embryos described above) were crossed with GRZ wildtype fish. F1 fish and some subsequent generations were genotyped to identify SP-oScarlet progeny (see “Genotyping,” below). We outcrossed F1 SP-oScarlet fish with wildtype GRZ fish for >5 generations. Both

GRZ male x SP-oScarlet female and SP-oScarlet male x GRZ female crosses were used. In one cohort, we generated homozygous SP-oScarlet fish by crossing heterozygous parents and noted the homozygotes had relatively poor survival at the fry stage (<10% surviving to one month of age). Therefore, we maintained the line at the heterozygous state and we used heterozygotes for all experiments.

### Genotyping

In most generations, SP-oScarlet progeny of SP-oScarlet x GRZ wildtype crosses were identified by visual inspection of embryos. Nonetheless, in the F1 generation and some subsequent generations, we used PCR genotyping to confirm the presence of an insertion at the *ELAVL3* locus, using a three-primer PCR strategy with e.g., primers pFL1, pRL1, and pmCF (sequences in **Supplementary Table 2**). All primers used were synthesized by IDT.

Genomic DNA was extracted from a small (<1 mg) amount of tail tissue cut from a live fish, either from a 1 day old fry or an adult fish, anesthetized either on a dish placed over ice (for fry) or in 300 mg/L Tricaine (dissolved in system water; e.g., Pentair, TRS1) buffered with sodium bicarbonate to pH ∼7 (for adults).

The tail tissue was lysed in 30 μL (for fry) or 100 μL (for adults) 2% (volume/volume) Proteinase K (e.g., Invitrogen, 25-530-049) in DirectPCR lysis solution (Viagen Biotech, 102-T) in a PCR tube (e.g., USA Scientific, 1402-4708) using a thermocycler program of 55°C for 2 hours (lysis) and 80°C for 10 minutes (Proteinase K inactivation).

Next, 2 μL (for fry) or 3 μL (for adults) of the resulting lysate (unpurified) was added to 10 μL 2x GoTaq Green Master Mix (Promega, M7123), 6 μL Milli-Q water, and 0.5 μL each of primers pFL1, pRL1, and pmCF (each at 10 μM in Milli-Q water) following the manufacturer’s instructions for PCR with a total reaction volume of 19-20 μL and a thermocyler program of 98°C (pre-heating); 98°C for 30 seconds; 30 cycles of 98°C for 30 seconds, 55°C for 1 minute, and 72°C for 90 seconds; and finally 72°C for 5 minutes.

Then, 8 μL PCR product was resolved on a 1% agarose gel to verify the size of amplified bands. Samples with only one band of size 971 bp (pFL1, pRL1 band) were identified as wildtype, while samples with two bands (824 bp; pmCF, pRL1 band from knockin allele and 971 bp; pFL1, pRL1 band from wildtype allele) were identified as heterozygous SP-oScarlet (the >1.5 kb pFL1, pRL1 band from the knockin allele would not be expected to be amplified efficiently under these PCR conditions).

Separately, to validate the sequence of the genomic insertion in a SP-oScarlet heterozygous fish, we performed PCR with primers pFL1 and pRL1 outside the homology arms of the knockin construct. For this, genomic DNA was extracted from the sidebody muscle of heterozygous SP-oScarlet killifish (78 days old male). The fish was sacrificed in ice water and <25 mg of sidebody muscle was cut out with a scalpel (Integra, 4-122), chopped, and placed in 200 μL of DNEasy Blood & Tissue kit (Qiagen, 69504/6) Buffer AL in a 1.5-mL tube (e.g., Eppendorf, 0030108418). The DNEasy Blood & Tissue kit manufacturer’s protocol was then followed to extract DNA, which was eluted in 200 μL of Buffer AE.

DNA amount and quality was assessed by NanoDrop and 1 μL (<200 ng) of DNA was used as input in a PCR with 1 μL LongAmp polymerase (New England Biolabs, M0323S), 5 µL 5x LongAmp Taq Reaction Buffer (New England Biolabs, M0323S), 3.75 μL 2 mM DNTPs (e.g., Sigma-Aldrich, D7295-.2ML), 12.25 μL Milli-Q water, and 1 μL each of primers pFL1 and pRL1 (each at 10 μM in Milli-Q water) following the manufacturer’s instructions for PCR with a total reaction volume of 25 μL and a thermocyler program of 94°C (pre-heating); 94°C for 30 seconds, 30 cycles of 94°C for 30 seconds, 58°C for 1 minute, 65°C for 3 minutes; and finally 65°C for 10 minutes. Less than 8 μL PCR product was resolved on a 1% agarose gel to verify the size of amplified bands. The remaining PCR product was purified with an NEB Monarch^®^ DNA cleanup kit (New England Biolabs, T1030L) following the manufacturer’s protocol exactly and the eluted DNA sent to Primordium Labs for long-read (Oxford Nanopore) sequencing (“Linear/PCR” submission option).

Analysis of the long-read sequencing of the entire locus in SnapGene (by aligning to the expected insertion sequence as listed in **Supplementary Table 2**) revealed that there were two copies of the SP-oScarlet construct inserted. *In silico* translation with Expasy (https://web.expasy.org/translate/; 3’5’ Frame 3, oScarlet protein in frame) showed that there was a stop codon after the first copy, compatible with the presence of only one copy of oScarlet protein (**Fig. 1 – Fig. Supplement 1a**; full sequence of actual insertion in **Supplementary Table 2**).

### Preparation of killifish brain sections

To prepare killifish brain sections, we first sacrificed fish in ice water (in most experiments, unperfused fish; in experiments in Fig. 3b, fish perfused with sulfo-NHS-biotin, see “Intracardial perfusion,” below). Whole fish heads were cut off and incubated overnight at 4°C in the dark with rocking in 1-5 mL 4% paraformaldehyde (PFA; e.g., Electron Microscopy Sciences, 15710 diluted 1:4 in phosphate-buffered saline (PBS) (e.g., Corning, 21-040-CV)) in PBS (volume used dependent on fish head size) in a 15-mL conical tube (e.g., Falcon, 352196). The following day, 3 washes with PBS (same volume as 4% PFA used in the preceding step) were performed for at least 1 hour each at 4°C in the dark with rocking. After the final wash, fish brains were dissected out of the heads in cold PBS using forceps (e.g., Fine Science Tools, 11252-20), starting with removal of the lower jaw, then the upper part of the head and eyes, and finally freeing the brain from the skull. The brains were then sunk in 0.5-1 mL 30% (weight/volume) sucrose (Sigma-Aldrich, S0389-1KG) in PBS overnight at 4°C in the dark. Once the brains had sunken, they were embedded in Optimal Cutting Temperature (OCT) compound (Fisher HealthCare, 4585). OCT blocks containing the brains were frozen over dry ice and stored at −80°C until sectioning.

Sagittal sections with a thickness of 50 µm were generated using a cryostat (Leica, CM3050 S) and MX35 Ultra blade (Epredia, 3053835) and added to Superfrost Plus slides (Fisherbrand, 1255015) that were stored at −20°C until further processing. Sections with visible and undamaged forebrain, midbrain and hindbrain tissue were included, with approximately position-matched sections (based on visual inspection of brain tissue gross morphology) used where appropriate for comparisons. RNase-free reagents and technique (namely, spraying tools and surfaces with 70% ethanol and RNaseZap (Invitrogen, AM9780, AM9782)) were used for all experiments in which hybridization chain reaction (HCR) was later performed on the brain sections.

### Hybridization chain reaction

To perform hybridization chain reaction (HCR)^73^ on killifish brain sections, we first placed slides with 50 µm-thick killifish brain sections were placed in RNase-free PBS at room temperature for at least 20 minutes to wash off excess OCT (preceded by a heating step at 37°C for ∼10 minutes in some experiments to minimize the occurrence of sections detaching from the slides).

After washing off OCT, slides were then permeabilized in 1% Triton (e.g., Sigma-Aldrich T8787-50ML) in 100% methanol (e.g., Sigma-Aldrich 34860-100ML-R) for at least 1 hour at −20°C, then washed twice in RNase-free 2x saline sodium citrate (SSC; diluted from 20x SSC (ThermoFisher, AM9763) in RNase-free water) with 0.1% Triton (250 µL per slide) for 15 minutes per wash at room temperature. HCR hybridization buffer (Molecular Instruments, “HCR Probe Hybridization Buffer,” for tissue sections; 250 µL per slide; preheated to 37°C) was applied to the slides, which were incubated at 37°C for 30 minutes with a coverslip on top. After the 30 minutes incubation, this hybridization buffer was removed and replaced with hybridization buffer containing the appropriate probe sets diluted 1:100 from stock. Once the hybridization buffer with probes sets had been applied, the slides were then incubated at 37°C overnight with a coverslip on top.

Sequences for all HCR probe sets used in this study are listed in **Supplementary Table 2**. These sequences were determined by a Python script that used the most 5’ or 3’ (as noted in **Supplementary Table 2**) 1012 bp of the longest killifish mRNA sequence for the gene of interest (or the entire mRNA sequence, in the case of shorter genes) to design up to 22 pairs of probes 22 bp in length each, with a 2 bp spacer between probes. Probes sets were ordered in tubes as lyophilized 50 nM oPools (IDT custom-synthesized) and resuspended in 50 µL of TE8.0 (Invitrogen, AM9849). After a 5 minutes wait, the tubes were vortexed and 20 µL of the resuspension solution was added to 20 µL of TE8.0. This diluted resuspension solution was vortexed, spun down, and spit into 5 µL aliquots which were used as the probe set stocks.

After the overnight incubation with probes sets diluted in hybridization buffer, HCR wash buffer (Molecular Instruments, “HCR Probe Wash Buffer,” for tissue sections; 250 µL per slide; preheated to 37°C) was applied to was applied to the slides, which were incubated at 37°C for two washes of 30 minutes each with a coverslip on top. The slides were then washed twice in RNase-free 2xSSC with 0.1% Triton (250 µL per slide) for 15 minutes per wash at room temperature. Next, HCR amplification buffer (Molecular Instruments, “HCR Amplification Buffer,” for tissue sections; 100 µL per slide; equilibrated to room temperature) was applied to the slides, which were incubated at room temperature for at least 30 minutes with a coverslip on top.

During this incubation, amplification hairpins (Molecular Instruments; 2 µL per hairpin per slide) were prepared. Hairpins were first snap-heated at 95°C for 5 min, followed by 30 minutes of cooling at 4°C. Hairpins used for each probe in this study are listed in **Supplementary Table 2** (see “Notes”). Hairpin solution (100 total µL per slide) was prepared by adding hairpins (2 µL per hairpin per slide) to amplification buffer. After applying hairpin solution, slides were incubated at room temperature overnight in the dark.

After overnight incubation with hairpin solution, slides were washed twice in RNase-free 2xSSC with 0.1% Triton (250 µL per slide) for 5 minutes per wash at room temperature in the dark, then washed once in RNase-free 2xSSC with 0.1% Triton (250 µL per slide) for 5 minutes at room temperature in the dark, and then washed twice in RNase-free 2xSSC with 0.1% Triton (250 µL per slide) for 5 minutes per wash at room temperature in the dark before mounting in ProLong Gold with 4’,6-diamidino-2-phenylindole (DAPI; Thermo Fisher Scientific, P36931) under a coverslip. Mounted slides were dried overnight at room temperature in the dark and stored at 4°C in the dark until imaging.

For experiments in which HCR and immunofluorescence staining were performed sequentially on the same slides, after overnight incubation with hairpin solution, slides were washed five times in RNase-free 2xSSC with 0.1% Triton (250 µL per slide) for 5 minutes per wash at room temperature in the dark, before proceeding to blocking for immunofluorescence staining.

### Immunofluorescence staining

To perform immunofluorescence staining on killifish brain sections, we first heated slides with 50 µm-thick killifish sagittal brain sections for 10 minutes at 37°C, then placed the slides in PBS at room temperature for at least 20 minutes to wash off excess OCT. Slides were then permeabilized in 0.5% Triton in methanol for at least 1 hour at −20°C, then washed twice in PBS for 5 minutes per wash at room temperature.

Next, for blocking, circles were drawn around sections with a hydrophobic pen (Edding 751) blocking buffer (PBS with 5% normal donkey serum (NDS; Fisher Sci., SP-072-VX10) and 1% bovine serum albumin (BSA; from 10% BSA stock; Miltenyi Biotec, 130-091-376)) was added to the areas within the circles for a 30 minutes incubation at room temperature. After blocking, the blocking buffer was removed and primary antibodies were added for overnight incubation at 4°C in the dark. All primary antibodies used in this study are listed in **Supplementary Table 3** along with the dilution (in blocking buffer) at which they were used.

After incubation with primary antibodies, slides were washed 6 times with PBS with 0.1% Triton for 10 minutes each at room temperature in the dark, and then washed 3 times with PBS for 5 minutes each at room temperature in the dark, before the PBS was dried off and secondary antibodies and DAPI (Fisher Scientific, 62248) (all diluted 1:500 in blocking buffer) were added for 2 hours incubation at room temperature in the dark. All secondary antibodies used in this study are listed in **Supplementary Table 3**.

After incubation with secondary antibodies, slides were washed 6 times with PBS with 0.1% Triton for 10 minutes each at room temperature in the dark, and then washed 3 times with PBS for 5 minutes each at room temperature in the dark before mounting in ProLong Gold (ThermoFisher, P36930) under a coverslip. Mounted slides were dried overnight at room temperature in the dark and stored at 4°C in the dark until imaging.

### Tissue clearing

To visualize oScarlet protein in SP-oScarlet brains at a whole-brain level, we performed tissue clearing using a modified Adipo-Clear protocol^170^. A dissected young adult SP-oScarlet brain in PBS (processed as described in “Preparation of killifish brain sections,” above, up to dissection, then placed in room temperature PBS) was subjected to the Adipo-Clear protocol with ethyl cinnamate (Sigma-Aldrich, 112372) was used in place of dibenzyl ether. An anti-RFP antibody used to visualize oScarlet (details in **Supplementary Table 3**).

### Imaging

Unless otherwise noted, all images were taken with a Zeiss LSM900 confocal microscope with Zen3.0 Blue Edition software. Zeiss Immersol Oil 518F (Zeiss 4Y00-R0DY-1007-3VF3) was used with the 40x and 63x magnification objectives. Imaging parameters, including the magnification objective used for each image, are listed in **Supplementary Table 4**.

The images in Fig. 1d, Fig. 1e, and **Fig. 2 – Fig. Supplement 1a** were taken with a Miltenyi Biotec Ultramicroscope Blaze, an EVOS M5000 tissue culture microscope, and a Zeiss Elyra7 structured illumination microscope (at Stanford’s Cell Sciences Imaging Facility (RRID: SCR_017787)), respectively.

### Image processing and quantification

Image analysis scripts are listed in **Supplementary Table 4**. Confocal images taken were z-stacks with 10-20 slices 0.5 μm apart. Maximum intensity projections of 10 slices per image were generated in Fiji/ImageJ using the macros listed in **Supplementary Table 4** and saved as TIF files. The max projection TIF files were loaded into QuPath (v0.6.0)^171^ for automated cell detection and, where needed to detect RNA transcripts, subcellular spot detection using Groovy scripts as listed in **Supplementary Table 4**.

Unless otherwise noted, all images shown in this manuscript were processed (maximum intensity projection, brightness/contrast adjustment, cropping) with Fiji/ImageJ. The image in Fig. 1d was processed in Imaris (specialized software for 3D image processing) and further processed in Fiji/ImageJ. The image in **Fig. 2 – Fig. Supplement 1a** was pre-processed with Zen Blue Software (for structured illumination mathematical correction) on the microscope used for image acquisition (at Stanford’s Cell Sciences Imaging Facility, RRID: SCR_017787) prior to processing in Fiji/ImageJ.

Since *APOEB* HCR signal was strong and individual transcript spots were often difficult to distinguish, mean *APOEB* RNA intensity per cell in 10x magnification images was used as a readout of *APOEB* expression. For all other HCR probes used, individual transcript spots were counted in 40-63x magnfiication images. QuPath measurements were exported as .csv files and analyzed in R (v4.5.1).

In quantifications of oScarlet immunofluorescence staining, oScarlet^HIGH^ cells were defined as those with both a mean fluorescence intensity in the oScarlet channel of >2.5 standard deviations higher than the mean of all cells in the image and maximum fluorescence intensity in the oScarlet channel of >25,000, due to the punctate nature of bright oScarlet protein signal.

In quantifications of dextran-Oregon Green 488 native fluorescence, dextran-positive cells were defined as those with both a mean fluorescence intensity in the dextran channel of >2.5 standard deviations higher than the mean of all cells in the image and maximum fluorescence intensity in the dextran channel of >25,000, due to the punctate nature of dextran signal.

In quantifications of *APOEB* HCR signal, cells with high *APOEB* RNA expression were defined as those with a mean fluorescence intensity in the *APOEB* RNA channel of >2.5 standard deviations higher than the mean of all cells in the image, since *APOEB* RNA signal was largely cell body-filling.

In quantifications of colocalization with vasculature, vasculature was defined as cells with a mean fluorescence intensity in the sulfo-NHS-biotin channel of >1.5 standard deviations higher than the mean of all cells in the image, since sulfo-NHS-biotin signal was largely cell body-filling. Cells colocalizing with vasculature were defined as those with a centroid <5 μm from the centroid of any vasculature cell (inclusive of vasculature cells themselves).

For representative sampling of all brain regions for quantification, we imaged one region of interest from each of the forebrain, midbrain, and hindbrain across 1-3 sagittal sections per animal, avoiding areas of visually obvious tissue damage.

In Fig. 3a and **Fig. 3 – Fig. Supplement 1a**, four images (one image per animal for each of four animals; listed in **Supplementary Table 4**) were excluded from quantification because they contained regions with a visually obvious very high density of cells with high *APOEB* RNA expression, which could potentially reflect a response to injury (see also “Injection of dextran into killifish brains,” below).

For quantifications in Fig. 3b, brain external borders were generally avoided when selecting regions of interest, because sulfo-NHS-biotin signal was much brighter at these external borders as compared to other areas in the brain, which we could bias quantification. As such, our analysis mostly examined *APOEB*+ cells in the brain interior.

### Injection of dextran into killifish brains

To test the endocytic properties of cells in the killifish brain, we injected dextran-Oregon Green 488 (70 kDa) (Thermo Fisher Scientific, D7173), aiming for the ventricle to maximize distribution of dextran through the cerebrospinal fluid. Only females were used for these experiments since adult males have thick muscle above the skull that makes it difficult to visually locate the brain.

The brain injection procedure bears similarities to cerebroventricular microinjection procedures used for adult zebrafish^96,172^, though specific adaptations were made for killifish. Brain injection needles (made identically to embryo microinjection needles) were filled with a mixture of 90% dextran (1 mg/mL dissolved in PBS) and 10% phenol red (for visualization), for a final dextran concentration of 900 μg/mL, using Microloader tips.

After loading, the needles were unsealed by breaking their points with fine forceps. Prior to injection, fish were individually anesthetized in 300 mg/L Tricaine (dissolved in system water) buffered with sodium bicarbonate to pH ∼7. Successful anesthesia was confirmed by the fish being non-responsive to tail pinching, at which point the fish was placed in between pins on a Tricaine-filled dish underneath the injection microscope.

Injections were performed with a Pneumatic Picopump (World Precision Instruments, PV380) with settings Vacuum = 0 in Hg (vac setting), Hold = 0 psi, Eject = 5-10 psi (Vent = hold), Range = 10 s, Period = 50, Duration = timed, with the needle stabilized and positioned using a DC-3K right apparatus (World Precision Instruments, DC3001R, 00-45-201-0000). We used a micrometer (Fisherbrand, 12-561-SM1) to estimate an approximate ejection volume of <0.5 μL (based 5 test ejections of 10% phenol red in PBS into a mineral oil (e.g., Sigma-Aldrich, M5904-5ML) droplet with similar needles, unsealing procedure, and approximately equal pressure settings as used for brain injections; although injecting through the skull further unseals the needle). We used the minimum Eject pressure necessary to eject liquid for this procedure, to minimize damage to the brain.

To aim for the ventricle, needles were inserted at approximately a 45° angle through the skin and skull at the most medial point of the forebrain/midbrain border (where the two lobes of the telencephalon and two lobes of the optic tectum meet). This point was approximately aligned along the anterior-posterior axis with the most posterior part of the eyes.

No brain coordinates were used to estimate the depth the needle needed to reach before ejection (by pressing foot pedal); the depth of needle insertion was approximately optimized by practice injections, with immediate spread of phenol red used as a readout of successful injection. We note that engulfment of dextran by brain macrophages was observed in some cases where immediate spread of phenol red upon ejection was not observed, possibly due to clogging in the needle limiting the volume of liquid ejected.

Fish were then placed in recovery tanks with fresh system water (not on aquatic system) and sacrificed 1 day (∼20-26 hours) after dextran injection.

We excluded one animal from the quantifications in Fig. 3a and **Fig. 3 – Fig. Supplement 1a** based on visually obvious lack of dextran signal throughout the brain (**Fig. 3 – Fig. Supplement 1b**). Of the 38 remaining images from six remaining animals, we additionally excluded four images (one each from four different animals) that contained regions of interest with visually apparent very high densities of *APOEB*+ cells, which could reflect a response to injury. These four excluded images are listed in **Supplementary Table 4** (row 14, “Notes”).

### Intracardial perfusion

To visualize vasculature, we perfused wildtype killifish with sulfo-NHS-biotin (Thermo Scientific, 21335). The perfusion procedure was similar to that described by Costa et al., 2026^55^. Perfusion was performed with a syringe pump (KD Scientific, Legato 200 Series, 788200 syringe pump) with 30-gauge hubless needles with a point style 4 bevel (Hamilton Syringe, custom needle, 22030-01; 30 gauge, Hubless Needle, No Hub, 30 gauge, 1.5 inches length, point style 4 [12°]) attached using ∼0.25 meters of BD Intramedic PE Tubing (BD, 427400) to a 30 gauge metal hub blunt-end Luer needle (Hamilton Syringe, custom needle, 7748-16; 30 gauge, Metal Hub Needle, Point Style: 3; Needle Length: 0.375 inches) on a 20-mL disposable syringe with Luer Lock tip (‘Sterile Syringe Only with Luer Lock Tip’, Amazon, B08FJCSLFC).

Adult killifish were anesthetized in 300 mg/L Tricaine (dissolved in system water; e.g., Pentair, TRS1) buffered with sodium bicarbonate to pH ∼7, with the underbelly cut so as to expose the heart (full details of cutting described by Costa et al., 2026^55^), with the perfusion needle inserted into the bulbus arteriosus. The perfusion flow rate was 3.5 mL/minute. First, 6 mL of 50 mM Tris pH 8.0 (filled in the disposable syringe; Thermo Fisher Scientific, 15-568-025) was perfused, followed by 4 mL of 0.5 mg/mL sulfo-NHS-biotin in PBS, and then 2 mL of 50 mM Tris pH 8.0.

After perfusion, fish were sacrificed in ice water and their carcasses incubated overnight at 4°C in the dark with rocking in >5 mL 4% paraformaldehyde in PBS (e.g., diluted from 16% paraformaldehyde in PBS (Electron Microscopy Sciences, 15714-S)) in a 15-mL conical tube (e.g., Falcon, 352196). Cuts were made in the underbelly to expose the internal organs and allow for complete fixation. The following day, 3 washes with PBS (same volume as 4% paraformaldehyde used) were performed for at least 1 hour each at 4°C in the dark with rocking. After this, the PBS was exchanged with 100% methanol (Sigma-Aldrich, 34860-4L-R) and the fish were stored long-term (>1 year) at −20°C. Before sectioning, heads were cut off and rehydrated with a reverse gradient of 2 mL each of (1) 75% methanol and 25% PBS; (2) 50% methanol and 50% PBS; (3) 25% methanol and 75% PBS; and finally (4) 100% PBS, sequentially for 15-45 minutes each on ice. Then, an additional wash in 100% PBS was performed for 15 minutes, and then brains were dissected for preparation of killifish brain sections.

### Dissociation of killifish brains for single-cell RNA sequencing

We conducted two single-cell RNA sequencing experiments: one in October 2024 (Fig. 4e) and one in August 2025 (Fig. 1d). In the October 2024 experiment, we sequenced FACS-sorted oScarlet^HIGH^ cells from old male SP-oScarlet brains (n = 3 fish, pooled) as well as FACS-sorted Live cells from old male wildtype brains (n = 3 fish, pooled). In the August 2025 experiment, we sequenced FACS-sorted oScarlet^HIGH^ and oScarlet^LOW^ cells from young male SP-oScarlet brains (n = 4 fish, pooled) as well as FACS-sorted oScarlet^HIGH^ from old male wildtype brains (n = 4 fish, pooled).

For single-cell RNA sequencing experiments, fish were always sacrificed in the morning on a weekday, having not received feeding since the previous afternoon. Males were used for these experiments due to their larger size, maximizing cell yield for single-cell RNA sequencing.

For dissociation, we used protocols similar to those described by Mariën et al., 2024^173^. Fish (not perfused) were sacrificed in ice water and brains were dissected in ice-cold PBS using forceps, starting with removal of the lower jaw, then the upper part of the head and eyes, and finally freeing the brain from the skull. After dissection, brains were cut into pieces approximately the size of a half telencephalon. Per sample (each sample consisted of a pool of 3-4 brains matched in age, genotype, and sex), all tissue pieces were transferred into a 1.5-mL microcentrifuge tube containing 800 µL of ice-cold dissection medium (DMEM/F12 with HEPES (Thermo Fisher Scientific, 11330032) with actinomycin D (transcription inhibitor; Sigma-Aldrich, A1410-2MG) added at 1:200, triptolide (transcription inhibitor; Sigma-Aldrich, T3652-1MG) added at 1:1000, anisomycin (translation inhibitor; Sigma-Aldrich, A9789-5MG) added at 1:368.5 and put on ice until all brains were dissected (up to 2 hours).

Once all the dissected tissues were ready in dissection medium on ice, fresh papain solution was prepared. Papain solution consisted of 50 µL of papain (suspension, from papaya latex; Worthington, LS003126), 20 µL DNase I solution (10 mg/mL; e.g., STEMCELL Technologies, 07900; dissolved in Milli-Q water), and 40 µL of L-cysteine solution (12 mg/mL) per 1 mL of DMEM/F12, with actinomycin D added at 1:200, triptolide at 1:1000, anisomycin at 1:368.5. This solution was sterilized through a 0.2-0.22-µm-pore-size filter (Sigma-Aldrich, WHA9914-2502 or Fisher Scientific, SLGP033RS) and vortexed to mix. L-cysteine solution was prepared by adding 120 mg of L-cysteine hydrochloride (e.g., Sigma-Aldrich, C6852-25G) to 10 mL of filtered, autoclaved Milli-Q water, and stored for up to 2 weeks at 4°C or 1 year at −20°C.

The dissection medium was removed by pipetting and 1 mL fresh papain solution was added to each sample. To start digesting the brain tissue, tubes with the samples were placed in an incubator for ∼6 minutes at 37°C. After the 6 minutes, the tubes were removed from the incubator and the brain tissue was dissociated mechanically by pipetting up and down ∼13 times with a 1-mL pipette tip (e.g., Fisher Scientific 12-111-164) cut at 0.5 cm. Four additional rounds of incubation and mechanical dissociation were performed. In the second round, mechanical dissociation was performed with a 1-mL pipette tip cut at 0.2 cm, and in subsequent rounds, mechanical dissociation was performed with an uncut 1-mL pipette tip. Papain digest was limited to 5 rounds (30 minutes) total to avoid pre-apoptotic cells. After this digest, cells were transferred into 5-mL FACS tubes with strainer caps (Corning, 352235), using sufficient pressure to make sure that the cells passed through the strainer. Then 3 mL of washing solution for single-cell suspension was passed through the strainer to clean the strainer and stop the enzymatic reaction. Washing solution for single-cell suspension was prepared by adding 975 µL of glucose (e.g., Sigma, G7021-1KG) 15% (weight/volume) in Milli-Q water, 250 µL of HEPES 1 M (e.g., Sigma-Aldrich, H3375-100G; dissolved in Milli-Q water), and 2.5 mL of FBS (fetal bovine serum, e.g., Millipore Sigma, F2442-500ML) to 46.9 mL DPBS (Dulbecco’s phosphate-buffered saline; Thermo Scientific, 14190094), stored for up to 2 months at 4°C, and sterilized with a 0.2-0.22-µm-pore-size filter. After addition of washing solution, contents of the FACS tubes were poured into 50 mL conical tubes (e.g., Thermo Fisher Scientific, 14-432-22) for subsequent centrifugation.

The conical tubes containing the cell suspensions were centrifuged at 500g for 10 minutes at 4°C, and the supernatants were carefully decanted and discarded. Debris was removed using the debris-removal solution, according to the manufacturer’s protocol. Cells were resuspended carefully in each tube with 15.5 mL DPBS. Then 4.5 mL of cold debris-removal solution (Miltenyi Biotec, 130-109-398), was added and gently mixed by inverting. This solution was gently overlaid (drop by drop) with 20 mL of cold DPBS (creating two phases), before centrifugation at 3000g for 10 minutes at 4°C, with full acceleration and full brake. After centrifugation, three phases formed; the top two phases were aspirated and discarded. If the phases were hard to distinguish, a conservative approach was taken and the top 20 mL was aspirated and discarded. Then each tube was filled up with cold DPBS (20 mL), closed, mixed by inverting three times with no vortexing and centrifuged at 1000g for 10 minutes at 4°C. The supernatant was aspirated and discarded, and the pellet was resuspended in 500 μL PBS to generate the single-cell suspension.

### FACS for single-cell RNA sequencing

Single-cell suspensions were moved to 1.2-mL FACS library tubes (e.g., VWR, 83009-678), to which 5 μL Zombie NIR™ live/dead stain (BioLegend, 423105) diluted 1:10 in PBS was added. These tubes were incubated 15 minutes at room temperature in the dark, and then centrifuged at 300g for 5 minutes at 4°C. For each tube, the supernatant was removed to 100 μL, and the pellet was resuspended in 400 μL of PBS with 0.5% BSA (diluted from 35% BSA; Sigma-Aldrich A7979-50ML).

The contents of each tube were strained through a 5-mL FACS tube with strainer cap, vortexed to minimize cell clumping, and then run on a Sony MA900 cell sorter at about 1500-2000 events per second, with FSC threshold = 6%, FSC gain = 12, BSC gain = 35%, and all fluorophores gain = 40%. Compensation was not used due to the presence of only two fluorophores (oScarlet, same spectra as mScarlet; and Zombie NIR™) whose spectra were determined to be sufficiently non-overlapping per Thermo SpectraViewer. For each sample, 100,000 events were recorded and .fcs files were exported for analysis in FlowJo (v10.0.0). The flow cytometry plot in Fig. 1e was on Live cells as identified from the gating strategy shown in **Fig. 1 – Fig. Supplement 1e**. oScarlet^HIGH^ cells were defined as those having a FL3-A::mCherry-A (uncompensated) intensity of approximately >800 (October 2024 experiment) or >1000 (August 2025 experiment). oScarlet^LOW^ cells were defined as those having a FL3-A::mCherry-A (uncompensated) intensity of approximately <500.

Each sample from which oScarlet^HIGH^ cells were collected was run in its entirety on Normal mode (maximizes yield) to collect as many oScarlet^HIGH^ cells as possible (with yields ranging from approximately 35,000-65,000 oScarlet^HIGH^ cells according to Sony MA900 event count). For the Live cells collected from wildtype brains in the October 2024 experiment and the oScarlet^LOW^ cells collected from young adult SP-oScarlet fish in the August 2025 experiment, these were also run in Normal mode, but approximately 240,000 and 167,000 cells were collected respectively (according to Sony MA900 event count), which did not represent the entire sample, to avoid overloading the 10x chip in single-cell RNA sequencing. Cells were collected in 500 µL of RNase-free PBS with 1% BSA and 0.1% glucose in 1.5 mL DNA LoBind tubes (Eppendorf, 0030108418).

### Single-cell RNA sequencing

After cell collection, tubes containing the cells were centrifuged at 300g for 5 minutes at 4°C. Supernatants were removed to 43.2 μL, and Master Mix was prepared and added to each sample exactly following the instructions in the 10x Genomics Chromium Next GEM Single Cell 3’ Reagent Kits v3.1 User Guide. Cells were not counted, in order to load all cells collected in each tube. Once Master Mix was added, the contents of each tube were pipetted up and down (to resuspend the cells fully) and loaded in their entirety onto the 10x chip.

After this, the instructions in the 10x Genomics Chromium Next GEM Single Cell 3’ Reagent Kits v3.1 User Guide were followed exactly with 11-12 cycles used in the cDNA amplification PCR and 13-14 cycles used in the Sample Index PCR (October 2024 experiment, 12 and 13 cycles, respectively; August 2025 experiment; 11 and 14 cycles, respectively).

Libraries were sequenced by Novogene on a NovaSeqX Plus sequencer (150 bp paired-end reads) to an approximate depth of 215-333 million reads per sample.

### Identifying homologous genes across species

Killifish gene names were converted to their equivalents in human, mouse, or other species using the table presented in **Supplementary Table 5**, which was generated as described in the next paragraph. Killifish can have multiple paralogs corresponding to a single mammalian gene, due to whole-genome duplication in teleost fish^174,175^. In such cases, for visualization purposes, one killifish paralog was selected (specified in figure legend). In cases where one killifish paralog had a much higher expression level than the other paralogs, the highest-expressed killifish paralog was selected.

Over half of the killifish NCBI gene names (11,715 out of 22,194) consist of locus numbers. Therefore, we generated new gene symbols for all the killifish protein coding genes, based on NCBI gene descriptions for killifish and human, and orthologs between killifish and other animals. For ortholog identifications, we ran sequence comparison using BLAST (ncbi-blast-2.5.0+) between killifish and 36 other animals, including multiple tetrapod, and fish species. We then generated a prioritized scheme for gene symbol assignment based on the following rules. We kept the gene symbols assigned by NCBI annotation. If there were no gene symbol assigned by NCBI, we matched the NCBI gene description between killifish and human and assigned the human gene symbols where the gene descriptions matched between the two species. For rest of the genes, we generated a consensus gene symbol based on ‘majority rules’ using ortholog symbols from 36 other animals. In case of ties, priority was given to the tetrapod symbols. We were able to assign proper gene identities to 93% of the locus numbers and improve gene symbols for 96.4% of all protein coding genes.

In the text, figures, and legends of this manuscript, all killifish gene names are capitalized and presented in italics (e.g., *ELAVL3*).

### Analysis of single-cell RNA sequencing data

Demultiplexed reads provided by Novogene were aligned to the African turquoise killifish genome (Nfu_20140520) using Cell Ranger. A custom genome including the SP-oScarlet knockin insertion (as determined from sequencing as in **Fig. 1 – Fig. Supplement 1a**) as a separate chromosome was built according to the instructions provided by 10x Genomics (https://www.10xgenomics.com/support/software/cell-ranger/latest/tutorials/cr-tutorial-mr, “Add a custom gene to the FASTA and GTF”) for alignment of reads from SP-oScarlet fish, although we note that the *SP-oScarlet* transcript was difficult to detect even in *ELAVL3*+ cells. This is potentially explained by the *SP-oScarlet* transcript’s position several kilobases upstream of the 3’ end of *ELAVL3*, as *ELAVL3* has a long untranslated region. The *oScarlet* transcript could be detected by HCR in *ELAVL3*+ cells *in situ* (**Fig. 1 – Fig. Supplement 1c**).

After alignment, raw_feature_bc_matrix outputs were downloaded and a Seurat^176^ (v5.3.0) object was created (separately for each sample) using the Read10x and CreateSeuratObject functions (min.cells = 3, min.features = 200). For quality control, each of these Seurat objects was individually filtered to include cells with nCount_RNA > 1000, nFeature_RNA > 500, and <10% mitochondrial reads. No doublet removal was performed. After testing various doublet removal methods in preliminary analyses and ensuring that our key conclusions were robust to doublet removal method, we reasoned that due to the lack of reliable killifish cell type markers (particularly for myeloid cells), it would be challenging to assess the accuracy of singlet vs. doublet classifications. Quality control metrics per individual Seurat object are listed in **Supplementary Table 6**. Seurat objects were merged as appropriate. After each merge, JoinLayers was run on the merged object to allow Seurat to identify cluster markers and differentially expressed genes.

For all objects, merged or unmerged, clusters were defined by running NormalizeData (normalization.method = “LogNormalize”, scale.factor = 10000), FindVariableFeatures (selection.method = “vst”, nfeatures = 2000), ScaleData, RunPCA, RunUMAP (dims = 1:30), FindNeighbors (dims = 1:30) and finally FindClusters (resolution = 0.1-0.2). Cell types were defined by running FindMarkers for low-resolution clusters and manually matching markers to cell types based on literature, most notably Ayana et al., 2024^68^ (killifish-specific) and Tabula Muris^78^. Full lists of low-resolution cluster marker genes among all young adult cells from the August 2025 experiment can be found in **Supplementary Table 7**.

For subsetting to macrophages, *APOEB*+ *IBA1*+ *LCP1*+ clusters were selected based on expression from VlnPlot. These markers were used by Ayana et al. to define “Microglia” in their wildtype killifish forebrain single-cell RNA sequencing datasets^68^, although our analysis shows that *APOEB*+ *IBA1*+ *LCP1*+ killifish brain cells have an expression profile more similar to that of mouse monocyte-derived macrophages rather than mouse microglia.

Young adult wildtype killifish forebrain macrophages were identified by downloading the dataset SRR24058857 (from Ayana et al., 2024^68^) from NCBI Gene Expression Omnibus/Sequence Read Archive, aligning it to the Nfu_20140520 genome using Cell Ranger, and creating a Seurat object and performing quality control exactly as described above for this study’s original datasets, and then subsetting to the (one) *APOEB*+ *IBA1*+ *LCP1*+ cluster.

To compare killifish and mouse brain macrophages, we used a Seurat object corresponding to all young adult mouse brain immune cells analyzed by Barr et al., 2025 (see Barr et al., 2025^75^ for details of single-cell RNA sequencing, genome alignment, quality control, annotation of cell types, etc.). This object, which includes cells collected at multiple circadian timepoints, was first subset to the circadian timepoint ZT12 (Time = ZT12), which corresponds to the beginning of the active phase (roughly equivalent to the circadian timepoint at which our killifish single-cell data was collected). The object was then subset to microglia, BAMs, and MDMs (Broad_Cell_Annotation %in% c(“Microglia”, “PBM”, “MDM”).

Then, we determined markers of mouse microglia, BAMs, and MDMs using FindAllMarkers (test.use = “wilcox”, min.pct = 0.1, logfc.threshold = 0.5). All markers of microglia, BAMs, or MDMs, including genes that were markers of multiple cell types (or all killifish homologs of these genes) were used for principal component analysis (PCA). This PCA was performed starting with a Seurat object for each of the following cell populations:

1. Young adult mouse brain macrophages (subset from all young adult mouse brain immune cells; Barr et al., 2025^75^; Broad_Cell_Annotation %in% c(“Microglia”, “PBM”, “MDM”, “IFN-M”))
2. Young adult oScarlet^HIGH^ cells (this study)
3. Young adult wildtype killifish forebrain macrophages (Seurat object generated as described above; Ayana et al., 2024^68^)

and subsetting to only the marker genes as described above, then re-running NormalizeData (normalization.method = “LogNormalize”, scale.factor = 10000), FindVariableFeatures (selection.method = “vst”, nfeatures = 2000), ScaleData, and finally RunPCA.

The strongest markers of mouse microglia, BAMs, and MDMs as depicted in the heatmaps in **Fig. 2 – Fig. Supplement 3a** and **Fig. 2 – Fig. Supplement 4a** were defined using using FindAllMarkers (test.use = “wilcox”, min.pct = 0.5, logfc.threshold = 3).

DimPlot was used to highlight cells by orig.ident (i.e., sample of origin) or cell type (reduction = “umap”) and produce PCA plots (reduction = “pca”). FeaturePlot and VlnPlot were used to highlight expression of specific genes. VlnPlot was also used to highlight sum log-normalized expression of genes in specific Gene Ontology terms. DoHeatmap was used to highlight expression of top killifish cell type marker genes as ranked by area under curve according to Presto^177^ (no thresholds applied; cell types defined manually as described above) or to highlight expression of killifish homologs of the strongest mouse microglia, BAM, and MDM markers. Full lists of cell type marker genes among all young adult cells from the August 2025 experiment as determined by Presto can be found in **Supplementary Table 8**.

Differentially expressed genes between samples were determined using FindMarkers (test.use = “wilcox”, min.pct = 0.1, logfc.threshold = 0.5), and Gene Ontology terms upregulated among these genes were determined using enrichR (“GO_Biological_Process_2025”) with a background of all human genes with killifish homologs represented among the rownames of the combined Seurat object (containing both samples) and plotted using ggplot2. Differentially expressed genes were always calculated at the level of killifish genes, then converted to human gene names for Gene Ontology analysis (enrichR disregards duplicates). The sizes of the circles in Fig. 4d and **Fig. 4 – Fig. Supplement 1d** correspond to the numbers of unique human genes with any killifish homologs among the differentially expressed genes for the comparison made. As brains from n = 3-4 individual male fish were pooled to achieve sufficient numbers of oScarlet^HIGH^ cells, statistics were calculated at the per-cell level.

Full lists of differentially expressed genes per comparison made can be found in **Supplementary Tables 9 and 12**. Full lists of Gene Ontology terms upregulated per condition can be found in **Supplementary Tables 10-11 and 13-14**.

### Dissociation of killifish brains for *ex vivo* engulfment experiments

To assess acute endocytic capacity of killifish brain macrophages, we performed *ex vivo* engulfment of ovalbumin-Alexa Fluor^®^ 647 (a 45 kDa protein) (Fisher Scientific, O34784) as well as dextran-Oregon Green 488 (a 70 kDa polysaccharide) (Thermo Fisher Scientific, D7173). For *ex vivo* experiments, fish were always sacrificed in the morning on a weekday, having not received feeding since the previous day.

Dissociation protocols largely followed those described by Mariën et al., 2024^173^. Fish were sacrificed in ice water and brains were dissected in ice-cold PBS using forceps, starting with removal of the lower jaw, then the upper part of the head and eyes, and finally freeing the brain from the skull. After dissection, brains were cut into pieces approximately the size of a half telencephalon. Per sample (each sample consisted of 1 brain), all tissue pieces were transferred into a 1.5-mL microcentrifuge tube containing 500 µL of ice-cold dissection medium (DMEM/F12 + HEPES) and put on ice.

Once all the dissected tissues were ready in dissection medium on ice, fresh papain solution was prepared. Papain solution consisted of 50 µL of papain (suspension, from papaya latex), 20 µL DNase I solution (10 mg/mL; dissolved in Milli-Q water), and 40 µL of L-cysteine solution (12 mg/mL) per 1 mL of DMEM/F12, with the appropriate substrates added at 25 µg/mL each; see details of individual experiments below, “*Ex vivo* engulfment of ovalbumin” and “*Ex vivo* engulfment of dextran”). This solution was sterilized through a 0.2-0.22-µm-pore-size filter and vortexed to mix. L-cysteine solution was prepared by adding 120 mg of L-cysteine hydrochloride to 10 mL of filtered, autoclaved Milli-Q water, and stored for up to 2 weeks at 4°C or 1 year at −20°C.

The dissection medium was removed by pipetting and 0.5 mL fresh papain solution was added to each sample. To start digesting the brain tissue, tubes with the samples were placed in an incubator for 6 minutes at 37°C. Tubes were covered with foil for all incubations, due to the use of light-sensitive fluorescent substrates. After the 6 minutes, the tubes were removed from the incubator and the brain tissue was dissociated mechanically by pipetting up and down 13 times with a 1-mL pipette tip cut at 0.5 cm. Four additional rounds of incubation and mechanical dissociation were performed. In the second round, mechanical dissociation was performed with a 1-mL pipette tip cut at 0.2 cm, and in subsequent rounds, mechanical dissociation was performed with an uncut 1-mL pipette tip. Papain digest was limited to 5 rounds (30 min) total to avoid pre-apoptotic cells. After this digest, cells were transferred into FACS tubes with strainer caps, using sufficient pressure to make sure that the cells passed through the strainer. Then 2 mL of washing solution for single-cell suspension was passed through the strainer to clean the strainer and stop the enzymatic reaction. Washing solution for single-cell suspension was prepared by adding 975 µL of glucose 15% in water, 250 µL of HEPES 1 M (dissolved in Milli-Q water), and 2.5 mL of FBS to 46.9 mL DPBS, stored for up to 2 months at 4°C, and sterilized with a 0.2-0.22-µm-pore-size filter. After addition of washing solution, contents of the FACS tubes were transferred to 15 mL conical tubes for subsequent centrifugation. It is easier to see the pellet in conical tubes than in FACS tubes; while a wide variety of tubes are likely suitable for these centrifugation steps, we note that choice of tube can affect yield.

The conical tubes containing the cell suspensions were centrifuged at 500g for 10 minutes at 4°C, and the supernatants were carefully decanted and discarded. Debris was removed using the debris-removal solution, according to the manufacturer’s protocol. Cells were resuspended carefully in each tube with 1550 µL DPBS. Then 450 µL of cold debris-removal solution was added and gently mixed by inverting. This solution was gently overlaid (drop by drop) with 2 mL of cold DPBS (creating two phases), before centrifugation at 3000g for 10 minutes at 4°C, with full acceleration and full brake. After centrifugation, three phases formed; the top two phases were aspirated and discarded. If the phases were hard to distinguish, a conservative approach was taken and the top 2 mL was aspirated and discarded. Then each tube was filled up with cold DPBS (2 mL), closed, mixed by inverting three times with no vortexing and centrifuged at 1000g for 10 minutes at 4°C. The supernatant was aspirated and discarded, and the pellet was resuspended in 500 μL PBS with Zombie Aqua™ (BioLegend, 423101) diluted 1:1000 to generate the single-cell suspension.

### Flow cytometry and data analysis for *ex vivo* engulfment experiments

To evaluate *ex vivo* engulfment by flow cytometry, 500 μL of single-cell suspension per dissociated brain were moved from 15 mL conical tubes to 1.2-mL FACS library tubes for incubation with Zombie Aqua™ live/dead dye. The suspensions were incubated 15 minutes at room temperature in the dark, and then centrifuged at 300g for 5 minutes at 4°C. For each tube, the supernatant was removed down to 100 μL, and the pellet was resuspended in 400 μL of added PBS with 0.5% BSA.

The contents of each tube were then strained through a 5-mL FACS tube with strainer cap, vortexed to minimize cell clumping, and then run on a Sony MA900 cell sorter at about 1500-2000 events per second (for some more dilute samples, 1000 events per second), with FSC threshold = 6%, FSC gain = 12, BSC gain = 35%, and all fluorophores gain = 40%. Single-color compensation controls were used (see details for individual experiments below). For each sample, 250,000 events were recorded and .fcs files were exported for analysis in FlowJo (v10.0.0). The gating strategy used to identify Live cells was similar to that shown in **Fig. 1 – Fig. Supplement 1e**, except with Zombie Aqua™ as the live/dead dye rather than Zombie NIR™. Scale data from Live cells were exported as .csv files that were analyzed in R (v4.5.1) for quantification.

### Development of an *ex vivo* engulfment assay for ovalbumin

Data for ovalbumin engulfment were acquired over three experiments. In all experiments, ovalbumin-AlexaFluor 647 [ovalbumin] was used at 25 µg/mL (diluted from a 1 mg/mL ovalbumin stock dissolved in PBS) and the fluorophores used on the Sony MA900 sorter were FL3::mCherry (for oScarlet), FL7::BrilliantViolet510 (for Zombie Aqua™ live/dead dye), and FL10::Alexa Fluor^®^ 647 (for ovalbumin).

Importantly, in optimizing the *ex vivo* ovalbumin engulfment assay, we sorted ovalbumin+ cells and verified by confocal microscopy that ovalbumin was localized inside cells rather than binding to the surface of cells.

In the first experiment, four male 41 days old (young adult) and four male 160 days old (old) heterozygous SP-oScarlet killifish were used, as well as single-color compensation controls. The compensation matrix used is included in **Supplement 2**. Ovalbumin was added to a portion the papain solution after syringe filtration. In this experiment, ovalbumin+ cells were defined as having ovalbumin (FL10-A::Alexa Fluor^®^ 647-A, compensated) intensity of >1000.

In the second experiment, four female 57 days old (young adult) and four female 161 days old (old) heterozygous SP-oScarlet killifish were used, as well as single-color compensation controls (independent compensation controls were used for this experiment). The compensation matrix used is included in **Supplement 3**. Ovalbumin was added to a portion the papain solution after syringe filtration. In this experiment, ovalbumin+ cells were defined as having ovalbumin (FL10-A::Alexa Fluor^®^ 647-A, compensated) intensity of >1000.

In the third experiment, two male 63 days old (young adult), and two male 148 days old (old), two female 63 days old (young adult), and two female 167 days old (old) heterozygous

SP-oScarlet killifish were used. No new compensation samples were used in this experiment, as the sex-respective compensation matrices from the previous experiments were re-used. Unlike in the previous experiments, ovalbumin was added to the papain solution before syringe filtration. In this experiment, ovalbumin intensity values were generally lower than observed in the previous two experiments and ovalbumin+ cells were defined as having ovalbumin (FL10-A::Alexa Fluor^®^ 647-A, compensated) intensity of >500.

Our conclusions were not affected by whether ovalbumin was added to the papain solution before or after filtration. However, there was likely some ovalbumin loss when ovalbumin was added before filtration in the third experiment, and the effective ovalbumin concentration in the *ex vivo* engulfment assay was likely lower than the 25 µg/mL reported.

Data on brightness of oScarlet^HIGH^ cells in young adult and old fish were from these ovalbumin engulfment experiments. oScarlet^HIGH^ and oScarlet^LOW^ cells were defined as having oScarlet (FL3-A::mCherry-A, compensated) intensities of >1000 (>500 in **Fig. 5 – Fig. Supplement 2b**) and <500, respectively. Values presented in Fig. 5c**-d**, **Fig. 5 – Fig. Supplement 1c-d**, and **Fig. 5 – Fig. Supplement 2b-c** were normalized to the average value among for all control samples (always presented as the leftmost category on the x-axis) of the same respective sex and experiment day.

### *Ex vivo* engulfment assay for dextran

Data for dextran engulfment were acquired in one experiment. Dextran-Oregon Green 488 (70 kDa) [dextran] was used at 25 µg/mL (diluted from a 1 mg/mL dextran stock dissolved in PBS) and the fluorophores used on the Sony MA900 sorter were FL1::Alexa Fluor^®^ 488 (for dextran), FL3::mCherry (for oScarlet), and FL7::BrilliantViolet510 (for Zombie Aqua™ live/dead dye).

Four male 56 days old (young adult) and four male 156 days old (old) heterozygous SP-oScarlet killifish were used, as well as single-color compensation controls run the previous day. The compensation matrix used is included in **Supplement 4**. On the day the eight experimental fish were run, dextran was added to the papain solution before syringe filtration.

In this experiment, dextran+ cells were defined as having dextran (FL1-A::Alexa Fluor^®^ 488-A, compensated) intensity of >800. oScarlet^HIGH^ and oScarlet^LOW^ cells were defined as having oScarlet (FL3-A::mCherry-A, compensated) intensities of >1000 (>500 in **Fig. 5 – Fig. Supplement 2d**) and <500, respectively. Values presented in Fig. 5e**-f**, **Fig. 5 – Fig. Supplement 1e-f**, and **Fig. 5 – Fig. Supplement 2d-e** were normalized to the average value among for all control samples (always presented as the leftmost category on the x-axis) of the same respective sex and experiment day.

### Statistics

No statistical methods were used to determine sample sizes. Sample sizes were determined based on previous experiments with similar types of assays in mice and killifish. No randomization or blinding methods were used, but for dissociation-based experiments, each processing step was carried out in an alternating-by-condition order (e.g., young adult, then old, then young adult, etc.) to minimize the effects of order on the results. All image quantification was automated with scripts in QuPath (v6.0.0). All statistical tests were performed in R (v4.5.1) and plots made with ggplot2 or Seurat. Unless otherwise noted in the figure legends, Wilcoxon rank-sum tests were used to determine statistical significance (paired for matched comparisons; unpaired for all other comparisons). Numbers of experimental replicates, numbers of animals per condition, and whether statistical tests were performed at the per-animal or per-cell level are indicated in individual figure legends.

## References

1. Kaur, J. et al. Waste Clearance in the Brain. Front Neuroanat 15, 665803 (2021).

2. Smyth, L.C.D., Beschorner, N., Nedergaard, M. & Kipnis, J. Cellular Contributions to Glymphatic and Lymphatic Waste Clearance in the Brain. Cold Spring Harb Perspect Biol 17(2025).

3. Kipnis, J. et al. Resolving the mysteries of brain clearance and immune surveillance. Neuron 113, 3908–3923 (2025).

4. Kida, S., Steart, P.V., Zhang, E.T. & Weller, R.O. Perivascular cells act as scavengers in the cerebral perivascular spaces and remain distinct from pericytes, microglia and macrophages. Acta Neuropathol 85, 646–52 (1993).

5. Gaudi, A.U. et al. Convergent evolution of scavenger cell development at brain borders. Nature (2026).

6. Venero Galanternik, M., et al. A novel perivascular cell population in the zebrafish brain. Elife 6(2017).

7. Uphoff, K., Suárez, I., van Impel, A. & Schulte-Merker, S. dab2 is required for the scavenging function of lymphatic endothelial cells in the zebrafish meninges. Sci Rep 14, 27942 (2024).

8. Mato, M. et al. Involvement of specific macrophage-lineage cells surrounding arterioles in barrier and scavenger function in brain cortex. Proc Natl Acad Sci U S A 93, 3269–74 (1996).

9. Kim, W.K. et al. CD163 identifies perivascular macrophages in normal and viral encephalitic brains and potential precursors to perivascular macrophages in blood. Am J Pathol 168, 822–34 (2006).

10. Drieu, A. et al. Parenchymal border macrophages regulate the flow dynamics of the cerebrospinal fluid. Nature 611, 585–593 (2022).

11. Da Mesquita, S. & Rua, R. Brain border-associated macrophages: common denominators in infection, aging, and Alzheimer’s disease? Trends Immunol 45, 346–357 (2024).

12. Vara-Pérez, M. & Movahedi, K. Border-associated macrophages as gatekeepers of brain homeostasis and immunity. Immunity 58, 1085–1100 (2025).

13. Beiter, R.M., Sheehan, P.W. & Schafer, D.P. Microglia phagocytic mechanisms: Development informing disease. Curr Opin Neurobiol 86, 102877 (2024).

14. Dong, Y. et al. Oxidized phosphatidylcholines found in multiple sclerosis lesions mediate neurodegeneration and are neutralized by microglia. Nat Neurosci 24, 489–503 (2021).

15. Lawrence, A.R. et al. Microglia maintain structural integrity during fetal brain morphogenesis. Cell 187, 962–980.e19 (2024).

16. Guldner, I.H. et al. Ageing promotes microglial accumulation of slow-degrading synaptic proteins. Nature 650, 930–941 (2026).

17. Sokolowski, J.D. & Mandell, J.W. Phagocytic clearance in neurodegeneration. Am J Pathol 178, 1416–28 (2011).

18. Da Mesquita, S. et al. Functional aspects of meningeal lymphatics in ageing and Alzheimer’s disease. Nature 560, 185–191 (2018).

19. Jeong, Y.M. et al. Differential Clearance of Aβ Species from the Brain by Brain Lymphatic Endothelial Cells in Zebrafish. Int J Mol Sci 22(2021).

20. Da Mesquita, S. et al. Aging-associated deficit in CCR7 is linked to worsened glymphatic function, cognition, neuroinflammation, and β-amyloid pathology. Sci Adv 7(2021).

21. Caron, N.S. et al. Cerebrospinal fluid biomarkers for assessing Huntington disease onset and severity. Brain Commun 4, fcac309 (2022).

22. Kaiser, S. et al. A proteogenomic view of Parkinson’s disease causality and heterogeneity. NPJ Parkinsons Dis 9, 24 (2023).

23. Johnson, E.C.B. et al. Cerebrospinal fluid proteomics define the natural history of autosomal dominant Alzheimer’s disease. Nat Med 29, 1979–1988 (2023).

24. Dammer, E.B. et al. Proteomic analysis of Alzheimer’s disease cerebrospinal fluid reveals alterations associated with APOE ε4 and atomoxetine treatment. Sci Transl Med 16, eadn3504 (2024).

25. Shen, Y. et al. CSF proteomics identifies early changes in autosomal dominant Alzheimer’s disease. Cell 187, 6309–6326.e15 (2024).

26. Peña-Bautista, C. et al. Defining Alzheimer’s Disease through Proteomic CSF Profiling. J Proteome Res 23, 5096–5106 (2024).

27. Ali, M. et al. Multi-cohort cerebrospinal fluid proteomics identifies robust molecular signatures across the Alzheimer disease continuum. Neuron 113, 1363–1379.e9 (2025).

28. Oh, H.S. et al. A cerebrospinal fluid synaptic protein biomarker for prediction of cognitive resilience versus decline in Alzheimer’s disease. Nat Med 31, 1592–1603 (2025).

29. Farinas, A. et al. Disruption of the cerebrospinal fluid-plasma protein balance in cognitive impairment and aging. Nat Med 31, 2578–2589 (2025).

30. Seo, D. et al. Cerebrospinal fluid proteomic signatures in cognitively normal individuals identify distinct clusters linked to neurodegeneration. Nat Aging 5, 2125–2141 (2025).

31. Tarasoff-Conway, J.M. et al. Clearance systems in the brain-implications for Alzheimer disease. Nat Rev Neurol 11, 457–70 (2015).

32. van Dyck, C.H. et al. Lecanemab in Early Alzheimer’s Disease. N Engl J Med 388, 9–21 (2023).

33. Sims, J.R. et al. Donanemab in Early Symptomatic Alzheimer Disease: The TRAILBLAZER-ALZ 2 Randomized Clinical Trial. Jama 330, 512–527 (2023).

34. Li, X. et al. Promising outcomes 5 weeks after a surgical cervical shunting procedure to unclog cerebral lymphatic systems in a patient with Alzheimer’s disease. Gen Psychiatr 37, e101641 (2024).

35. Bonnar, O., Eyre, B. & van Veluw, S.J. Perivascular brain clearance as a therapeutic target in cerebral amyloid angiopathy and Alzheimer’s disease. Neurotherapeutics 22, e00535 (2025).

36. Keil, S.A., Jansson, D., Braun, M. & Iliff, J.J. Glymphatic dysfunction in Alzheimer’s disease: A critical appraisal. Science 389, eadv8269 (2025).

37. Erhardt, E.B. et al. The influence of intermittent hypercapnia on cerebrospinal fluid flow and clearance in Parkinson’s disease and healthy older adults. NPJ Parkinsons Dis 11, 334 (2025).

38. Hu, C.K. & Brunet, A. The African turquoise killifish: A research organism to study vertebrate aging and diapause. Aging Cell 17, e12757 (2018).

39. Terzibasi, E. et al. Large differences in aging phenotype between strains of the short-lived annual fish Nothobranchius furzeri. PLoS One 3, e3866 (2008).

40. Terzibasi, E. et al. Effects of dietary restriction on mortality and age-related phenotypes in the short-lived fish Nothobranchius furzeri. Aging Cell 8, 88–99 (2009).

41. Tozzini, E.T., Baumgart, M., Battistoni, G. & Cellerino, A. Adult neurogenesis in the short-lived teleost Nothobranchius furzeri: localization of neurogenic niches, molecular characterization and effects of aging. Aging Cell 11, 241–51 (2012).

42. Harel, I. et al. A platform for rapid exploration of aging and diseases in a naturally short-lived vertebrate. Cell 160, 1013–1026 (2015).

43. Smith, P. et al. Regulation of life span by the gut microbiota in the short-lived African turquoise killifish. Elife 6(2017).

44. Kelmer Sacramento, E., et al. Reduced proteasome activity in the aging brain results in ribosome stoichiometry loss and aggregation. Mol Syst Biol 16, e9596 (2020).

45. Vanhunsel, S. et al. The killifish visual system as an in vivo model to study brain aging and rejuvenation. NPJ Aging Mech Dis 7, 22 (2021).

46. Van Houcke, J. et al. Aging impairs the essential contributions of non-glial progenitors to neurorepair in the dorsal telencephalon of the Killifish Nothobranchius furzeri. Aging Cell 20, e13464 (2021).

47. Vanhunsel, S. et al. The age factor in optic nerve regeneration: Intrinsic and extrinsic barriers hinder successful recovery in the short-living killifish. Aging Cell 21, e13537 (2022).

48. Louka, A. et al. New lessons on TDP-43 from old N. furzeri killifish. Aging Cell 21, e13517 (2022).

49. Teefy, B.B. et al. Dynamic regulation of gonadal transposon control across the lifespan of the naturally short-lived African turquoise killifish. Genome Res 33, 141–153 (2023).

50. Van Houcke, J. et al. A short dasatinib and quercetin treatment is sufficient to reinstate potent adult neuroregenesis in the aged killifish. NPJ Regen Med 8, 31 (2023).

51. Astre, G. et al. Genetic perturbation of AMP biosynthesis extends lifespan and restores metabolic health in a naturally short-lived vertebrate. Dev Cell 58, 1350–1364.e10 (2023).

52. Teefy, B.B. et al. Widespread sex dimorphism across single-cell transcriptomes of adult African turquoise killifish tissues. Cell Rep 42, 113237 (2023).

53. Moses, E. et al. The killifish germline regulates longevity and somatic repair in a sex-specific manner. Nat Aging 4, 791–813 (2024).

54. Di Fraia, D. et al. Altered translation elongation contributes to key hallmarks of aging in the killifish brain. Science 389, eadk3079 (2025).

55. Costa, E.K., et al. Multi-tissue transcriptomic aging atlas reveals predictive aging biomarkers in the killifish. Nat Aging (2026).

56. Morabito, G. et al. Spontaneous aging-associated inflammation and genome instability in the immune system of turquoise killifish. Nature Aging 6, 665–681 (2026).

57. Rozenberg, I., et al. A genetic toolbox for the turquoise killifish identifies sporadic age-related cancer. *bioRxiv*, 2023.05.01.538839 (2024).

58. Burgos-Ruiz, A.M. et al. Neotenic transcriptomic features in the adult turquoise killifish brain. bioRxiv, 2025.03.24.645095 (2025).

59. Williams, R.G. et al. A multi-omic atlas in the African turquoise killifish reveals increased glucocorticoid signaling as a hallmark of brain aging. *bioRxiv*, 2026.04.09.717549 (2026).

60. Matsui, H., Kenmochi, N. & Namikawa, K. Age- and α-Synuclein-Dependent Degeneration of Dopamine and Noradrenaline Neurons in the Annual Killifish Nothobranchius furzeri. Cell Rep 26, 1727–1733.e6 (2019).

61. Bagnoli, S., Fronte, B., Bibbiani, C., Terzibasi Tozzini, E. & Cellerino, A. Quantification of noradrenergic-, dopaminergic-, and tectal-neurons during aging in the short-lived killifish Nothobranchius furzeri. Aging Cell 21, e13689 (2022).

62. de Bakker, D.E.M. & Valenzano, D.R. Turquoise killifish: A natural model of age-dependent brain degeneration. Ageing Res Rev 90, 102019 (2023).

63. Valenzano, D.R. et al. Resveratrol prolongs lifespan and retards the onset of age-related markers in a short-lived vertebrate. Curr Biol 16, 296–300 (2006).

64. McKay, A. et al. An automated feeding system for the African killifish reveals the impact of diet on lifespan and allows scalable assessment of associative learning. Elife 11(2022).

65. de Bakker, D.E.M. et al. Amyloid beta precursor protein contributes to brain aging and learning decline in short-lived turquoise killifish (<em>Nothobranchius furzeri</em>). bioRxiv, 2024.10.11.617841 (2025).

66. Bedbrook, C.N. et al. Lifelong behavioral screen reveals an architecture of vertebrate aging. Science 391, eaea9795 (2026).

67. Bedbrook, C.N., Nath, R.D., Nagvekar, R., Deisseroth, K. & Brunet, A. Rapid and precise genome engineering in a naturally short-lived vertebrate. Elife 12(2023).

68. Ayana, R. et al. Single-cell sequencing unveils the impact of aging on the progenitor cell diversity in the telencephalon of the female killifish N. furzeri. Aging Cell, e14251 (2024).

69. Bindels, D.S. et al. mScarlet: a bright monomeric red fluorescent protein for cellular imaging. Nat Methods 14, 53–56 (2017).

70. Fenno, L.E. et al. Comprehensive Dual- and Triple-Feature Intersectional Single-Vector Delivery of Diverse Functional Payloads to Cells of Behaving Mammals. Neuron 107, 836–853.e11 (2020).

71. Almagro Armenteros, J.J., et al. SignalP 5.0 improves signal peptide predictions using deep neural networks. Nat Biotechnol 37, 420–423 (2019).

72. Szymczak, A.L. et al. Correction of multi-gene deficiency in vivo using a single ’self-cleaving’ 2A peptide-based retroviral vector. in Nat Biotechnol, Vol. 22 589–94 (United States, 2004).

73. Choi, H.M.T. et al. Third-generation in situ hybridization chain reaction: multiplexed, quantitative, sensitive, versatile, robust. Development 145(2018).

74. Marsh, S.E. et al. Dissection of artifactual and confounding glial signatures by single-cell sequencing of mouse and human brain. Nat Neurosci 25, 306–316 (2022).

75. Barr, H.J., et al. The circadian clock regulates scavenging of fluid-borne substrates by brain border-associated macrophages. bioRxiv (2025).

76. Barr, H.J. et al. Conserved phenotype and function of human brain border-associated macrophages in iPSC-derived models. bioRxiv, 2025.12.11.693582 (2025).

77. Becht, E. et al. Dimensionality reduction for visualizing single-cell data using UMAP. Nature Biotechnology 37, 38–44 (2019).

78. Single-cell transcriptomics of 20 mouse organs creates a Tabula Muris. Nature 562, 367–372 (2018).

79. Hammond, T.R. et al. Single-Cell RNA Sequencing of Microglia throughout the Mouse Lifespan and in the Injured Brain Reveals Complex Cell-State Changes. Immunity 50, 253–271.e6 (2019).

80. Rovira, M. et al. A single-cell transcriptomic atlas reveals resident dendritic-like cells in the zebrafish brain parenchyma. Elife 13(2025).

81. Sherr, C.J. et al. The c-fms proto-oncogene product is related to the receptor for the mononuclear phagocyte growth factor, CSF-1. Cell 41, 665–76 (1985).

82. Stanley, E.R. & Chitu, V. CSF-1 receptor signaling in myeloid cells. Cold Spring Harb Perspect Biol 6(2014).

83. Mildenberger, W., Stifter, S.A. & Greter, M. Diversity and function of brain-associated macrophages. Curr Opin Immunol 76, 102181 (2022).

84. Thion, M.S. & Garel, S. Microglial ontogeny, diversity and neurodevelopmental functions. Curr Opin Genet Dev 65, 186–194 (2020).

85. Utz, S.G. et al. Early Fate Defines Microglia and Non-parenchymal Brain Macrophage Development. Cell 181, 557–573.e18 (2020).

86. Munro, D.A.D. et al. CNS macrophages differentially rely on an intronic Csf1r enhancer for their development. Development 147(2020).

87. Wu, X., Saito, T., Saido, T.C., Barron, A.M. & Ruedl, C. Microglia and CD206(+) border-associated mouse macrophages maintain their embryonic origin during Alzheimer’s disease. Elife 10(2021).

88. Masuda, T. et al. Specification of CNS macrophage subsets occurs postnatally in defined niches. Nature 604, 740–748 (2022).

89. Brioschi, S. et al. A Cre-deleter specific for embryo-derived brain macrophages reveals distinct features of microglia and border macrophages. Immunity 56, 1027–1045.e8 (2023).

90. Ren, X.S. et al. Hematopoietic Growth Factors Regulate the Entry of Monocytes into the Adult Brain via Chemokine Receptor CCR5. Int J Mol Sci 25(2024).

91. Bastos, J. et al. Monocytes can efficiently replace all brain macrophages and fetal liver monocytes can generate bona fide SALL1(+) microglia. Immunity 58, 1269–1288.e12 (2025).

92. Du, S. et al. Diversity and immune dynamics of choroid plexus macrophages are shaped by distinct developmental origins. Nat Neurosci 29, 717–731 (2026).

93. Du, S. et al. Brain-engrafted monocyte-derived macrophages from blood and skull-bone marrow exhibit distinct properties. Neuron (2026).

94. Ferrero, G. et al. Embryonic Microglia Derive from Primitive Macrophages and Are Replaced by cmyb-Dependent Definitive Microglia in Zebrafish. Cell Rep 24, 130–141 (2018).

95. Ferrero, G., Miserocchi, M., Di Ruggiero, E. & Wittamer, V. A csf1rb mutation uncouples two waves of microglia development in zebrafish. Development 148(2021).

96. Kizil, C., Iltzsche, A., Kaslin, J. & Brand, M. Micromanipulation of gene expression in the adult zebrafish brain using cerebroventricular microinjection of morpholino oligonucleotides. J Vis Exp, e50415 (2013).

97. Schonhoff, A.M. et al. Border-associated macrophages mediate the neuroinflammatory response in an alpha-synuclein model of Parkinson disease. Nat Commun 14, 3754 (2023).

98. Adler, D. et al. Functional border-associated macrophages limit Alzheimer’s Disease progression. bioRxiv (2026).

99. Shi, S.M. et al. Glycocalyx dysregulation impairs blood-brain barrier in ageing and disease. Nature 639, 985–994 (2025).

100. Levard, D. et al. Central nervous system-associated macrophages modulate the immune response following stroke in aged mice. Nat Neurosci 27, 1721–1733 (2024).

101. Navarro Negredo, P., et al. Targeting immune cells in the aged brain reveals that engineered cytokine IL-10 enhances neurogenesis and improves cognition. Immunity 59, 458–476.e13 (2026).

102. Mrdjen, D. et al. High-Dimensional Single-Cell Mapping of Central Nervous System Immune Cells Reveals Distinct Myeloid Subsets in Health, Aging, and Disease. Immunity 48, 380–395.e6 (2018).

103. Van Hove, H. et al. A single-cell atlas of mouse brain macrophages reveals unique transcriptional identities shaped by ontogeny and tissue environment. Nat Neurosci 22, 1021–1035 (2019).

104. Zelco, A. et al. Single-cell atlas reveals meningeal leukocyte heterogeneity in the developing mouse brain. Genes Dev 35, 1190–1207 (2021).

105. Kim, J.S. et al. A Binary Cre Transgenic Approach Dissects Microglia and CNS Border-Associated Macrophages. Immunity 54, 176–190.e7 (2021).

106. Karam, M., Janbon, H., Malkinson, G. & Brunet, I. Heterogeneity and developmental dynamics of LYVE-1 perivascular macrophages distribution in the mouse brain. J Cereb Blood Flow Metab 42, 1797–1812 (2022).

107. Ostkamp, P. et al. A single-cell analysis framework allows for characterization of CSF leukocytes and their tissue of origin in multiple sclerosis. Sci Transl Med 14, eadc9778 (2022).

108. Silvin, A. et al. Dual ontogeny of disease-associated microglia and disease inflammatory macrophages in aging and neurodegeneration. Immunity 55, 1448–1465.e6 (2022).

109. Benhar, I. et al. Temporal single-cell atlas of non-neuronal retinal cells reveals dynamic, coordinated multicellular responses to central nervous system injury. Nat Immunol 24, 700–713 (2023).

110. Robert, S.M. et al. The choroid plexus links innate immunity to CSF dysregulation in hydrocephalus. Cell 186, 764–785.e21 (2023).

111. Dissing-Olesen, L. et al. FEAST: A flow cytometry-based toolkit for interrogating microglial engulfment of synaptic and myelin proteins. Nat Commun 14, 6015 (2023).

112. Jeon, J. et al. Deciphering perivascular macrophages and microglia in the retinal ganglion cell layers. Front Cell Dev Biol 12, 1368021 (2024).

113. Wang, L. et al. CCR2(+) monocytes replenish border-associated macrophages in the diseased mouse brain. Cell Rep 43, 114120 (2024).

114. Jiang, X. et al. A single-cell atlas of mouse central nervous system immune cells reveals unique infection-stage immune signatures during the progression of meningitis caused by Streptococcus suis. Commun Biol 8, 1312 (2025).

115. Sun, L. et al. Deciphering the temporal transcriptional landscape of human fetal leptomeninges. Brain 148, 2218–2232 (2025).

116. Schweigreiter, R., Roots, B.I., Bandtlow, C.E. & Gould, R.M. Understanding myelination through studying its evolution. Int Rev Neurobiol 73, 219–73 (2006).

117. Baraban, M., Mensch, S. & Lyons, D.A. Adaptive myelination from fish to man. Brain Res 1641, 149–161 (2016).

118. McNamara, N.B. et al. Microglia regulate central nervous system myelin growth and integrity. Nature 613, 120–129 (2023).

119. Zupanc, G.K. Adult neurogenesis and neuronal regeneration in the brain of teleost fish. J Physiol Paris 102, 357–73 (2008).

120. Ribeiro Xavier, A.L., Kress, B.T., Goldman, S.A., Lacerda de Menezes, J.R. & Nedergaard, M. A Distinct Population of Microglia Supports Adult Neurogenesis in the Subventricular Zone. J Neurosci 35, 11848–61 (2015).

121. Ganz, J. & Brand, M. Adult Neurogenesis in Fish. Cold Spring Harb Perspect Biol 8(2016).

122. De Vlaminck, K. et al. Differential plasticity and fate of brain-resident and recruited macrophages during the onset and resolution of neuroinflammation. Immunity 55, 2085–2102.e9 (2022).

123. Rebejac, J. et al. Meningeal macrophages protect against viral neuroinfection. Immunity 55, 2103–2117.e10 (2022).

124. Kim, M.W. et al. Endogenous self-peptides guard immune privilege of the central nervous system. Nature 637, 176–183 (2025).

125. Faraco, G. et al. Perivascular macrophages mediate the neurovascular and cognitive dysfunction associated with hypertension. J Clin Invest 126, 4674–4689 (2016).

126. Park, L. et al. Brain Perivascular Macrophages Initiate the Neurovascular Dysfunction of Alzheimer Aβ Peptides. Circ Res 121, 258–269 (2017).

127. Pedragosa, J. et al. CNS-border associated macrophages respond to acute ischemic stroke attracting granulocytes and promoting vascular leakage. Acta Neuropathol Commun 6, 76 (2018).

128. Gericke, C., Mallone, A., Engelhardt, B., Nitsch, R.M. & Ferretti, M.T. Oligomeric Forms of Human Amyloid-Beta(1-42) Inhibit Antigen Presentation. Front Immunol 11, 1029 (2020).

129. Rajan, W.D. et al. Defining molecular identity and fates of CNS-border associated macrophages after ischemic stroke in rodents and humans. Neurobiol Dis 137, 104722 (2020).

130. Wan, H., Brathwaite, S., Ai, J., Hynynen, K. & Macdonald, R.L. Role of perivascular and meningeal macrophages in outcome following experimental subarachnoid hemorrhage. J Cereb Blood Flow Metab 41, 1842–1857 (2021).

131. Ochocka, N. et al. Single-cell RNA sequencing reveals functional heterogeneity of glioma-associated brain macrophages. Nat Commun 12, 1151 (2021).

132. Spiteri, A.G. et al. PLX5622 Reduces Disease Severity in Lethal CNS Infection by Off-Target Inhibition of Peripheral Inflammatory Monocyte Production. Front Immunol 13, 851556 (2022).

133. Warrington, J.P. et al. Pial Vessel-Associated Microglia/Macrophages Increase in Female Dahl-SS/Jr Rats Independent of Pregnancy History. Int J Mol Sci 23(2022).

134. Calero, M. et al. Lipid nanoparticles for antisense oligonucleotide gene interference into brain border-associated macrophages. Front Mol Biosci 9, 887678 (2022).

135. Ivan, D.C. et al. Insulin-like growth factor-1 receptor controls the function of CNS-resident macrophages and their contribution to neuroinflammation. Acta Neuropathol Commun 11, 35 (2023).

136. Mutoh, T., Kikuchi, H., Jitsuishi, T., Kitajo, K. & Yamaguchi, A. Spatiotemporal expression patterns of ZBP1 in the brain of mouse experimental stroke model. J Chem Neuroanat 134, 102362 (2023).

137. Kearns, N.A. et al. Dissecting the human leptomeninges at single-cell resolution. Nat Commun 14, 7036 (2023).

138. Uekawa, K. et al. Border-associated macrophages promote cerebral amyloid angiopathy and cognitive impairment through vascular oxidative stress. Mol Neurodegener 18, 73 (2023).

139. Colella, P. et al. CNS-wide repopulation by hematopoietic-derived microglia-like cells corrects progranulin deficiency in mice. Nat Commun 15, 5654 (2024).

140. Ganz, T. et al. Targeting CNS myeloid infiltrates provides neuroprotection in a progressive multiple sclerosis model. Brain Behav Immun 122, 497–509 (2024).

141. Glavan, M. et al. CNS-associated macrophages contribute to intracerebral aneurysm pathophysiology. Acta Neuropathol Commun 12, 43 (2024).

142. Santisteban, M.M. et al. Meningeal interleukin-17-producing T cells mediate cognitive impairment in a mouse model of salt-sensitive hypertension. Nat Neurosci 27, 63–77 (2024).

143. Xu, H. et al. The choroid plexus synergizes with immune cells during neuroinflammation. Cell 187, 4946–4963.e17 (2024).

144. Anfray, A. et al. A cell-autonomous role for border-associated macrophages in ApoE4 neurovascular dysfunction and susceptibility to white matter injury. Nat Neurosci 27, 2138–2151 (2024).

145. Yang, S. et al. Amantadine modulates novel macrophage phenotypes to enhance neural repair following spinal cord injury. J Transl Med 23, 60 (2025).

146. Xiong, X.Y. et al. Single-nucleus RNA sequencing reveals the specific molecular signatures of myeloid cells responding to brain injury after microglial replacement. Front Immunol 16, 1625673 (2025).

147. Okuno, N. et al. Brain macrophages and pial fibroblasts promote inflammation in a hypomyelination model. Acta Neuropathol Commun 13, 145 (2025).

148. Strehle, J. et al. The border-associated macrophage marker MRC1 contributes to an early neuroprotective inflammatory response to traumatic brain injury in mice. Acta Neuropathol Commun 13, 220 (2025).

149. Ohki, T. et al. Depletion of CD169(+) border-associated macrophages induces Parkinson’s disease-like behavior. Front Neurosci 19, 1688394 (2025).

150. Van Hove, H. et al. Interleukin-34-dependent perivascular macrophages promote vascular function in the brain. Immunity 58, 1289–1305.e8 (2025).

151. Wu, X., Miller, J.A., Lee, B.T.K., Wang, Y. & Ruedl, C. Reducing microglial lipid load enhances β amyloid phagocytosis in an Alzheimer’s disease mouse model. Sci Adv 11, eadq6038 (2025).

152. Dyckhoff-Shen, S. et al. Pharmacologic depletion of border-associated macrophages worsens disease in a mouse model of meningitis. Acta Neuropathol Commun 13, 191 (2025).

153. Hu, M. et al. Senescent-like border-associated macrophages regulate cognitive aging via migrasome-mediated induction of paracrine senescence in microglia. Nat Aging 5, 2039–2054 (2025).

154. Zhou, J. et al. Remote ischemic postconditioning improves cognitive dysfunction after subarachnoid hemorrhage by driving metabolic reprogramming of border-associated macrophages through the IL-33/ST2 axis. J Neuroinflammation 22, 257 (2025).

155. Baluszek, S. et al. Determining the Biological Features of Aggressive Meningioma Growth with Transcriptomic Profiling. Cancers (Basel) 17(2025).

156. Yu, N. et al. Changes in border-associated macrophages after stroke: Single-cell sequencing analysis. Neural Regen Res 21, 346–356 (2026).

157. Gao, Y. et al. Meningeal blood vessel blockage enhances anti-glioblastoma immunity. Cell 189, 1305–1322.e28 (2026).

158. Liu, N. et al. Early glymphatic failure in AppNL-F knock-in mice is linked to parenchymal border macrophages loss. Brain, awag080 (2026).

159. Guo, P. et al. Glibenclamide alleviates hydrocephalus after intraventricular hemorrhage by targeting metabolic reprogramming of border-associated macrophages. Exp Neurol 401, 115751 (2026).

160. Seferbekova, Z. et al. Spatial Transcriptomics Characterisation of Radionecrotic Changes in Glioblastoma Patients. Neuro Oncol (2026).

161. Tang, F. et al. BMAL1-GPX3 axis in the choroid plexus mitigates Aβ pathology in an amyloid mouse model. J Neuroinflammation 23(2026).

162. Garcia-Bonilla, M. et al. Post-hemorrhagic hydrocephalus of prematurity is associated with disruption of tight junctions and increased macrophage activity in the choroid plexus. Fluids Barriers CNS (2026).

163. Ratzabi, A. et al. Brain metastases exhibit distinct spatial patterns of resident and infiltrating macrophages. Cell Death Discov (2026).

164. Philippe, T.J., et al. A Multi-omic Atlas of Human Choroid Plexus in Alzheimer’s Disease. bioRxiv (2025).

165. Zawadzki, M.E., et al. Fluid-Niche and Microglial Signatures Prime Robust Intraventricular Macrophage Response to Blood During Brain Development. bioRxiv (2025).

166. Bhatta, S., Grzybowski, A., Ortiz, G. & DiStasio, M. Mechanobiological Specialization of Choroid Plexus Macrophages Defined by Titin Expression. bioRxiv (2026).

167. Kim, S., et al. Meningeal inflammation and arachnoid barrier breakdown in a mouse model of neonatal bacterial meningitis. bioRxiv (2026).

168. Colacurcio, D.J. & Nixon, R.A. Disorders of lysosomal acidification-The emerging role of v-ATPase in aging and neurodegenerative disease. Ageing Res Rev 32, 75–88 (2016).

169. D’Angelo, L. Brain atlas of an emerging teleostean model: Nothobranchius furzeri. Anat Rec (Hoboken) 296, 681–91 (2013).

170. Chi, J., Crane, A., Wu, Z. & Cohen, P. Adipo-Clear: A Tissue Clearing Method for Three-Dimensional Imaging of Adipose Tissue. J Vis Exp (2018).

171. Bankhead, P. et al. QuPath: Open source software for digital pathology image analysis. Sci Rep 7, 16878 (2017).

172. Bhattarai, P. et al. IL4/STAT6 Signaling Activates Neural Stem Cell Proliferation and Neurogenesis upon Amyloid-β42 Aggregation in Adult Zebrafish Brain. Cell Rep 17, 941–948 (2016).

173. Mariën, V., Arckens, L. & Van Houcke, J. Preparation of High-Viability Single-Cell Suspensions from African Turquoise Killifish Brain Tissue. Cold Spring Harb Protoc 2024, 107829 (2024).

174. Glasauer, S.M. & Neuhauss, S.C. Whole-genome duplication in teleost fishes and its evolutionary consequences. Mol Genet Genomics 289, 1045–60 (2014).

175. Singh, P.P. et al. Evolution of diapause in the African turquoise killifish by remodeling the ancient gene regulatory landscape. Cell 187, 3338–3356.e30 (2024).

176. Butler, A., Hoffman, P., Smibert, P., Papalexi, E. & Satija, R. Integrating single-cell transcriptomic data across different conditions, technologies, and species. Nat Biotechnol 36, 411–420 (2018).

177. Korsunsky, I., Nathan, A., Millard, N. & Raychaudhuri, S. Presto scales Wilcoxon and auROC analyses to millions of observations. bioRxiv, 653253 (2019).

